# Complementary multiphoton tools to create 3D architectures in soft hydrogels for epithelial tissue engineering

**DOI:** 10.64898/2026.03.31.715498

**Authors:** Simon Moser, Amelia Hasenauer, Xueting Shen, Shivaprakash N. Ramakrishna, Lucio Isa, Robert Style, Marcy Zenobi-Wong

## Abstract

Curvature provides essential mechanical cues for epithelial cells, playing a key role in cell differentiation and morphology. Repeatable manufacture of precisely controlled curvature in soft hydrogel materials is therefore essential to study epithelial mechanobiology and function. Multiphoton (MP) based biofabrication holds promise due to its high resolution and three-dimensional design flexibility. Here, we leverage MP’s advantages while increasing print speed to develop two complementary tools based on replica molding and multiphoton ablation. These can provide scalable hydrogel curvatures with tunable surface properties relevant for epithelial tissue engineering. In replica molding, MP prints are transferred into PDMS used to pattern centimeter scale arrays in hydrogels. In multiphoton ablation, hydrogels are locally degraded to generate precisely controlled curvatures and surface topography. With both methods, we repeatably guide epithelial cells into alveolar and duct-like shapes. Concave alveolar-like surfaces are shown to enhance the formation of thicker epithelial layers. We observe that surface properties, controlled by both tools, could enhance cytoskeletal organization. Using these biofabrication techniques, individual effects of curvature, surface properties, hydrogel composition, and bulk stiffness on epithelial cells can be studied. Both approaches offer high curvature control and throughput, providing a viable alternative to traditional 3D culture and other printing methods.

## 1. Introduction

Epithelial tissues are curved over multiple scales, and this curvature has been shown to influence cell behavior and function in vitro.^[1]^ In intestine cells, cylindrical curvature improves cell differentiation.^[2]^ In kidney cells, cylindrical curvatures introduce stresses, increasing epithelial layer thickness.^[3]^ Saddle shaped curvatures affect chromatin packing and gene accessibility.^[4]^ In mammary epithelial cells, curvature is hypothesized to affect expression of lactation markers.^[5]^ These findings suggest that curvature can play an important role in vivo and should be included when studying cells in vitro. However, these findings are based on substrates which fail to replicate the curvatures found in vivo.

Epithelial tissues such as pancreas, breast and lung have concave spherical shapes with radii between 25 and 125 µm.^[6-8]^ The intestine wall is characterized by ellipsoids which are concave in the crypts and convex in the villi.^[2]^ Current fabrication techniques for cell-culture substrates are limited in achieving such curvatures. The most common technique to create curved surfaces in vitro is soft lithography which can easily produce 2.5D structures, ^[2,9-11]^ but has limited capabilities in 3D.^[9,12,13]^ Alternatives, such as DLP printing fail to deliver the resolution required for epithelial-relevant curvatures.^[4,14,15]^ Organoids on the other hand have an inherently variable shape and are closed systems with limited lifespan, due to waste accumulation in the lumen, something which also hampers analysis of secreted components important in many epithelial tissues.^[16]^ Additionally, the enclosed lumen makes it hard to investigate the effect of luminal composition and diseases on the integrity and function of epithelial layers.^[17,18]^

These limitations highlight the need for versatile techniques that can produce a range of physiologically-relevant, complex-shaped surfaces to enable mechanistic studies on the effects of curvature on cell behavior, and to determine which curvature is needed for tissue function. Here we demonstrate two complementary approaches to achieve complex curvatures, taking advantage of the micron level resolution of multiphoton printing. In the first approach, we adapt a replica molding approach introduced by Callens et al.^[19]^ to soft hydrogels relevant for epithelial tissue engineering (e.g. 0.7 to 6 kPa in breast tissue ^[15,20]^), starting from two-photon printed molds. We refer to this method as 2-photon polymerization and replica molding (2PP+RM). In the second approach, we use multiphoton ablation (MPA) to locally degrade hydrogels using a focused laser beam.

While recent advances in commercial 2PP printers have enabled increased scanning speed.^[21,22]^, printing is still slow and mostly limited to small structures. In 2PP+RM, repeatable use of molds enhances scalability, and removes the need for the end user to have access to expensive 2PP printing equipment. In MPA, we reduce printing time by drastically increasing the distance between scanned lines while maintaining micrometer resolution.

Importantly, both techniques allow us to tailor surface properties of the hydrogel in a manner which greatly enhances cell attachment. In 2PP+RM, we can tune the hydrogel molding step to produce surfaces that are significantly softer than the bulk hydrogel. This results from the hydrogel precursor not fully polymerizing against a PDMS surface.^[23,24]^ In MPA, we use its high local control to partially degrade the hydrogel creating softer surfaces or controlled roughness.

## 2. Results

Figure 1A shows the principle behind multiphoton absorption leading to its high resolution in contrast to single photon absorption. The multiphoton process relies on the simultaneous absorption of multiple photons at the same time and place. This requires a high irradiance corresponding to a high photon density, making it much more likely that multiple photons meet. This condition is only met in a tiny focal spot, whereas single photon absorption occurs within a much larger volume. The small focal spot where excitation occurs leads to confined initiation and high-resolution printing (Figure 1A, right).^[25,26]^ In order to prevent ‘boiling’ the sample, the energy needs to be supplied in short, femtosecond pulses.^[25]^ The multiphoton process can be both adapted to additive manufacturing by polymerization as well as subtractive degradation of hydrogels.

**Figure 1:**
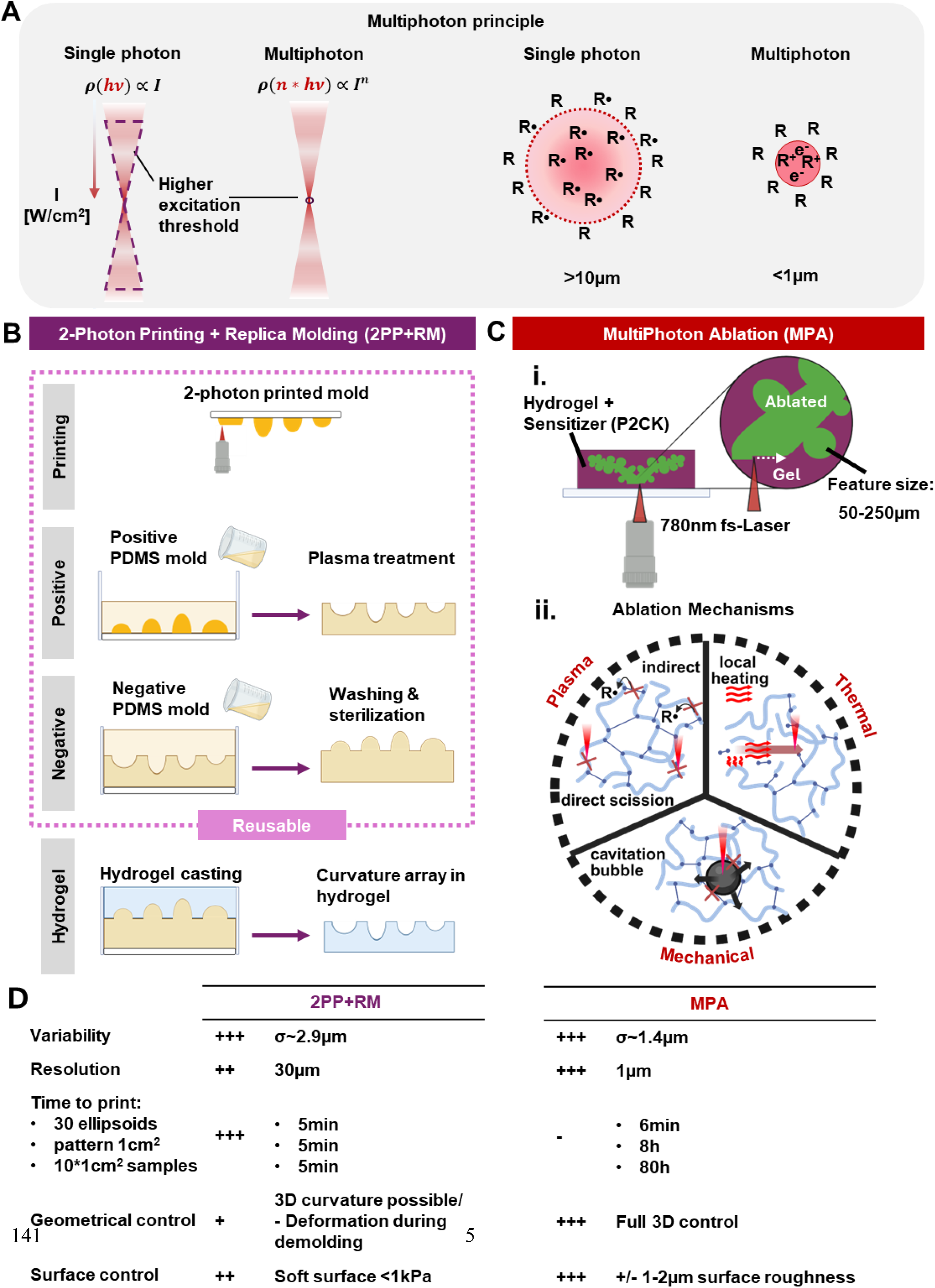
Multiphoton based biofabrication of precise curvature arrays: **[A]** Multiphoton printing relies on the near simultaneous adsorption of multiple photons leading to localized initiation. Single photon adsorption is proportional to the irradiance (I) resulting in a large excitation threshold whereas multiphoton adsorption is proportional to In, where n is the number of photons. This lower excitation threshold leads to a smaller and more confined initiation. R• and R+ are radicalized or ionized molecules respectively which are the starting point for polymerization or ablation. **[B]** Adaptation of traditional PDMS soft lithography process using a high resolution 2PP printed mold using commercial resins with higher flexibility over achievable curvatures. **[C] i**. Multiphoton ablation with an IR femtosecond(fs) laser adapted to larger duct and alveoli like features relevant to epithelial tissue engineering in hydrogels using 2PP sensitizer P2CK. **ii**. Main mechanisms driving multiphoton ablation: Plasma-induced indirect scission of polymer chains, thermal heating and mechanical forces resulting from cavitation bubble formation. **[D**] Side by side overview of advantages/disadvantages of 2PP+RM and MPA. Time to print is referring to: thirty 200 µm half spheres, 1cm2 fully packed with half spheres of same size and 10 samples of this

Figure 1B shows the 2PP+RM method based on additive two photon printing. Here we take advantage of the high resolution of two-photon printing while reducing printing time per curvature array by replicating into polydimethylsiloxane (PDMS). PDMS provides great flexibility and reusability of the molds.

In this method, we first manufacture a positive mold using a commercial 2PP printer. To get to the final desired concave curvature we replicate twice into PDMS with in-between air plasma treatment to allow mold release. The final negative mold is then washed and sterilized and used for hydrogel casting. Alternatively, we print a negative mold first which requires more volume to print but it allows us to skip one molding step (Figure S1, S2, Supporting Information). This is the favored approach if access to printing time is not a limiting factor.

Figure 1C shows the subtractive MPA process where the hydrogel is locally degraded to manufacture complex duct and alveoli like geometries. Since only the smaller cavities and not the whole bulk hydrogel has to be printed this is faster compared to additive printing. However, this process is still slow, requiring hours to print relevant structures (Figure S9, Supporting Information).

Thus, we soak the hydrogels with P2CK, a sensitizer previously shown to reduce the required laser power^[27]^ and energy,^[28]^ to increase multiphoton absorption which in turn increases printing speed and broadens the parameter space in which ablation occurs (further referred to as ablation window) (Figure S9-S12, Supporting Information). The increased absorption increases plasma formation and heat generation beneficial for multiphoton ablation (Figure 1C ii). However, it also increases cavitation bubble formation leading to mechanical deformation. We avoid cavitation by increasing printing speed avoiding energy build up that would lead to bubble formation.

Both 2PP+RM and MPA provide high spatial resolution, low variability and control over surface properties (Figure 1D). For applications where high throughput screening is needed, the replica molding approach allows us to produce multiple samples in just 5 minutes. When full geometrical control is required, MPA is the method of choice.

As we shall see below, both 2PP+RM and MPA allow one to introduce a soft surface layer into the constructs. With 2PP+RM, the soft surface layer (<1 kPa) is produced during demolding and combined with the underlying stiffer substate allows for excellent handling. Similarly, MPA allows local and highly resolved texturing/softening of the ablated surface. In both MPA and 2PP+RM we can tune the bulk mechanical properties independent of the printing process.

### 2.1 2PP-based Replica Molding of soft hydrogels retains high repeatability with favorable surface properties

2PP+RM involves three key steps. The high precision of 2PP printing, the first step, has been well established. However, it is unknown how well geometries are replicated into PDMS and hydrogels subsequently. Thus, we investigate how precise shape control is retained over each step.

First, we looked at the shape fidelity and repeatability of the final PDMS molds. To mimic epithelial tissues,^[2,6-8]^ we designed concave half ellipsoids with varying radii ranging from 50 to 250µm and transferred them into PDMS (Figure 2A). Average radii of the PDMS half ellipsoids were measured showing high repeatability and shape fidelity (Figures 2B, S1, Supporting Information).

**Figure 2:**
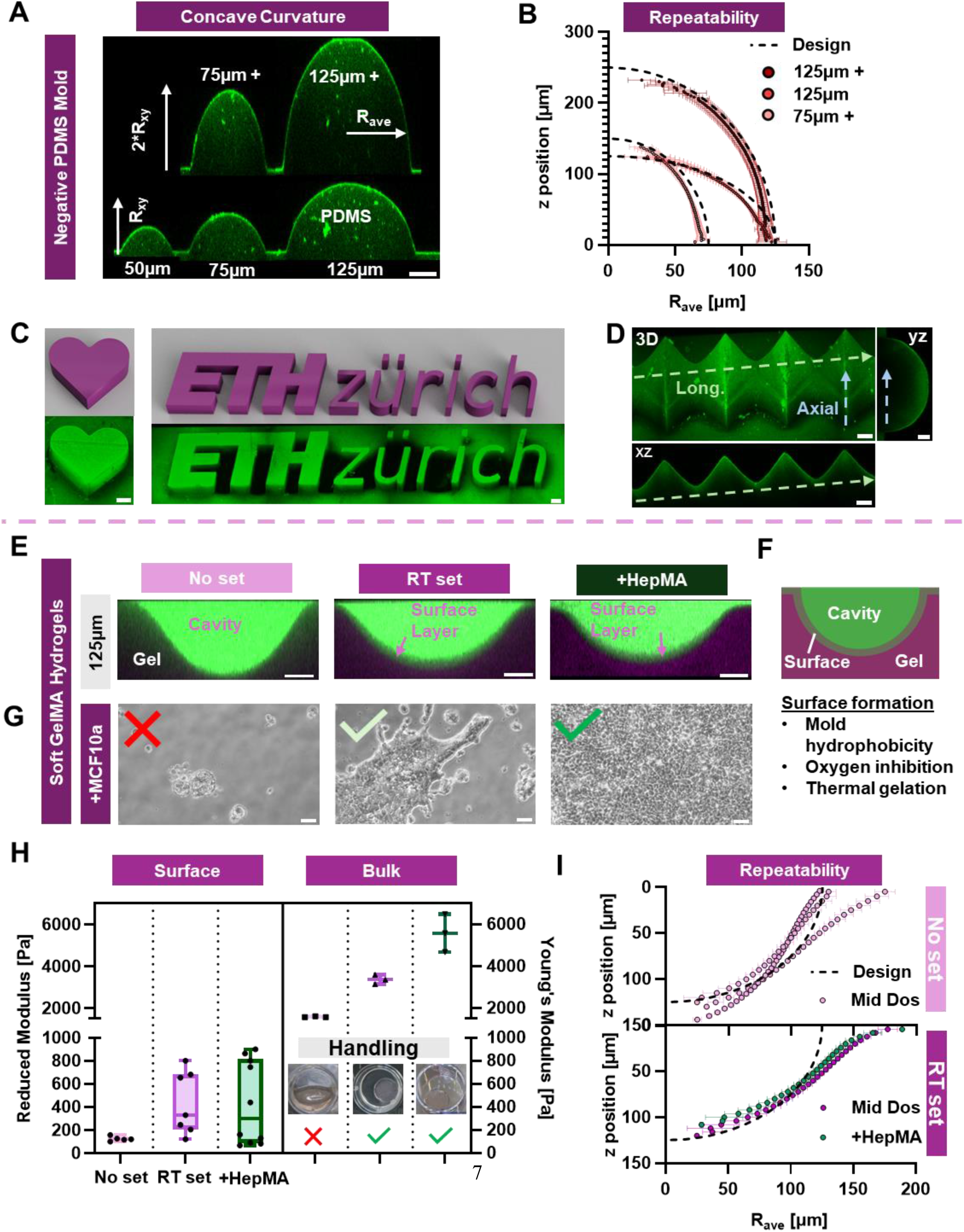
Characterization of final two replica molding steps and the usability of hydrogels for mammary epithelial cell culture: **(A)** Top row: XZ cross-section of negative PDMS mold Ellipsoid depth=2*Rxy (150, 250 µm), obtained by confocal microscopy. Bottom row: half spheres with varying radius of curvature in plane (100; 150; 250 µm). PDMS was stained with Rhod B. (B) Average in plane radius of 250 µm half sphere and 250 and 150 µm ellipsoids are given for every z position, wells (w)=2; samples (n)=3. (C) Heart and ETH Zürich logo designs (pink) and final PDMS mold (green), scale bar 50um. (D) PDMS mold of with convex axial curvature (concave in hydrogel) and alternating concave and convex curvature longitudinally. The mold was directly replicated from a 2PP negative mold. (E) Cross sectional confocal images of 250 µm half sphere transferred into 5 wt% GelMA based hydrogels exposed to different pre-crosslinking treatments. From left to right: direct UV crosslinking (no set), 1h RT treatment (RT set), 1h RT treatment + 1 wt% HepMA (+HepMA), arrow indicates formed surface layer. Gels were crosslinked with Rhod B acrylate (magenta) and cavity was perfused with 2 MDa FITC Dextran (green). (F) Sketch of surface layer formation using a combined effect of mold hydrophobicity, oxygen inhibition and thermal pre-crosslinking. (G) Bright field images of MCF10a cell attachment after 4 days of culture on no set, RT set and +HepMA hydrogels n>4 (H) Reduced surface modulus measured by AFM indentation (left) and bulk Young’s modulus (right) measured by compression tests for a no set, RT set, +HepMA gels. Inset shows compactness of hydrogel during cell culture. (I) Average radius (Rave) obtained from confocal z stacks as a function of z position for the three conditions. All three replicates are shown for the no set hydrogels. w=2 n=3. Scale bars are 50 µm if not indicated otherwise.

2PP+RM is not limited to concave half ellipsoidal shapes. Provided the design allows mold release, any arbitrary shape can be replicated, even challenging small or complex features such as a heart shape (Figure 2C, S4B-D Supporting Information) or the ETH Zürich Logo (Figures 2C, S4A Supporting Information). This is important since it enables us to produce molds with more complex, physiologically relevant geometries including both concave and convex principal curvatures (Figures 2D, S3, Supporting Information). 2PP+RM is freely scalable, showcasing the combination of millimeter-sized features with micrometer-scale resolution.

As a last step, we molded GelMA-based resins against PDMS to create a more representative environment for the cells (Figure 2E). GelMA allows great tunability of mechanical properties by thermal crosslinking and subsequent UV polymerization,^[29]^ while containing cell-adhesion motifs, making it a suitable substrate. We successfully replicated 250 µm half spherical geometries containing concave curvatures. Molding against hydrophobic and oxygen permeable PDMS in combination with an oxygen sensitive resin creates a weakly crosslinked surface layer (Figure 2E, 2F, S6A, Supporting Information).^[23,24]^ The propagating methacryloyl radicals of GelMA react with dissolved oxygen inhibiting polymerization at the surface.^[30]^ Thermal pre-gelation further strengthens the surface creating a clearly defined layer (Figure S6-8 Supplementary Information). Moreover, it reduces slipperiness in handling observed for direct UV crosslinked gels.^[23,24]^

In addition to thermal crosslinking and mold material we observe that a reduction in power density (Figure S6D Supporting Information) and UV curing time (Figure S8 Supporting resin (Figure S8C-H Supporting Information). Despite the thick surface layer of some conditions, concave curvatures were faithfully replicated into such hydrogels and supported cell attachment (Figure S6, Supporting Information).

Next, we assessed whether the introduced differences in surface structure had any effect on cell attachment. We used the mammary epithelial cell line MCF10a as a model system. On medium degree substitution GelMA (Mid DoS, Table S1 Supporting Information) on day 4, cells showed a rounded morphology indicating a reduced ability to interact with the substrate (Figure 2G, left). In contrast, pre-gelation at room temperature (RT) of GelMA did allow cell attachment but no confluent layer formed yet. Adding HepMA (a modified component of the natural ECM)^[6, 31]^ cells reached confluence by day 4 and displayed a typical cobble stone morphology (Figure 2G, right).

The influence of surface structure on cell attachment was further confirmed by direct UV crosslinking of GelMA with different degree of substitution against substrates of varying oxygen permeability and hydrophobicity (PDMS, polycarbonate (PC) and glass, Figure S5, Supporting Information), a strategy known to influence surface properties.^[23,24]^ For Mid DoS GelMA cell attachment improved slightly when shifting from PDMS to PC and improved more when using a glass mold. For High DoS GelMA cell confluency improved again when switching to PC but when molding against glass cells elongated, a sign of higher stiffness.^[32]^ Cell attachment on directly UV crosslinked gels showed that neither thermal crosslinking nor hydrophobicity was necessary for cell attachment (Figure S7, Supporting Information). Despite possibilities to print glass molds with 2PP,^[33]^ we refrained from using it due to introduction of surface defects.

Consistent with confocal imaging we found that the surface stiffness was softer compared to bulk stiffness for the highlighted conditions (Figure 2H). Surface stiffness was below 1 kPA, which is difficult to achieve with other methods. This was also the case for directly UV crosslinked gels which shows that stiffness is not the sole factor determining cell attachment. For RT set and + HepMA gels, higher values in surface stiffness and a higher variability were observed in agreement with the lower surface thickness measured for those conditions (Figure S8, Supporting Information). Constructs with bulk stiffness below 2 kPa exhibited reduced integrity and were challenging to handle, unlike gels with higher stiffness (Figure 2H, Inset).

Lastly, we wanted to confirm whether the high precision of 2PP was transferred into the final hydrogel curvatures. Thermal gelation led to an observable reduction of sample-to-sample variation (Figure 2I). Overall, we observed a widening towards the top of the hydrogel features for thermally crosslinked gels, whereas non set gels showed this effect in only one sample. This deformation is introduced during demolding as it is not observed in the PDMS mold. Hydrogel shape variability was low despite the soft stiffness (Figure 2I), with inter-sample and intra-sample standard deviations of 5.08 and 2.12 µm, respectively, for +HepMA gels. Since the deviation from the original design is repeatable and systematic, future iterations on the model could compensate for this effect.

### 2.2 Scaling up of Multiphoton ablation (MPA) to fabricate more complex alveolar and ductal like geometries

In contrast to RM+2PP, Multiphoton Ablation (MPA) allows us to create more complex three-dimensional structures as seen in Figure 1C. However, this technique has so far been limited to µm-scale structures not suitable for epithelial tissue engineering. To print larger structures in a reasonable time, we systematically explored the parameters shown in Figure 3A.

**Figure 3:**
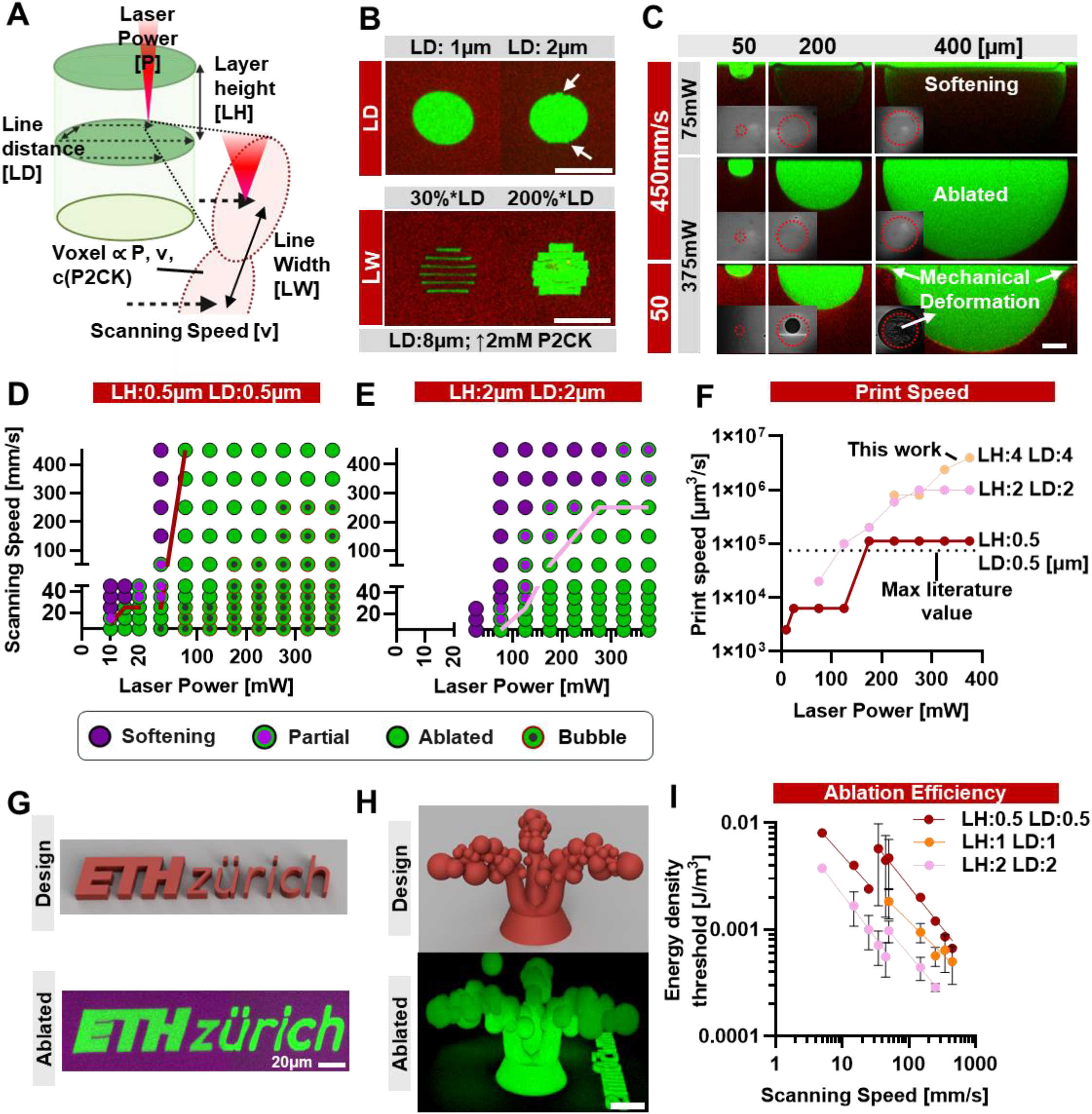
Optimization of Multiphoton ablation for larger and more complex geometries: **(A)** Sketch of the most important laser parameters varied in this study to enable printing of features >50 µm, Voxel size in plane is artificially stretched by changing the line width (LW). Voxel size further depends on Laser power (P), Scanning Speed (v) and concentration of the sensitizer (P2CK). **(B)** Effect of increased line distance on surface quality of a 50 µm cylinder (top row, 375 mW 50 mm s^-1^) Effect of linewidth shown for a line distance of 8µm bottom row, (2 mM P2CK LH:2 µm). RT set GelMA (Mid DoS) **(C)** Ablation of 50, 200 and 400µm half spheres with varying Laser power and scanning speeds. Cavitation bubble formation is shown in inset, mechanical deformation resulting from expanding bubbles are highlighted with white arrows. LH:0.5 µm LD: 0.5 µm 1 mM P2CK for all conditions. **(D)** High surface quality dose test (LH:0.5 µm LD: 0.5 µm) of RT set Mid DoS GelMA + 1mM P2CK. An array of 50 um cylinders was designed, varying laser power between 5-25mW and 25-375 mW and Scanning speeds from 5-45mm s^-1^ and 50-450mm s^-1^. Cylinders where median (n=3) is ablated are shown in green, cavitation bubble formation is indicated in black and incomplete or softened cylinders are indicated in magenta. **(E)** Higher throughput dose test (LH:2 µm LD: 2 µm) **(F)** Theoretical print speed for varying hatching distances showing a orders of magnitude higher speed compared to previously reported values.^[22]^ Median is shown **(G)** Design and ablated ETH Logo imaged with confocal microscopy **(H)** Design and Ablated version of alveolar + duct like structure (1mM P2CK 0.5 µm 0.5 µm 375mW 50mm s^-1^). All cavities are stained with 2M Da FITC dextran (green), gels crosslinked with Rhod B acrylate (red). All data was analyzed from confocal images. **(I)** Energy density threshold needed for ablation for hatching distances 0.5,1 and 2 µm as a function of scanning speed. Non-linear log-log fit was applied to show trend. For all subfigures n=3. Scale bars are 50 µm if not indicated otherwise.

In addition to scanning speed and Laser power, two relevant parameters are layer height (LH): the distance between planes, and line distance (LD): the distance between scanned lines (referred together as hatching distance). Additionally, we can optically stretch the Laser focal spot (Voxel) orthogonally to the scanning direction, a parameter called line width (LW). Since LW can reduce the gap between lines, increasing LD in combination with LW increases print speed similarly to LH but has a lower effect on the ablation window (Figure S13, Supporting Information). The ideal parameter set depends on the sensitizer concentration (P2CK) with an increase enabling higher hatching distances and scanning speeds (Figure S13-S16, Supporting Information). Together, optimization of all these parameters allows for a reduction in the total distance the laser has to travel, thereby reduce the total printing time.

While a higher hatching distance gives the potential to increase print speed, it can also introduce surface roughness (Figure 3B). A hatching distance of 1 µm gives a smooth surface whereas 2 µm already introduces a slight roughness as seen in the xy plane (Figure 3B, top) but also in z (Figure S13, Supporting Information). A further increase in LD to 8 µm led to a considerable roughness and reduction circularity. However, this effect is smaller for larger structures relevant for epithelial tissue engineering (Figure S16E, Supporting Information). Such an increase in LD is only possible due to an increase in LW and P2CK concentration to ensure that adjacent voxels overlap (Figure 3B, bottom).

While larger structures are affected less by the introduced surface roughness, their ablation window decreases, which is problematic for epithelial tissue engineering (Figure 3C, S13J, Supporting Information). Increasing the diameter of a half sphere from 50 µm to 400 µm subsequently increases the power needed to ablate while at lower speeds onset of bubble formation is earlier for larger geometries (Figure 3C inset). Small bubbles cause no issues. However, when they expand, they mechanically deform the geometry, narrowing down the ablation window.

We saw the opposite, an increase in ablation window when we increased the field of view (FOV), the area the Laser can scan, shifting the ablation threshold towards lower power (Figure S18, Supporting Information). In this work, we split designs into equally sized FOVs to avoid local differences in the ablation window. We were able to increase scanning speed to 450 mm s^-1^ compared to previous studies by a factor of 1.5,^[22]^ with this higher scanning speed working better for ablation. However, the same low hatching distance was used making fabrication slow.

To further increase the print speed, we increased the hatching distance from 0.5 µm (Figure 3D) to 2 µm (Figure 3E). This decreased the ablation window but positively avoided bubble formation. A further increase of hatching distance to 4 µm required an increase of sensitizer concentration from 1 mM to 2 mM (Figures S13, S15, Supporting Information). With this, we achieve an increase in print speed (see S9, Supporting Information for calculation) by two orders of magnitude (Figure 3F). This is a theoretical value used to compare different parameters and materials. Actual print times deviate especially for smaller structures due to acceleration and deceleration of the laser (Figure S9, Supporting Information). Nevertheless, we were able to print five 200 µm half spheres with tunable surface roughness in 1 minute (Figure S17, Supporting Information).

Even when the hatching distance was increased, we saw no negative effect on the resolution (Figure S19, Supporting Information). Adding P2CK slightly decreased the achievable resolution from sub-micron to 1 µm. In pure hydrogels, higher order multiphoton adsorption of water is more dominant leading to a higher resolution^[26]^ whereas adding P2CK presumably shifts it to a 2-photon adsorption regime. We further showcased the resolution by ablating multiscale features in the example of an ETH Zürich logo with smallest feature sizes down to 5 µm (Figure 3G). Printing of feature sizes down to 1 µm is possible as well but the prints start to distort due to the softness of the hydrogel (Figure S19, Supporting Information).

Next, we applied the previous learnings to ablate a more complex alveolar and ductlike design that incorporate curvatures representative of human breast tissue (Figure 3H, top). We increased hatching distance (see Table S2, Supporting Information for detailed ablation parameters used in this work) and adapted both laser power and scanning speed to account for the larger design. When using parameters that lead to successful ablation in the dose test (200 mW, 200 mm s^-1^), the alveolar and duct model were only softened (Figure S20, Supporting Information). Increasing the power and speed to 375 mW and 450 mm s^-1^ led to ablation in the duct and alveoli-like shape on top but not in the larger base. Therefore, we decreased scanning speed for the base individually, saving printing time. We can further change the size of the design to investigate its effect on epithelial morphology. (Figure S20, Supplementary Information).

Lastly, we calculated how much energy density (j) was required to achieve ablation for different hatching distances using equation [ 1 ] with P being the laser power, *v*_*max*_ the maximum scanning speed, LH the layer height and LD the line distance:

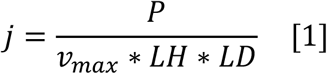

We saw with increasing print speed, both in terms of scanning speed but also by increasing the hatching distance we were able to reduce the required energy to ablate (Figure 3I). By increasing *v*_*max*_, LH and LD we decrease voxel overlap, avoiding excessive energy that goes into heating and eventually cavitation bubble formation. These findings complement our findings that multiphoton ablation is more efficient at higher printing speeds (Figure 3C, D).

### 2.3 Expansion of 2PP+RM and MPA techniques to biomaterials relevant to epithelial biology

Having shown previously that half spherical structures with a diameter of 250 µm can be reproducibly replicated with 2PP+RM (Figure 2H), we next assessed whether other curvatures relevant to epithelial tissue engineering could be reproduced. Figure 4A (left column) shows that curvatures with radii between 75 and 250 µm can be reproducibly manufactured using Mid DoS GelMA supplemented with HepMA, the substrate on which cells showed the best attachment, proliferation and morphology. Similar to the 250 µm half spherical curvature, we observed repeatable deformations and the formation of a soft surface layer.

**Figure 4:**
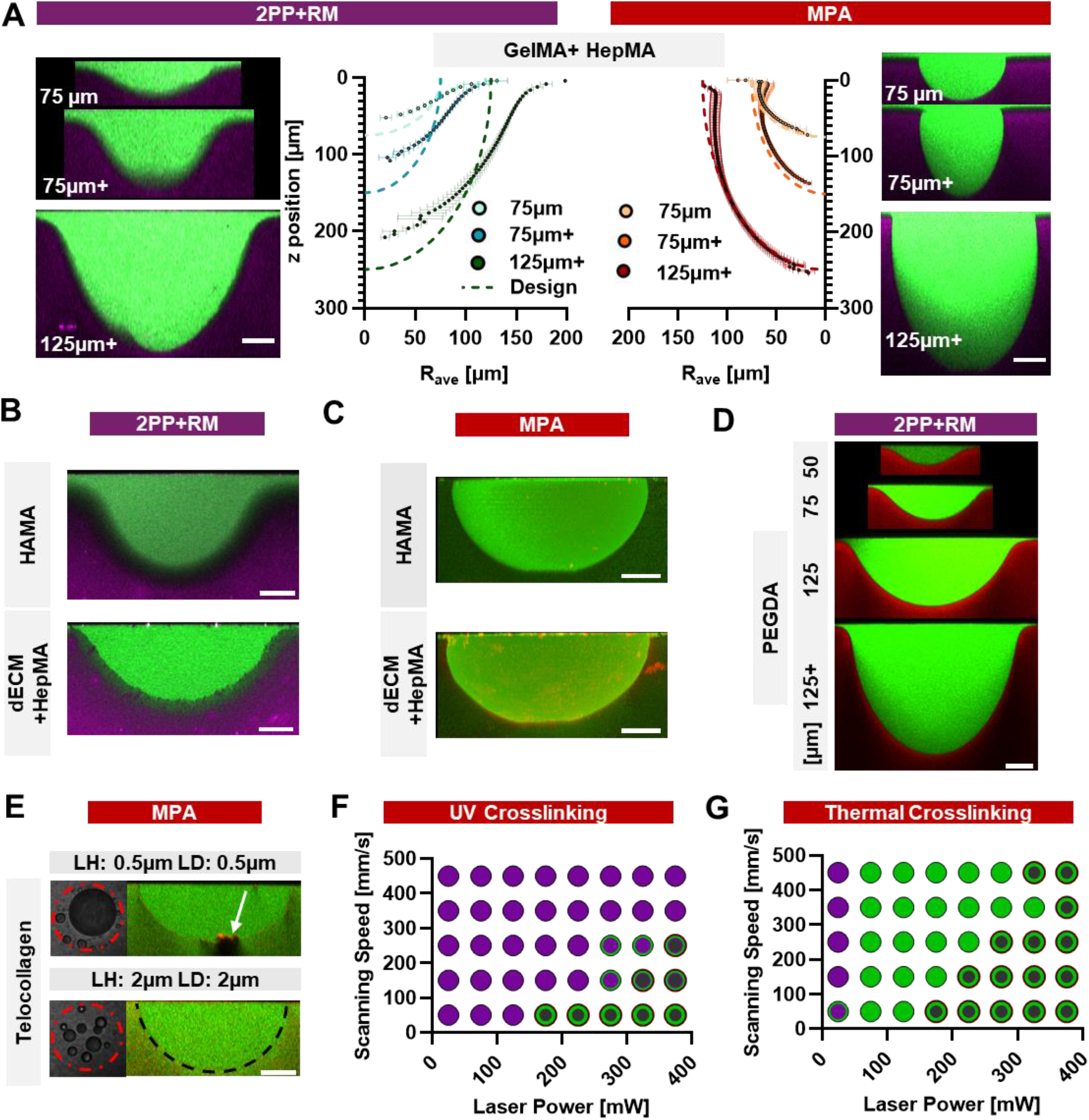
Comparison of repeatability of RM+2PP and MPA techniques and its applicability to different materials systems. **(A)** Curved features in Mid DoS GelMA + HepMA hydrogels fabricated with RM+2PP (left section) and MPA (right section). Average radius was calculated from each slice of a z-stack and ploted against the z-position. n=3; w=2 (B) Adaptation of 2PP+RM to non GelMA-based materials such as HAMA (top) or dECM + HepMA (bottom) showcased on the example of a 250 µm half sphere (C) Same material systems used for MPA (D) Various Ellipsoidal shapes transferred into 5 wt% PEGDA 700 hydrogels (E) Ablation in a soft 3 mg ml^-1^ thermally crosslinked Telocollagen gel using low hatching distances (top row) and high hatching distances (bottom row), bubble formation is shown on the left, confocal cross-section on the right. Mechanical deformation for low hatching distance is indicated with a white arrow (F) Dose test for High DoS GelMA which is mainly UV crosslinked and (G) low DoS GelMA which is thermally crosslinked at 4ºC before UV crosslinking, (LH 0.5 µm, LD: 0.5 µm). All images and analysis are based on confocal microscopy with cavities being perfused with 2M Da FITC Dextran and gels stained with Rhod B acrylate. Median is shown, n=3. All scale bars are 50 µm if not indicated otherwise.surface layer formed, indicating that this is a more general phenomenon with PDMS based replica molding.

Next, we used MPA to reproduce the same curvatures within the same material. To accommodate the increased stiffness of GelMA + HepMA gels, we first adapted the ablation settings by increasing the P2CK concentration (Figure S21, Supporting Information). This expanded the ablation window making it easier to optimize for larger and more complex structures. It also shifted the onset of bubble formation mostly determined by hatching distance and P2CK concentration rather than material composition (Figure S22, Supporting Information). In contrast the ablation border is influenced by material composition and stiffness due to additionally introduced bonds that need to be broken.

In terms of reproducibility, both 2PP+RM (Figure 4A, left) and MPA (Figure 4A, right) show low variance whereas MPA demonstrates higher shape fidelity. Additionally, MPA allowed us to ablate half spheres down to 50 µm (Figure S21, Supporting Information). To compare the repeatability between the two approaches we calculated the distribution of standard deviations of the average radii. We found the standard deviation to be between 1.36 µm (50 µm, MPA) and 5.85 (250 µm +, 2PP+RM). No significant differences were observed between the techniques, except for 250 µm + ellipsoids (Figure S23, Supporting Information). For both methods, variation was predominantly due to differences between samples rather than variation between wells within a sample.

Next, we used 2PP+RM and MPA to shape materials based on proteins and glycosaminoglycans (GAGs) found in native extracellular matrixes (ECM). The variety of material composition is large in tissue engineering and requirements vary depending on application, cell type and preference of the researchers.^[34]^ Therefore, we produced concave curvatures in some commonly used materials. First, we tested methacrylated hyaluronic acid (HAMA), its unmodified form being present in the extracellular matrix of many tissues including cartilage and skin.^[35]^ We successfully manufactured 250 µm half spheres with both 2PP+RM (Figure 4B, top row) and MPA (Figure 4C, top row). As observed with GelMA, a We further tested decellularized ECM (dECM)-based hydrogels, which are of broad interest for tissue engineering applications because they closely resemble the *in vivo* environment. Again, a surface layer was observed when dECM, extracted from bovine udders according to a previously published protocol,^[15]^ was crosslinked using a Ruthenium/SPS system, demonstrating that surface layer formation is not limited to a particular photoinitiator or material. HepMA was added to increase stiffness and improve ablation performance in addition to its affinity to growth factors (Figure S24, Supporting Information).^[31]^

In contrast to previous opinion,^[25]^ we found that MPA performs particularly well in hydrogels containing GAGs compared to purely protein-based hydrogels. Ablation in protein-based hydrogels, such as Collagen I (a main component of mammary ECM)^[15]^ works, but adsorption of the fibrils in the IR spectrum^[36]^ complicates parameter optimization. Nevertheless, we successfully ablated complex alveolar and duct like geometries in dECM hydrogels containing either HAMA or HepMA (Figure S25, Supporting Information).

Stiff, chemically crosslinked hydrogels are excellent for 2PP+RM (Figure 4D) but less compatible with MPA (Figure S25B, Supporting Information). PEGDA is a commonly used hydrogel in tissue engineering, often combined with cell attachment motifs such as RGD.^[37]^ We successfully replicated geometries relevant to epithelial tissue engineering into PEGDA. Interestingly, no surface layer formation was observed in confocal imaging, which is surprising since acrylates are reported to be more oxygen sensitive.^[38]^ This observation may be explained by our previous findings that the surface layer formation is not solely determined by oxygen inhibition but also by additional factors such as hydrophilicity differences of the resin/gel system, consistent with reports in the literature.^[23,24]^ In contrast, ablation in PEGDA did not work well, which we attribute to its high stiffness and the presence of non-cleavable bonds. (Figure S25B, Supporting Information).

On the contrary, we were able to generate concave curvatures in soft and viscoelastic hydrogels based on low concentration telocollagen using MPA (Figure 4E) but not 2PP+RM (Figure S24, Supporting Information). Upon demolding the thermally crosslinked telocollagen gels (3 mg ml^-1^) underwent substantial deformation (Figure S25 E, Supporting Information). In contrast, with MPA we successfully ablated 250 µm half spheres (Figure 4E). As observed for GelMA, printing quality was improved by increasing print speed with an increased hatching distance. Although telocollagen supports robust cell attachment highlighting the relevance of soft hydrogels (Figure S28, Supporting Information), handling such soft hydrogels is considerably more challenging compared to GelMA-based gels featuring the soft surface layer established in this work.

Thermal crosslinking is beneficial for MPA ablation and enable structuring of stiffer samples (Figure S26, Supporting Information). In contrast, chemically crosslinked gels (by UV exposure) based on High DoS GelMA showed poor adaptivity to MPA (Figure 4F), with bubble formation even preceding complete ablation, unlike stiffness-matched Mid DoS+HepMA gels (Figure S26, Supporting Information). Thermally crosslinked gels (Low DoS GelMA precured at 4ºC) on the other hand showed a large ablation window (Figure 4G). Notably, the degree of thermal crosslinking, varied here by the gelation temperature, did not influence the ablation window at low hatching distances (Figure S26, Supporting Information). However, at larger hatching distances it had a considerable effect reducing the ablation window relative to the softer hydrogels suggesting the presence of a threshold for complete local degradation.

### 2.4 Influence of the soft surface layer introduced by 2PP+RM on cell confluency and epithelial layer organization

Having established two tools, 2PP+RM and MPA, that allow us to shape a multitude of materials relevant for epithelial tissue engineering we moved on to investigate the impact of the surface properties controlled with both methods. First, we investigated the influence of the soft surface layer introduced by 2PP+RM on cell attachment. Consistent with observations in directly UV crosslinked GelMA (Figure S5, Supporting Information), we see substantial differences in cell confluency at day 1 on thermally pre-crosslinked gels molded against PDMS compared to those molded against PC (Figure 5A, top). The influence of the soft surface layer introduced by molding against PDMS and not PC (as shown in Figure S6A, Supporting Information) is reflected by the elongated morphology of MCF10a cells on PC molded gels due to an increase in effective stiffness. Moreover, molding against PDMS improved mold release avoiding surface defects. Mold release was further improved by addition of HepMA which additionally increased initial cell confluency compared to the stiffness matched GelMA hydrogels for both molding materials (Figure 5A, bottom). An increase in confluency was also observed in softer GelMA hydrogels (Figure S27, Supporting Information). While we saw these initial differences in cell attachment and proliferation, all conditions grew to 100% confluency (Figure S27, Supporting Information).

**Figure 5:**
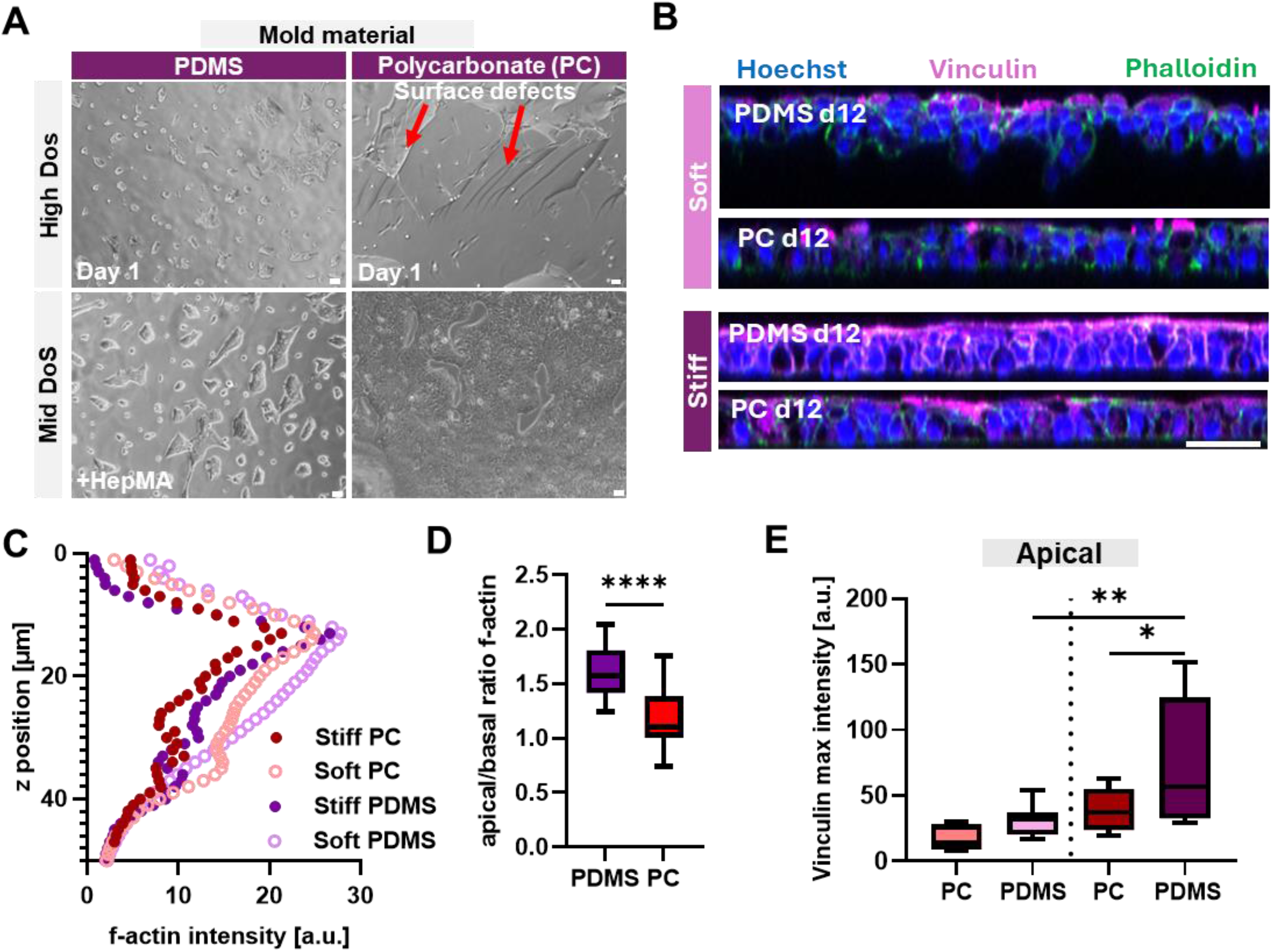
Influence of the soft surface layer on cell confluency and epithelial layer organisation. (A) BF images of MCF10a cell attachment on three different GelMA based material systems molded against PDMS (left row) or Polycarbonate (Petridish, right row) after 1 day. n=4 (B) Hoechst (blue) Phalloidin (green) and vinculin (magenta) expression of MCF10a cell layer after 12 days of culture on soft hydrogels (Mid Dos GelMA) and stiff hydrogels (+HepMA) molded against PDMS (top) and PC (bottom). Red arrows indicate infiltration of MCF10a cells into the underlying substrate. w=2; n=4 (C) Average F-actin intensity obtained by phalloidin staining as a function of z-position for all four conditions. w=2; n=4. (D) Pooled Apical to basal f-actin ratio for both conditions molded against PDMS or PC. w=2; n=8 (E) Quantitative analysis of maximal (apical) vinculin expression on the various substrates. w=2; n=3-4 All scale bars are 50 µm if not indicated otherwise.

Therefore, we further investigated the effect of the soft surface layer on confluent epithelial layer morphology and cytoskeletal organization on the soft hydrogels and on stiffer hydrogels containing HepMA (Figure 5B). Analysis of F-actin distribution showed a clear effect of the soft surface layer on cytoskeleton organization in particular on the apical surface (Figure 5C). PDMS molding increased the apical to basal ratio of F-actin compared to PC fabricated samples (Figure 5D). This was significant for the stiffer +HepMA hydrogels (Figure S28 C-D, Supporting Information). The bulk hydrogel composition and mechanical properties by themselves did not seem to have a significant impact on the apical to basal ratio.

The importance of a soft surface layer was further substantiated by staining for vinculin, a component associated with the cytoskeleton that mediates in the linkage between focal adhesions and cell-cell junctions and the actin cytoskeleton^[39,40]^ and is essential for lactation in mammary epithelial cells.^[41]^ Maximum apical vinculin expression increased on samples molded against PDMS (Figure 5B, E). In contrast to the F-actin distribution, apical vinculin expression increased by the addition of HepMA showing the combinatory effect of bulk and surface hydrogel properties.

### 2.5 MPA allows us to locally soften and texture hydrogel surfaces

On flat regions, this combinatory effect is also seen on soft PDMS molded GelMA hydrogels where cells started to migrate into the soft substrate, a behavior not observed for PC molded or stiffer hydrogels (Figure 5B top, 6A, S28B, Supporting Information). On curved half spheres produced with 2PP+RM, cells were able to deform the surface, as indicated by a reduced solidity of the basal surface (Figure S33, Supporting Information). However, we did not observe clear invasion into the soft substrate.

To investigate the influence of the soft surface layer introduced by 2PP+RM we fabricated the same geometry with MPA without this layer. Absence of this soft surface layer increased the solidity of the basal surface (Figure S33B, Supporting Information). Additionally, the epithelial layer thickness increased (Figure 6A). On the stiff hydrogels containing HepMA, differences in epithelial layer thickness were less pronounced (Figure S28 G/H, Supporting Information). However, the integrity of the epithelial layer was improved for PDMS molded samples containing a soft surface layer.

**Figure 6:**
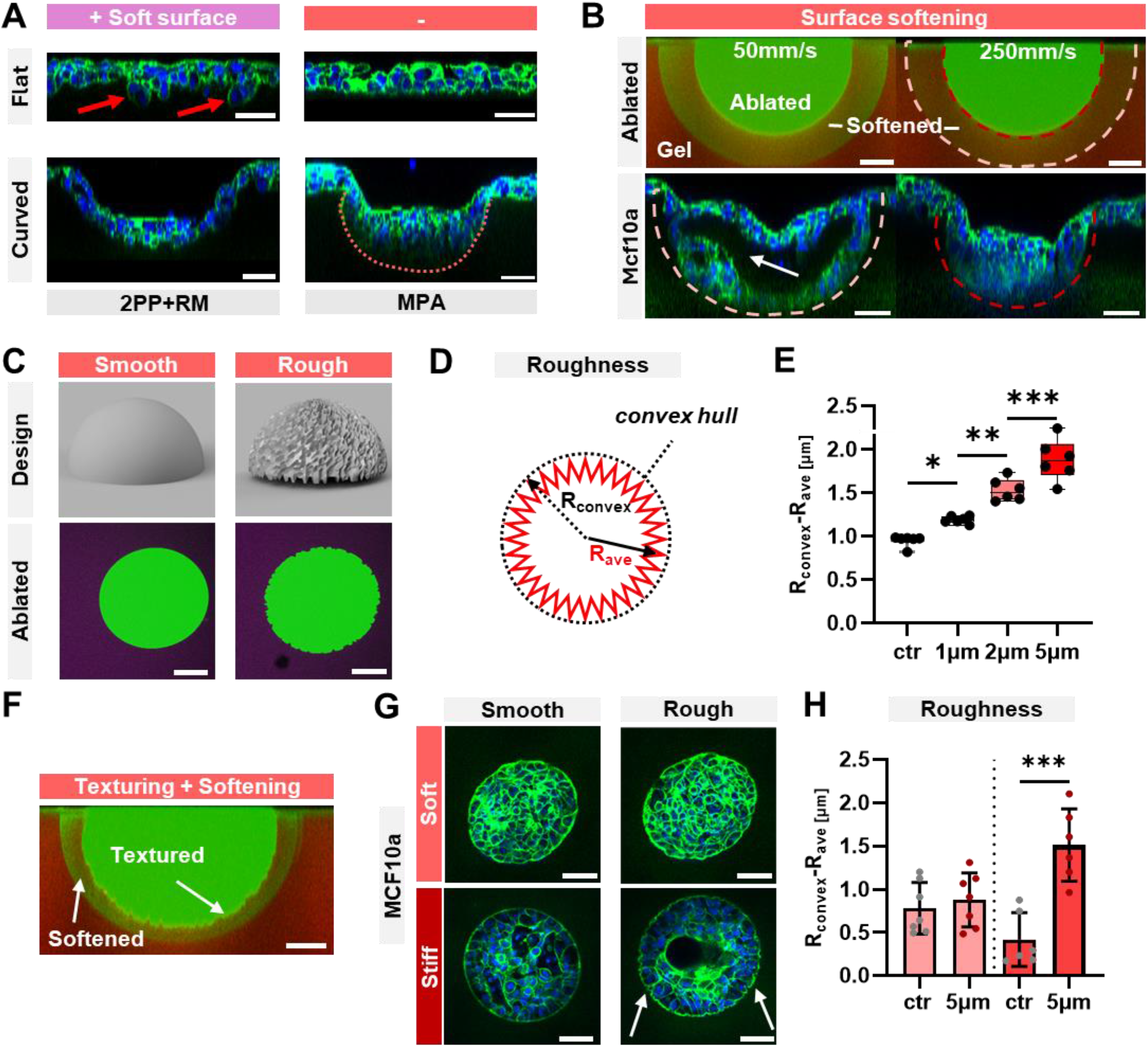
Influence of fabrication induced surface properties on cell attachment and morphology. (A) MCF10a cells on substrates with a soft surface layer (left) and without (right) for flat (top) and curved regions (bottom) on soft hydrogels (Mid DoS). The soft surface layer was introduced using 2PP+RM by molding against PDMS, whereas samples without a soft surface layer were molded against PC and curved features were produced with MPA. Red dotted line indicated basal surface. n=4 (B) Controlled softening of surface by locally varying Laser power (25 mW Softening, 275 mW Ablation). Decreasing the scanning speed from (250 mm/s (right colum) to 50 mm/s (colum) leads to a softer surface layer enabling cells to grow into those areas (bottom row). Dark red shows border of ablated half sphere, light red shows border of softened half sphere. n=3 (C) Design (top) and ablated structure (bottom) in soft hydrogels with a smooth surface(left) and a roughened surface (right). (E) Analysis of the introduced surface roughness for various designs on soft hydrogels calculated by subtracting the average radius from the convex radius calculated from the convex hull area. w=2; n=3. (F) MCF10a cells after 12 days on soft (top) or stiff hydrogels (bottom, Mid DoS +HepMA). Left colum is showing cells on a half sphere with a smooth surface, right colum on a half sphere where we introduced controlled roughness (+/-2.5 µm) w=2; n=3. (G) Roughness of basal surface of MCF10a cells for soft (Mid DoS) and stiff (+HepMA) Hydrogels for a controlled smooth surface and an artificially roughened surface. w=2; n=3. (H) Combination of surface softening and texturing in soft Mid DoS GelMA. Scale bars show 50 µm if not indicated otherwise.

While MPA fabricated curvatures do not come with a soft surface layer by default, the method allows us to locally soften the hydrogel surface (Figure 6B). To mimic cell invasion observed in soft GelMA hydrogels molded against PDMS (Figure 5B,6A) we locally softened the surface of a 250 µm half sphere. Informed by previous dose tests (Figure S13, Supporting Information) we varied the scanning speed to introduce high softening (Figure 6B, left) and low softening (Figure 6B, right), as indicated by increased FITC dextran infiltration. Moreover, we can control the thickness of the modified surface layer (Figure S29, Supporting Information). After 12 days of culture, MCF10a cells exhibited growth into the softened matrix only under the higher softening condition (Figure 5B). As observed for curved regions made by 2PP+RM, the cells lined the outer surface of the softened half sphere rather than infiltrating further into bulk hydrogel. The expansion of the epithelial cells into the softened surface appeared to impose mechanical stress on the epithelial layer, as evidenced by delamination within the cell layer.

Furthermore, MPA allows us to locally introduce surface roughness in the stiffness range that is relevant for epithelial tissue engineering (Figure 6C). While previous studies have shown the importance of surface roughness on epithelial cell behavior they have mostly been performed on stiff, synthetic hydrogels.^[42,43]^ Here we introduce controlled roughness in the physiologically relevant stiffness range for epithelial tissue engineering (Figure 6C). We can design half spheres with a smooth surface, but we can also artificially introduce controlled roughness of +/-2.5 µm (5 µm). We quantified our control over the introduced roughness by subtracting the radius of the convex hull from the average radius (Figure 6D) and demonstrated that four groups with significantly different surface roughness can be generated (Figure 6E, S31, Supporting Information). The high spatial control of MPA further allows us to align the introduced roughness (Figure S31, Supporting Information) and combine it with surface softening (Figure 6F).

Lastly, we wanted to investigate how epithelial morphology is changed by the introduced surface roughness. This roughness is retained in the basal surface of MCF10a cells on stiff but not on soft hydrogels (Figure 6G, S30, Supporting Information). We quantified the morphology of the basal cell surface and found a significant difference between cells on smooth and rough surfaces but only for the stiff condition (Figure 6H). The slight increase in stiffness from 3.3 kPa to 5.6 kPa prevented the cells from smoothing their substrate.

### 2.6 Guiding epithelial layer morphology by 2PP+RM and MPA to increase repeatability and study their behavior in complex geometries

Both tools allow us to shape epithelial layers into complex shapes. With 2PP+RM we can fabricate curvature arrays on the centimeter scale allowing us to study the effect of various curvatures in one sample (Figure 7A). This technique comes with the typical rounding of edges (Figure S32, Supporting Information). In addition to the higher control over surface properties established before MPA provides micrometer-scale resolution (Figure 7B). While cells grow into larger structures, they deform smaller surface structures when the matrix is sufficiently soft. For stiffer substrates we were able to keep shapes below 10µm (Figure S32, Supporting Information).

**Figure 7:**
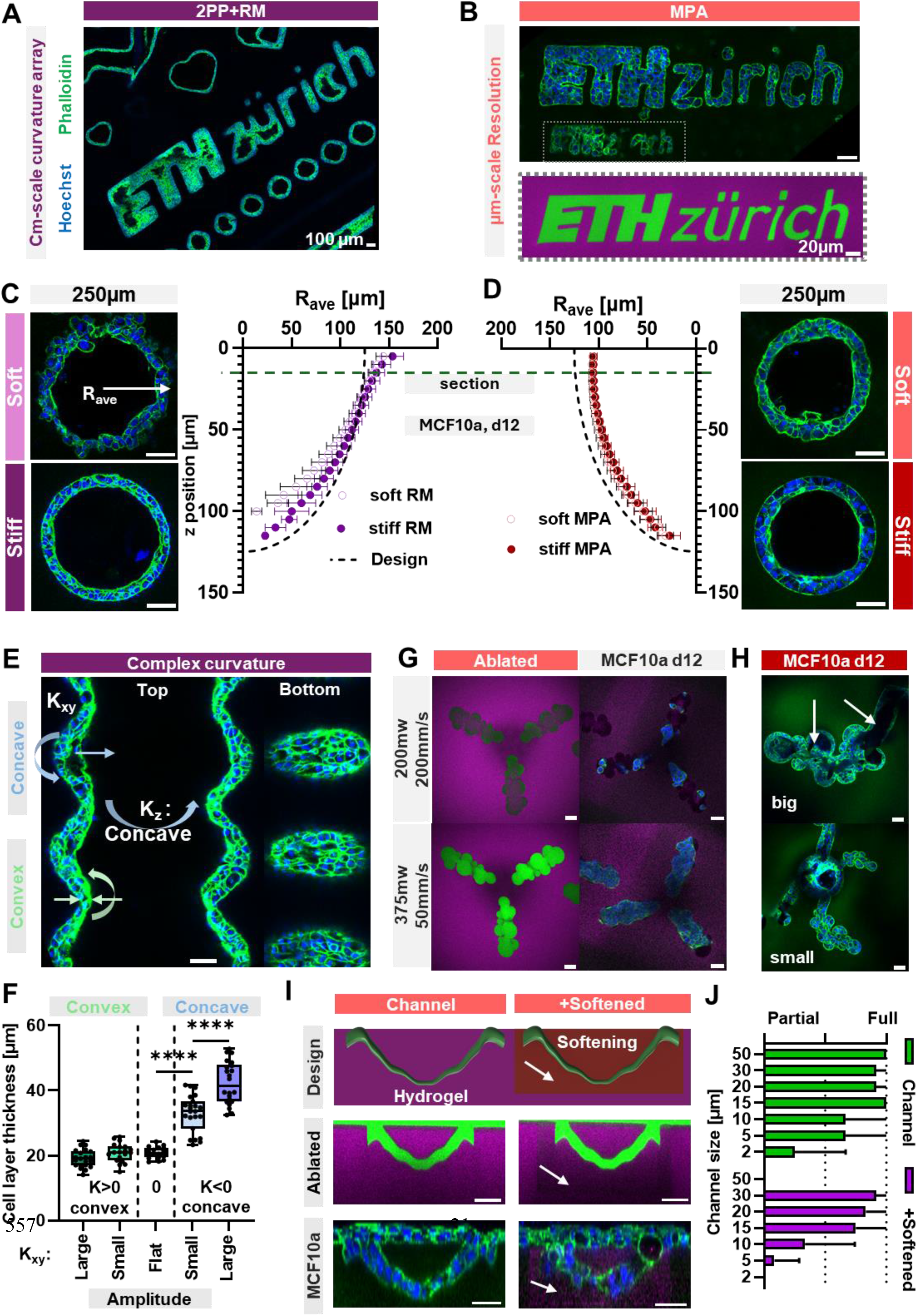
Repeatable guidance of cell morphology with complex substrate curvatures. (A) Confocal z cross-section of replica molded array in Mid DoS GelMA + HepMA gels containing different curvatures including ETH Zürich Logo. n=3 (B) ETH Logo fabricated with MPA after d12 of MCF10a culture in Mid DoS gels and comparison to as fabricated structure. n=3 (C) Cross-section of 250µm half sphere occupied by cells for Soft (Mid DoS) and Stiff (Mid DoS+HepMA) hydrogels and the average outer radius of the cell layer as a function of z position w=2; n=4. (D) 250 µm half sphere fabricated with MPA cultured with MCF10a cells for twelve days. w=2; n=4. (E) Cross-section of MCF10a cells on a complex curvature including concave out of plane curvature and changing concave and convex curvatures in plane as seen in Figure 2D. Crossection at the top (left) and bottom (right) of the complex curvature. w=2; n=3. (F) Cell layer thickness for various curvature regions and for varying amplitude of curvature. w=2; n=3. (G) Fabrication of duct + alveolar like architectures in soft hydrogels (Mid DoS). Top Row: Partial Ablation Bottom Row: Complete ablation. Right column corresponding ingrowth of MCF10a cells depending on the ablation degree. w=2; n=3. Hydrogels are stained with Rhod B (magenta) and perfused with high MW FITC Dextran (green) (H) Cells in alveoli and duct like features of varying size (Top: big, bottom: small) in stiff +HepMA gels, opening of lumen in alveoli and duct-like regions for bigger features size are indicated with a white arrow. n=3. (I) From Left to right: Design of connected channels going into the substrate with (right column) and without softening (left column). White arrows indicate softening. Ablated channels were perfused with high MW FITC dextran (green); hydrogel is stained with Rhod B (magenta). MCf10a cells occupying the ablated channels after twelve days of culture are shown. w=2; n=3. (J) Distribution of channels that were either fully, partially or not occupied for both softened and non-softened channels. w=2; n=3. MCF10a cells are stained with Hoechst (blue) and Phalloidin (green). All scale bars are 50 µm if not indicated otherwise.

To avoid adding to the many sources of biological variance we investigated how repeatably we could culture MCF10a cells on half spheres with a diameter of 250 µm (Figure 6 C&D). We calculated the average radius and its variance of those half spheres for a soft (Mid DoS) and stiff conditions (Mid DoS + HepMA) for both 2PP+RM and MPA. Although curvatures produced with 2PP+RM showed a higher variance on the basal side directly in contact with the substrate, no significant difference was observed between the two techniques on the apical surface (Figure S33, Supporting Information). For 2PP+RM, a considerable fraction of the total standard deviation (0.39) came from the basal surface, whereas for MPA we could not allocate any deviation to the basal surface. For the soft replica molded hydrogels, we once again see deformation of the surrounding matrix (Figure 7C) as quantified by a lower solidity (Figure S33, Supporting Information). Interestingly, this did not have a significant effect on the variability of the basal surface. Additionally, this did not significantly affect apical solidity. The only significant difference between the solidity of the apical surface was when we pooled the stiff conditions together and compared them to samples on soft substrates (Figure S33, Supporting Information) with a decrease in apical solidity for softer samples.

The possibility to shape mammary epithelial cells repeatability into complex architectures allows us to study their effect on the epithelial cell layer. To assess this, we generated geometries with a concave curvature out of plane and switching concave and convex curvatures in plane (Figure 2E) in GelMA +HepMA gels with 2PP+RM and seeded them with MCF10a cells (Figure 7E). We found an increase in cell layer thickness for regions with two concave curvatures (ellipsoidal shape) and not for regions with one convex curvature (saddle shape) (Figure 7F) Moreover, we observed that having concave curvature in one direction and no curvature in the other (i.e. a cylindrical shape) leads only to a slight non-significant increase underlying the importance of two concave curvature axes. Additionally, a higher amplitude of the curvature increased the epithelial layer thickness even further (Figure 7F, S34 Supporting Information).

Similarly, we managed to guide mammary epithelial cells into complex duct and alveoli-like shapes using MPA (Figure 7G). First, we controlled cell infiltration by partial or complete ablation of the geometries. While MCF10a cells migrated into the construct after twelve days the cells were not connected in the partially ablated construct (Figure 7F, right column). On the contrary, in fully ablated constructs cells grew in completely. We further varied the size of the glandular architectures for stiff hydrogels containing HepMA (Figure 7G) as well as soft hydrogels (Figure S35, Supporting Information). For both conditions we observed that smaller sizes were fully occupied as seen in the previously ablated structures whereas there was lumen opening observed in the larger glands. Lumen clearance was more pronounced in duct like areas with cylindrical curvature whereas individual lumens were not fully connected in the alveolar area with two concave curvature axes.

Lastly, MPA allows us to study cell infiltration and its dependency on cavity size, mechanical properties and material composition independently of matrix breaking. To build on previous findings (Figure 5B, 6B, 7G) that softening enhances cell infiltration we successfully ablated channels with smallest dimension varying from 2 µm to 50 µm in soft (Figure 7I) and stiff hydrogels (Figure S36, Supporting Information). For soft hydrogels, cells fully occupied the channels down to a size of 15 µm and partially covered it even for some 2 µm sized channels (Figure 7J, S36, Supporting Information). For stiff hydrogels cells fully covered the channels down to 5 µm. When partially softening the matrix around the channels we observed less effective occupation of the empty channels for both conditions (Figure 7J, S36I, Supporting Information).

## 3. Discussion

In this study, we introduced two multiphoton based tools to produce complex, centimeter-scale curvature arrays with feature sizes between 10 and 250 µm: 2PP+RM and MPA. Both methods enable the repeatable fabrication of concave curvatures in a range of hydrogel materials. 2PP+RM reduces the time needed to fabricate centimeter-scale curvature arrays to only a few minutes. Even though the hydrogel demolding step introduces deformation, the low variance (2.8 µm for 150 µm half sphere) would allow easy adaptation of the design to compensate for this effect. Further reduction in variance could be achieved by controlling the mold quality, selecting only molds that meet a predefined size range. With MPA we achieve comparably low variance without strong deviations from the original design. However, higher geometrical fidelity and flexibility come at a cost of increased printing time. While we made it feasible to ablate tens of half spheres with diameters between 50 and 250 µm in minutes using MPA, only 2PP+RM currently enables realistic generation of fully patterned centimeter-scale curved substrates suitable for large-scale cell analysis.

Both approaches enable us to repeatably fabricate curvatures in various materials relevant for tissue engineering. In comparatively stiff hydrogels, 2PP+RM performs best, largely independent of chemical composition, whereas MPA struggles for purely UV crosslinked gels, but it is adaptable to soft, viscoelastic hydrogels. MPA is nonetheless compatible with stiff hydrogels, provided that the hydrogel network contains bonds which can be efficiently broken, with thermal crosslinks being more easily disrupted compared to chemically crosslinked bonds. Previous studies have either focused on protein-based hydrogel or introduced synthetic photocleavable bonds to accelerate MPA.^[22,25,27,28]^ We find that neither is necessary to rapidly ablate micro to millimeter-scale geometries in soft hydrogels. By optimizing ablation parameters, sensitizer concentration and material composition we were able to decrease print time by orders of magnitude compared to previous reports using photosensitive groups.^[22,27]^ In particular, we highlight the importance of hatching distance in multiphoton based biofabrication while previous work has focused solely on laser power and scanning speed.^[22,27,28]^ Regarding material composition, we demonstrate the excellent compatibility of GAG-based hydrogels with MPA. As GAGs are widely present in many tissues, this compatibility allows us to shape curvatures in materials with physiologically relevant biochemical cues.

Importantly, we achieved all of this in soft hydrogels between 2 and 6 kPA with the possibility to generate a surface layer below 1 kPa, influencing cell adhesion. Leveraging combined effects of thermal and UV crosslinking against a PDMS mold allows us to control this surface layer. The combination of “stiff” bulk and soft surface allows great handling while providing physiologically relevant mechanical cues. For breast tissue engineering, this is in the stiffness range ∼700 Pa to 6 kPa,^[15,20]^ which has been difficult to achieve so far. Our approach gives us the opportunity to shape soft hydrogels in this range into complex curved substrates.

We find that combining a relatively stiff bulk hydrogel containing HepMA (6 kPa) and a soft surface layer improves apical to basal polarization and cell morphology. Single mammary epithelial cells react to increasing stiffness by recruiting more vinculin to the focal adhesion.^[44]^ In contrast, in our confluent epithelial layer we did not observe changes in vinculin expression on the basal side, while it strongly increased at the apical side of the cells. This difference reflects the distinct behavior of epithelial cells in confluent layers. As cell density increases, forces are redistributed across the tissue through adherens junctions rather than being confined to basal focal adhesions. Thus, increased apical vinculin recruitment on stiffer substrates is not unexpected.

However, more surprising was our finding that vinculin expression increased on substrates featuring a soft surface layer. Energy dissipation appears to be an important property for cells as reflected by decreased cell area, yes-associated protein (YAP) nuclear localization and focal adhesion number of MCF10a cells cultured on viscoelastic substrates.^[45]^ The nonlinear stiffness introduced here, combining a 400 Pa soft surface layer and a 6 kPa bulk stiffness, may promote energy dissipation in a comparable matter. The shift in distribution of F-actin towards the apical surface further highlights the effect of a soft surface layer on cytoskeletal organization. In contrast, softer bulk substrates including a soft surface layer enable cells to infiltrate the underlying matrix.

Within the stiffness range relevant for breast tissue engineering (700-6 kPA)^[15,20]^ we observe pronounced differences in the ability of cells to infiltrate their substrate. Softer bulk substrates, including a soft surface layer, enable cells to infiltrate the underlying bulk matrix in flat regions. In curved regions produced with 2PP+RM, cells deformed the soft substrate but less infiltration into the underlying matrix was observed. We confirmed this using concave half spheres produced by MPA, where cells expanded into the locally softened matrix but not beyond, a finding supported by previous work.^[15]^ In contrast, we see less occupation of pre-ablated channels when the surface was softened with MPA consistent with the lower cell coverage on samples with a soft surface layer. Together, these findings suggest that higher cell infiltration in hydrogels with a softer surface is primarily driven by the cells’ ability to deform and break the underlying network which could be relevant in cancer models.

The cells’ ability to deform a softer (3.3 kPa) but not a stiffer (6 kPa) substrate is further demonstrated by the retention of 10 µm features in the later. We observed a similar finding when using MPA to locally roughen the surface. For our softer hydrogels the introduced roughness had no lasting effect as cells were able to smooth the surface. It suggests that they do not prefer a rough surface since they actively reduce the roughness. In contrast, slightly increasing substrate stiffness to 6 kPa prevented cells from completely smoothening the surface. It is striking how a small change in stiffness within their preferred adhesion window inhibits the cells from altering their surroundings. Fibroblasts on a 2D substrate have shown decreased deformation on stiffer hydrogels, but over a much larger window (6-120 kPA).^[46]^ Similarly, Epithelial cell sheets were able to deform substrates with a stiffness of 30 kPa.^[47]^ While the observed difference to previous work might be due to the 3D environment, we cannot rule out the effects of the different hydrogel substrate. Notably, we did not observe clear morphological changes in the epithelial layer itself. Future work should therefore examine whether different components involved in mechanotransduction pathways are modulated by surface roughness.

The curvature of the substrate also influences the epithelial layer. In contrast to previous studies we were able to controllably produce features with two concave curvature axes.^[3,4,11]^ Such curvatures led to a doubling of the epithelial layer thickness considerably higher compared to the slight increase on cylindrical curvatures we and a previous study have found.^[3]^ This gives the space for the formation of multiple cell layers as found in breast alveoli. Additionally, we demonstrate the importance of curvature amplitude which further increases the space occupied by the cells. Differences in apical, lateral and basal stresses explain the effect of curvature-dependent effects on cell density and chromatin packing.^[3,4]^ Stronger bending of the epithelial layer, introduced by two concave curvature axis and higher amplitudes likely amplify these stress imbalances thereby enhancing effects of curvatures. These findings suggest that future work can deliberately tune curvature to control available space and mechanical stress, to form a differentiated multilayer. However, curvature induced stresses must be carefully managed as they can negatively affect the integrity of the epithelial cell layer. Our two tools allow one to find the right material, surface and curvature combination balancing these stresses. Additionally, the high repeatability of both 2PP+RM and MPA will facilitate the search for the mechanistic link between curvature and functioning tissues, for which many indications exist.^[1-4,15,48]^

By leveraging the multiphoton principle, we achieve high spatial resolution and reproducibility while substantially increasing print speeds. This approach enables the production of complex curvatures that maintain low standard deviation even after prolonged cell culture while still allowing the cells to reshape their environment. This reduces fabrication and material induced variability which historically has limited biological interpretations. These platforms provide scientists with robust tools to study effects of ECM composition and drug responses in a physiological relevant environment. Ongoing advances in hard- and software of multiphoton printers will further accelerate speed and scalability in fabricating precise 3D environments for epithelial tissue engineering.^[49-51]^

## 4. Materials and Methods

### 4.1 Mold fabrication

#### 4.1.1 Design

Curvature arrays and architectures were designed in Fusion 360 (Autodesk) and exported in high quality.

#### 4.1.2 NanoScribe Printing

Single-side polished silicon wafers were used as substrates. Prior to printing, the wafers were sequentially rinsed with acetone and isopropanol, dried under a nitrogen stream and plasma-treated for 30 s. A commercial direct laser writing setup (Photonic Professional GT2, NanoScribe GmbH) with a 10x/ 0.3NA objective with commercial Dip-in resin (IP-Q, NanoScribe) was used for template fabrication. The structures are fabricated at a printing speed of 10000 µm/s and 40% laser power. A slicing distance (hatching angle 90°, layer height) of 0.3µm and a hatching distance of 0.2 µm were used. The used laser had a 780nm wavelength, a pulse duration of 100 fs and a pulse repetition frequency of 80MHz. After fabrication, the samples were developed in propylene glycol monomethyl ether acetate (PGMEA) for 15 min to remove unpolymerized resist, followed by rinsing in isopropanol for 5 min. After drying of with a nitrogen air gun, the developed substrate is post-cured under UV light [306 nm] for at least 1 h. To decrease the adhesion of glass substrate and the mold, Trichloro[1H,1H,2H,2H-perfluorooctyl] silane was coated on the substrate by means of chemical vapor deposition, by exposing the substrate to a 100 µL droplet of the silane for 30 min and rinsed with isopropanol afterward.

#### 4.1.3. Upnano Printing

Molds were printed with commercial Upphoto resin (Upnano GmbH 4302-8-001) in VAT mode on a NanoOne1000 printer (Upnano). The Positive features were printed on top of a 1mm base plate printed with the same resin to ensure mold stability for repeated molding. A 10x air objective was used (UPLXAPO10X NA:0.4) with recommended coarse settings (LD:4.2 µm, LH:5 µm, Power 100 mW, Speed: 600 mm s^-1^). Printing files were sliced with Think3D software (Upnano). Printed parts were washed twice in IPA until no resin was diffusing out.

Molds were post treated in a UV box for 1h at 90 mW cm^-2^ and put on a heat plate at 50 ºC for 2 h.

#### 4.1.4 PDMS Molds

Polydimethylsiloxane (PDMS, Sylgard 184, Dow Corning) was mixed 10:1 in a Thinky mixer at 3000 rpm for 5 min and degassed in a desiccator before pouring onto the mold. After an additional 1h degassing step the PDMS was put into an oven for 24 hours at 65ºC. The negative PDMS mold was intercalated, washed with EtOH and later Plasma treated to avoid binding of the positive PDMS mold. Directly after Plasma treatment the molds were put into a 70% EtOH, 30% H2O solution to stabilize the hydrophilic surface. To obtain the final positive mold the process was repeated curing PDMS on top of the negative mold. For the negative molds printed with the Upnano printer the first step was skipped.

### 4.2 GelMA Synthesis

Gelatin type A from porcine skin (Sigma Aldrich(SA), G2500), 20g, was dissolved in 200ml cold 0.25mM Carbonate-carbonate buffer, pH9 (Sodium carbonate anhydrous; VVR 20K204120, Sodium hydrogen carbonate; Fisher 2065599) and mixed for 1h at 50ºC. Methacrylic Anhydride (SA, 276685-500) was added 5 times with 30min intermissions under 600rpm stirring according to table S1 to achieve the desired degree of substitution. The pH was measured (FiveEasyTMFE20, Mettler Toledo) and adjusted to 9 before each subsequent addition with 1M HCL and NaOH solutions. After the last addition the reaction was left for 1h and diluted with 200ml preheated distilled water (miliQ, ELGA Purelab Ultra Genetic) and pH was adjusted to 7.4. Unreacted MAA was separated by centrifugation (40 ºC, 3500 rpm) and filtrated (Vacuum filtration 500, TRP 99500). Next, the synthesized GelMA was dialised (Dialysis tubing, MWCO 12400, 99.99% retention, Merck 3110) against 5l of DI water for 4 days with 2-3 water changes a day.

### 4.3 Hydrogel curvature array fabrication

PDMS molds were washed and cleaned and sterilized with 70% EtOH under UV light for 30min together with an 11 mm inner diameter PDMS ring. The Molds were later dried under aseptic conditions. GelMA was dissolved in PBS 7.4pH (1x) (Gibco 10010-015) at 50mg/ml with 50 µl/ml Lithium-Phenyl-2,4,6-trimethylbenzoylphosphinat (LAP, SA 900889) and 1 µl/ml Acryloxyethyl thiocarbamoyl rhodamine B (Rhod B, 10mg/ml, PolySciences 25404) added. On top 200ul of filtered resin (0.2µm, Sarstedt 501692591) was added and either first exposed to the corresponding thermal treatment or directly UV crosslinked (90 mW/cm2 if not mentioned otherwise) in a UV Box (405nm) for 5-30min. HepMA (0.2-0.5 mmol/g ester groups, potency 180U/mg, Matexcel BINK-020) was added at 10 mg/ml. Collagen (TeloCol®-3 Advanced Biomatrix #5026) was used at a concentration of 3 mg/ml and was neutralized with the provided Solution (Advanced Biomatrix 5163) and thermally cured according to the manufacturers protocol. DECM was extracted from bovine udders according to a previously established protocol. It was dissolved in PBS 1x at 40 mg/ml together with 10 mg/ml HepMA, 0.05 w/v% LAP 5 mM Sodium Persulfate (AdBio #5248) and 0.5 mM Ruthenium (AdBio #5248). After curing the hydrogels PBS 1x was added on top to avoid sticking of the material to the PDMS ring, which was removed with a spatula. Samples were then intercalated with PBS and transferred into a well plate where they were washed and kept for storage at 4C until further use. Cell culture media was added 1 day prior to cell experiments.

### 4.4 Mechanical testing

#### 4.4.1 Rheology

[Photo]rheology tests were measured on an Anton Paar MCR 302e rheometer with a 20mm parallel plate (Anton Paar 3049). UV curing was performed with an Omnicure Series1000 lamp (Lumen Dynamics) with 10 mw/cm2 (60%) equipped with a widepass filter 400-500 nm and a 405 nm filter. Per test 85 µl of sample were loaded on to a bottom glass plate with excess resin being trimmed away. A wet tissue was put at the edges of the sample chamber to avoid drying over longer measurements. Oscillatory measurements were performed with a gap size of 0.2mm, a shear stress of 1% and an angular frequency of 5 rad/s. Samples were either pre-crosslinked for 1h at 23ºC or 4ºC respectively or directly UV crosslinked after 5min of equilibration time.

#### 4.4.2 Compression tests

Stress strain curves were measured in compression with a TA.XTplus Texture Analyzer [Stable Micro systems] with a 500 g load cell. A preload of 0.3 g was applied to assure sample contact. Samples were compressed to 15 % with a strain rate of 0.01 mm/s. Tests were repeated 5 times per sample with a total of 3 samples per condition. Young’s Moduli were calculated from the linear region of by applying a linear fit to the 0-3 % strain region.

#### 4.4.3 AFM

Mechanical properties were measured using a Bruker Dimension Icon AFM system. Tipless Au-coated cantilevers (CSC-38, Mikromash, Bulgaria) were used for the measurements. Cantilever normal spring constant was determined using the thermal-noise method and found to be 0.152 N/m.^[49]^ A 12 µm diameter silica particle (EKA Chemicals AB, Kromasil R) was attached to the end of the tipless cantilever using two-component epoxy glue using a home-built micromanipulator. The prepared colloidal probe was treated with UV/ozone for 30 min before the measurement. Indentation tests were performed on cells that were immersed in PBS 1x and measured over multiple arrays over the surface. Approximately 6 indentation curves were recorded and fitted using the built-in function of Nanoscope analysis software (Version 9.2) with the Hertz model to determine the Young’s modulus.

### 4.5 Ablation

Hydrogel resin was molded against a PC Petri dish in a PDMS ring, left at RT for 1h and UV cured under UV light (see Supporting Information for exact curing details & material composition. Hydrogels were washed twice in PBS 1x (Gibco) and transferred to a µ-Slide 8-well high glass bottom (Ibidi 80807). A stock solution of 10 mM P2CK (5.06 mg/ml, 506.4846 g/mol in 1x PBS) was diluted with PBS 1x containing 1 v/v% penicillin/streptomycin (Pen/Strep; Gibco, 15140122). P2CK was left to diffuse in for at least 24h at 4ºC. Before printing the samples were left at RT for at least 30 min after which the P2CK solution was removed. Ablation was performed with the NanoOne 1000 system and a 20x 0.4Na water object in top-down mode at the glass slide hydrogel interface. A refractive index of 1.36 for GelMA was taken from literature. Coarse mode and a line width of 100% was used when not stated otherwise. Laser parameters were 780 nm; 90 fs; 80Mhz respectively. Samples were transferred to a well plate and kept for a week in the fridge and washed in PBS multiple times to remove P2CK. The maximum speed available for the 20x objective was reduced to 400 mm/s with v2.4.4 of the Think3D software (Upnano).

### 4.6 Substrate Imaging

1µl/ml Rhod B Stock was added to stain PDMS molds and hydrogels. Cavities were perfused with 2 M Da FITC Dextran [Merck]. Samples were imaged with a Leica SP8 system with a 25x water objective. Average radius of curved features was calculated from the stained area at every z position of the z stack. Three samples were imaged with at least two geometries per sample.

### 4.7 Cell culture

A stock of MCF10a (ATCC CRL-10317) cells were kindly provided by the Bodemiller lab, ETHZ and expanded in DMEM/F12 1:1 1x (Gibco 31330-038) containing 5v/v% Horse Serum (New Zealand Origin, ThermoFisher(TF), 16050122) 1v/v% Pen/Strep, 500 μl Insulin (10μg/ml, SA #I-1882), 250μl Hydrocortisone (0.5 µg/ml, SA #H-0888), 50μl Cholera toxin (100ng/ml, SA #C-8052) and 100μl rhEGF (20ng/ml, Peprotech AF-100-15) according to a Brugge lab protocol. Cells were used between passage 12-20. For cell seeding, cells were washed with PBS 1x and trypsinized for 5 minutes and spined down. Cells were counted and resuspended in 20 ul per sample with a seeding density of 120k cells/sample. Well plates were put into incubator (37 Cº 5% CO) for 25 min to allow initial cell attachment after which Media was added. Media was changed every 1-2 days. Samples were washed twice with PBS and fixed in 4% PFA (30min) after twelve days of culture. Cell attachment was checked at d1 d2 d4 and d12 with a BF microscope (EVOS).

### 4.8 Immunohistochemistry

Samples were washed 2 times with PBS 1x to remove PFA and left at RT for 30minutes. Next, constructs were permeabilized and blocked with a 1% BSA solution containing 0.5% Triton X for 30 minutes. Primary antibodies [tables S2] were diluted in 1%BSA + 0.01% Triton X and left at 4ºC overnight. After 3 washing steps with PBS 1x secondary antibodies and dyes were added in 1wt% BSA + 0.01v/v% Triton X and incubated dark at RT for 1h30min before washing 3 times with PBS 1x. Samples were imaged with a Confocal Leica SP8-AOBS-CARS system.

### 4.9 Data Analysis

Data analysis was performed with at least 3 replicates with the number of replicates being stated in the individual figures. Fiji/ImageJ 1.54p, Matlab 2023a, Microsoft Excel and GraphPad Prism v10.6.1were used to curate and analyze the data. Object solidity was quantified in Fiji/ImageJ using built in shape descriptors. Figures show mean +/-standard deviation unless otherwise stated. Statistical significance was assed using unpaired t-tests and ordinary one-way ANOVA tests for comparison of multiple groups. Results were considered statistically significant with *p<0.05, **p < 0.01, and ***p < 0.001. Non significance is shown with ns.

### 4.10 Graphics and AI tools

The graphical abstract consists solely of schematic illustrations and does not present original experimental data or results. Schematics for the graphical abstract were created using PowerPoint (Microsoft 365) and Biorender [https://BioRender.com]. A large language model (ChatGPT, OpenAI) was used to assist with proofreading; all text and figures were verified and finalized by the authors.

## Acknowledgements

We thank Christian Gehre, Margherita Bernero, Wanwan Qin and Xiao-Hua Qin for the discussions on multiphoton biofabrication. We further thank all members of our laboratories for the discussions and inputs, especially Annalena Maier for her inputs on the P2CK synthesis and help with NMR. We thank the employes of UpNano GMBH for their support on use of the NanoOne1000. We are grateful to Dimitar Boev for his suggestions on the figures.

The authors gratefully acknowledge ScopeM for their support and assistance in this work. M.Z.-W. acknowledges the ALIVE initiative (Advanced Engineering with Living Materials) in the SFA-AM program (Strategic Focus Area – Advanced Manufacturing) and an ETH Foundation Grant (23-1 ETH-12). Lucio Isa and Xueting Shen have received funding from the European Research Council (ERC) under the European Union’s Horizon 2020 Research and innovation program Grant Agreement No. 101001514.

## Data Availability Statement

All data needed to evaluate the conclusions in the paper are present in the paper and/or the Supplementary Materials. Data also is available in the ETH Zurich Research Collection https://doi.org/10.3929/ethz-c-000797770 under the terms of the repository’s data-sharing policies.

Received: [[will be filled in by the editorial staff]]

Revised: [[will be filled in by the editorial staff]]

Published online: [[will be filled in by the editorial staff]]

## Supporting Information

Supporting Information is available from the Wiley Online Library or from the author.

