## Supporting Information for "Complementary multiphoton tools to create 3D architectures in soft hydrogels for epithelial tissue engineering"

**Title**

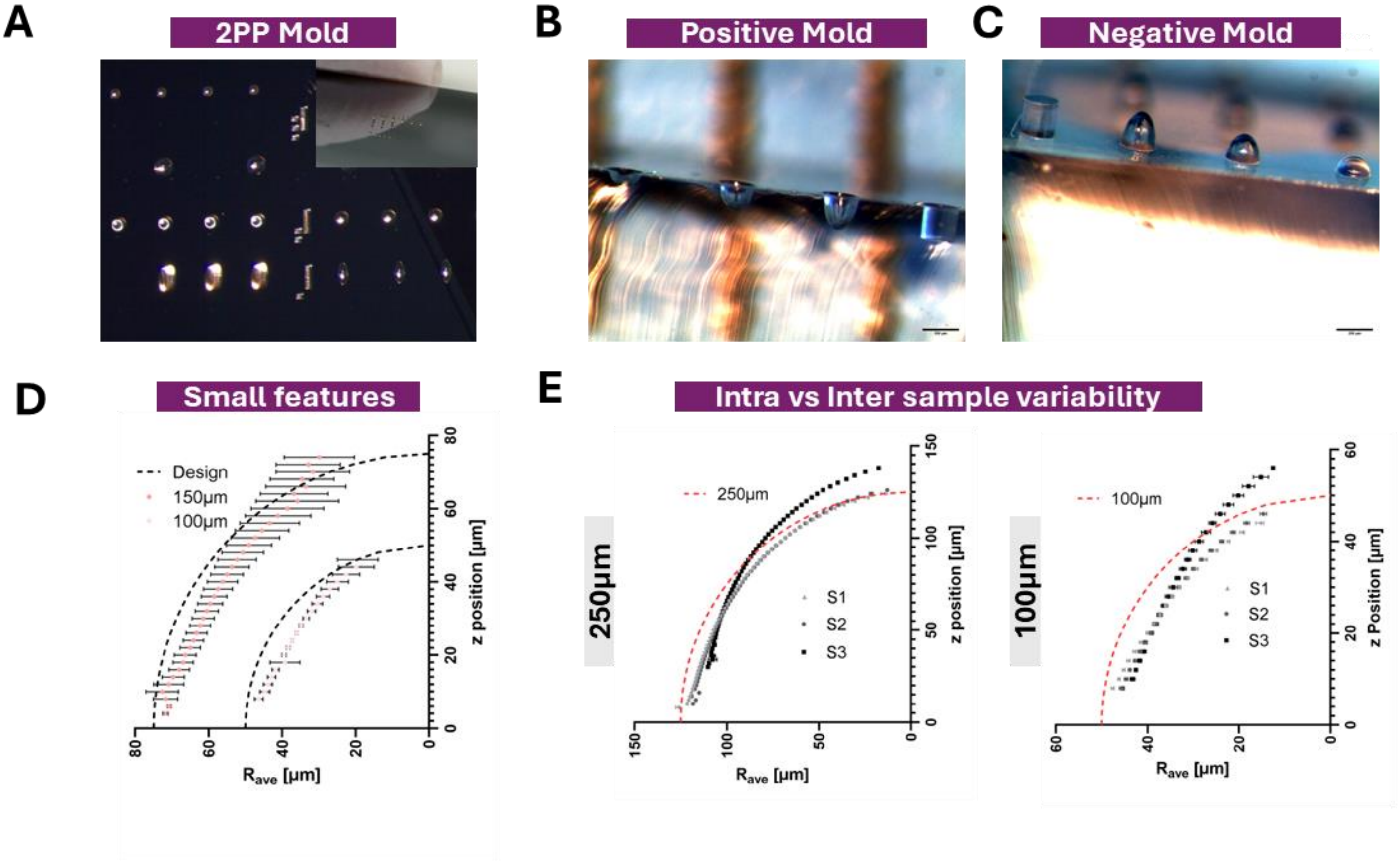


**Figure S1: Two-Photon printing and two step replica molding using nanoscribe.** (A) Negative of curvature array printed on a scilicone waver (B) Crossection of Positive PDMS mold containing features with a diameter of 250µm of varying depth and shape. (C) Negative PDMS mold containing the sample features (D) Repeatability of 100 and 150µm half spheres. Rave was calculated from confocal z-stacks of fluorescently labeled PDMS. (E) Repeatability of features within a sample shown for three replicates. Left: 250µm; Right: 100µm

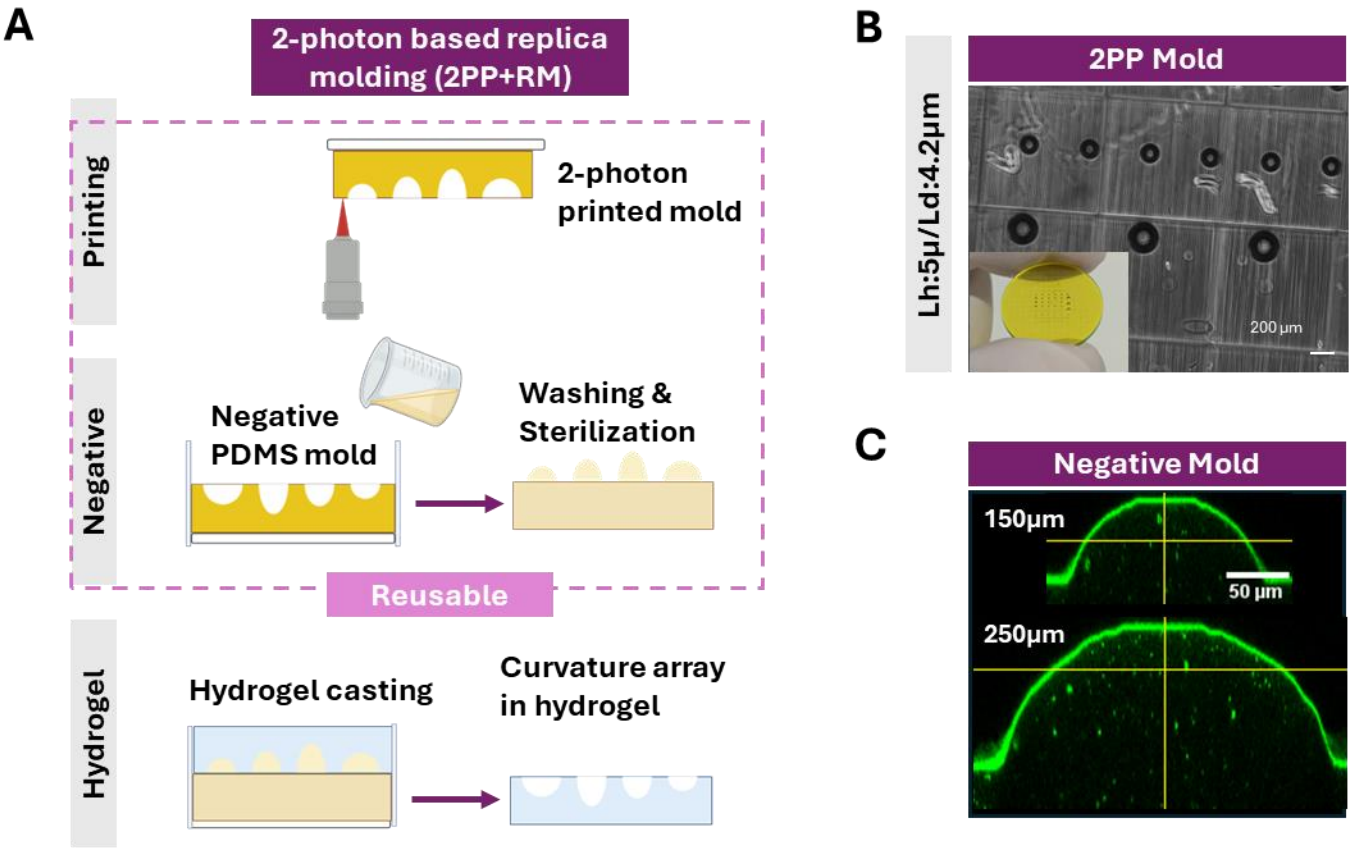


**Figure S2: 2PP-Printing of positive of curvature array and 1-step replica molding.** (A) Adapted protocol from figure 1B. Since a positive mold is printed only one PDMS molding step has to be performed in order to get the desired concave curvatures (B) Positive mold printed with a high layer height and line distance to reduce printing time. Molds were printed on a nanoone1000 (Upnano) using Upphoto (Upnano) in coarse mode. (C) Cross-section of negative PDMS features (150, 250µm) obtained with this adapted process.

To minimize printing time at high surface quality (Line distance (LD) 0.2µm; Layer height (LH) 0.3µm), we first printed a negative of the final concave curvature arrays. As an alternative approach when manual work rather than printing time is the limiting factor, we directly printed the negative mold skipping one PDMS replication step (figure S2) (20). We increase the hatching distance to 5µm and 4.2µm respectively to account for the increased volume, and we printed molds with more complex arrays containing both concave and convex principal curvatures, that were successfully replicated into PDMS. (figure 2C, S3).

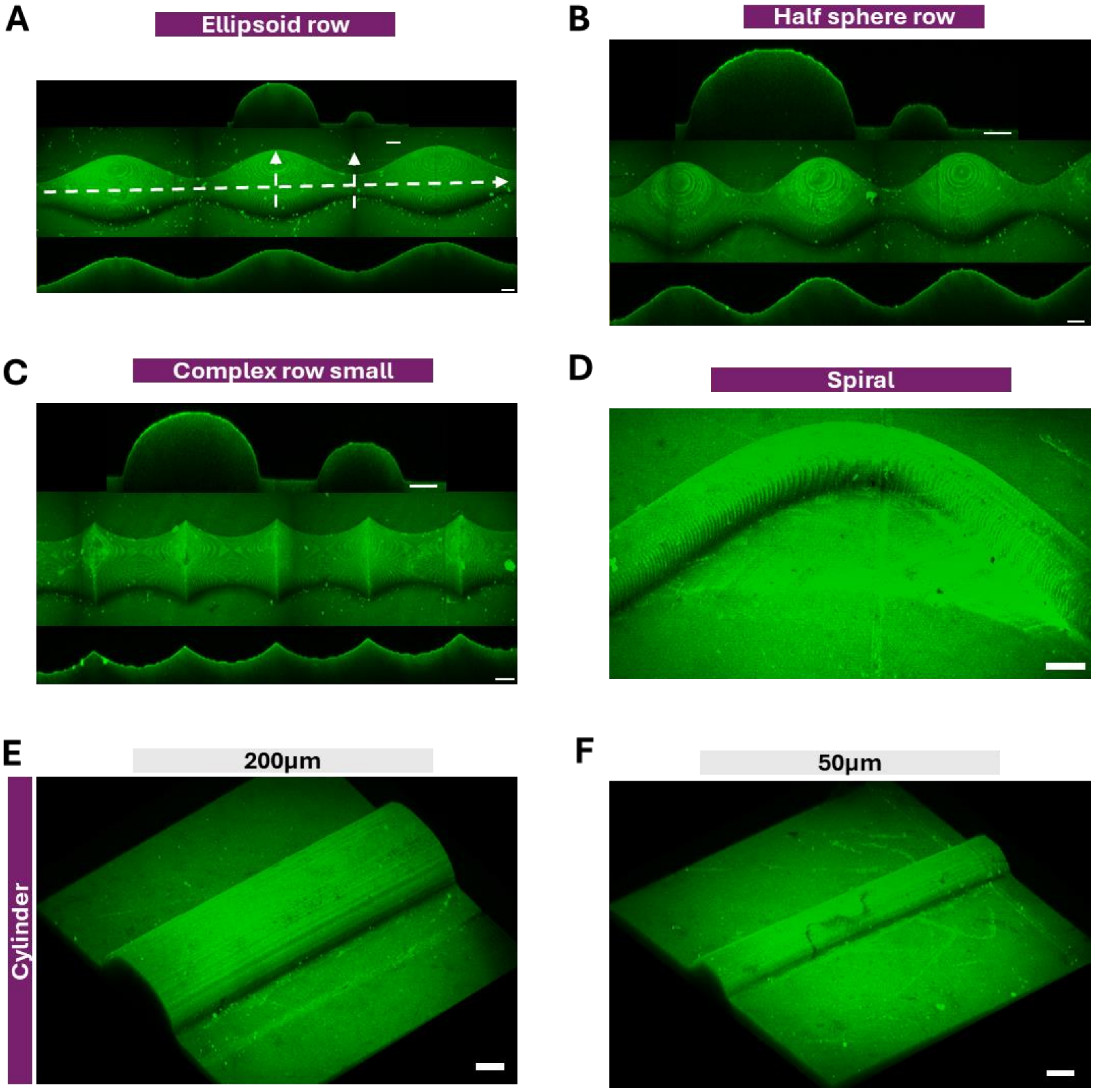


**Figure S3: PDMS molds with controlled curvatures in and out of plane obtained from a one step replica molding approach.** (A) 3d projection and cross-sections (top:axial bottom: longitudinal) of connected ellipsoids in PDMS (B) Connected half sphere row (C) small complex curvature model containing out of plane convex and alternating concave and convex curvature in plane (D) 3D Projection of spirale, (E) 200µm cylinder and (F) 50 µm cylinder. All images were obtained by confocal microscopy. PDMS was stained with Rhod B acrylate.

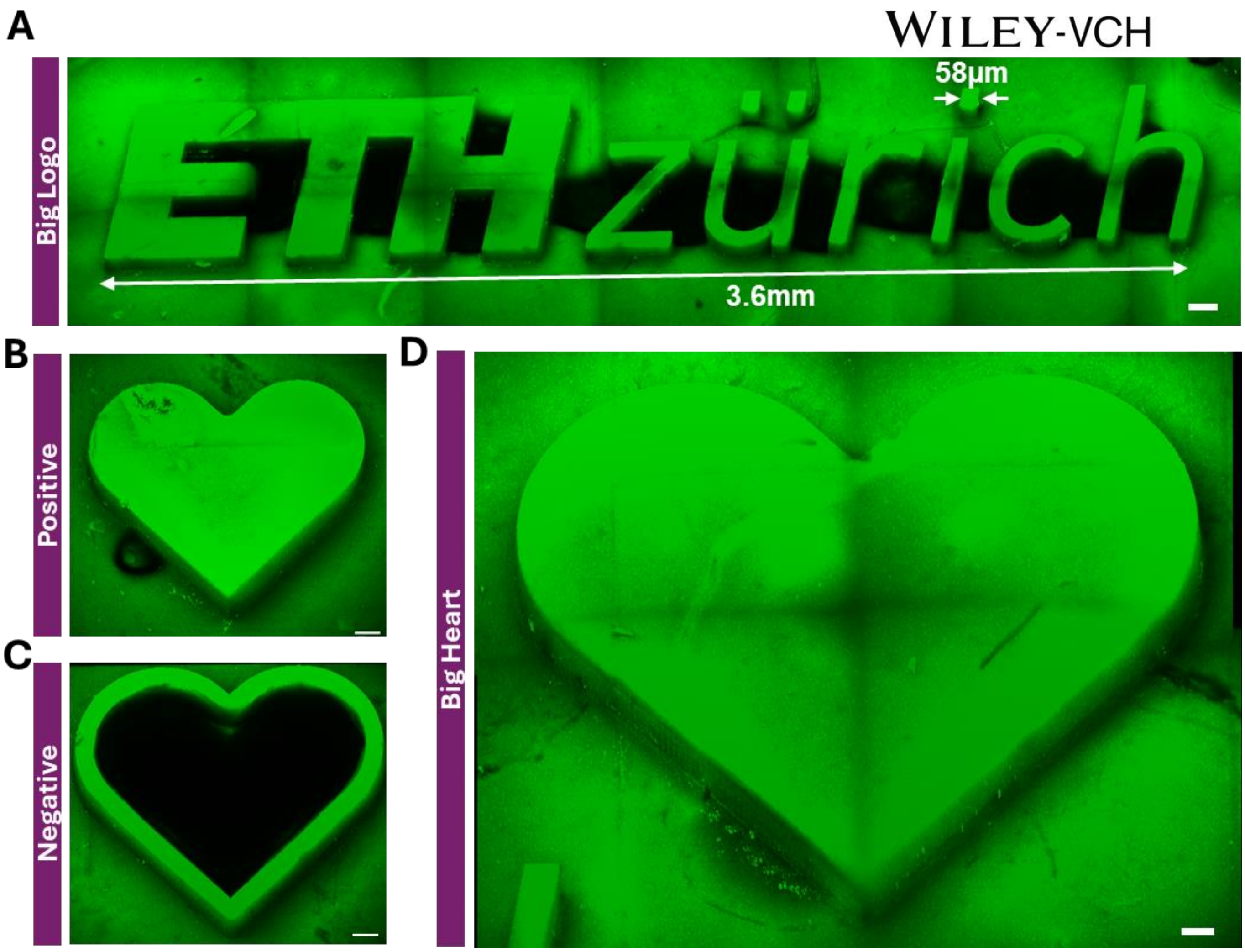


**Figure S4: Arbitrary shapes of containing large and small feature sizes simultaneously.** (A) 3.6mm wide ETH logo containing a minimal feature size of 58µm (indicated by white arrows). (B) Positive heart shape and (C) negative containing only outer wall (D) big heart shape.

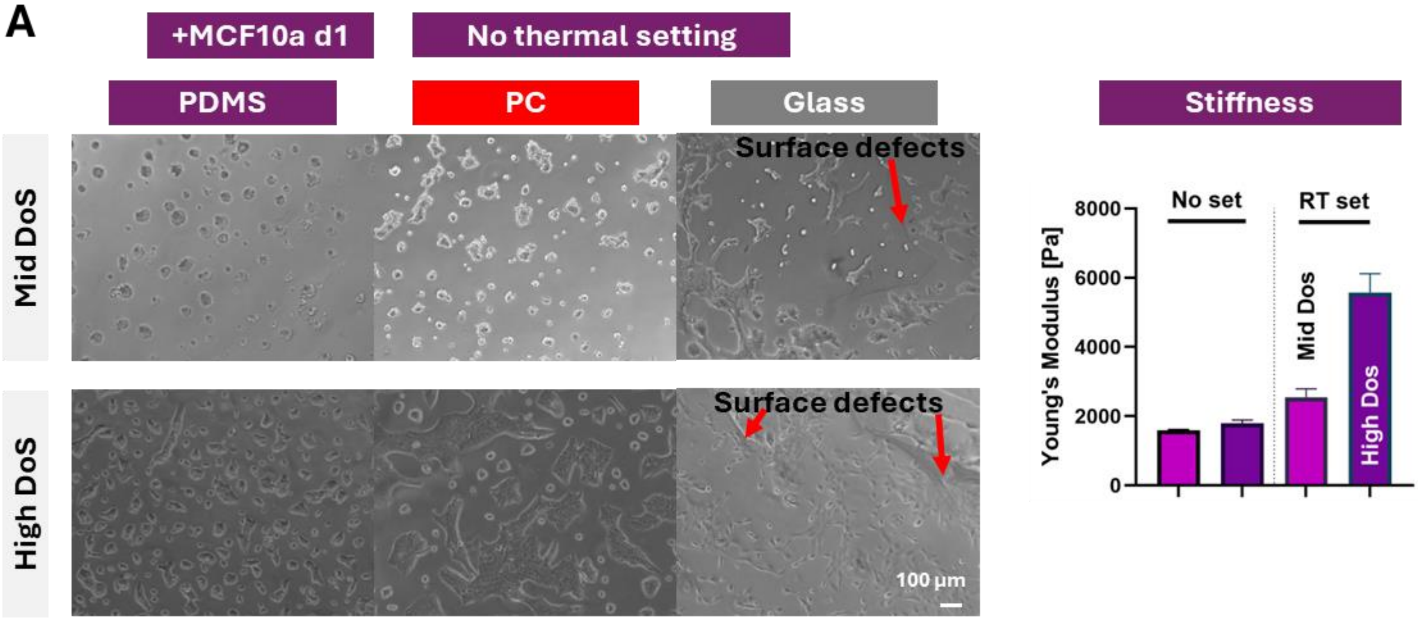


**Figure S5: Attachment of MCF10a cells cultured for 1 day on thermally non crosslinked gels: (A)** Top Row: GelMA Medium degree of substitution, bottom row: GelMA High degree of substistution. From left to right BF images of MCf10a cells on gels molded against PDMS, Polycarbonat (PC) and glass. Surface defects introduced through molding against glass are indicated with red arrows.

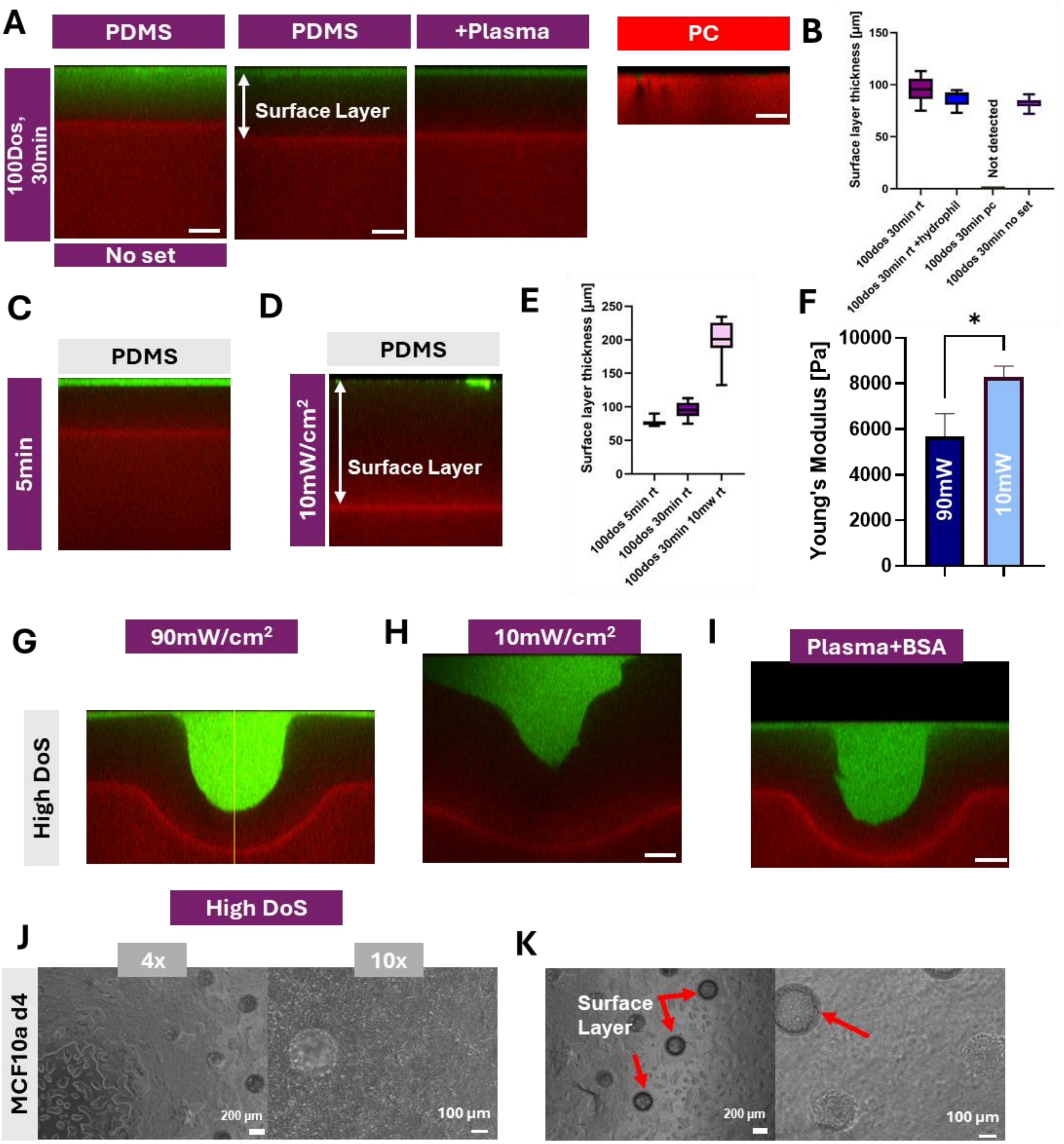


**Figure S6: Surface layer formation in High DoS GelMA:** (A) High degree of Substitution GelMA molded against PDMS (From left to right : non thermally crosslinked; thermally crosslinked, Plasma treated and BSA coated PDMS, Polycarbonate (PC) (B) Surface layer thickness of the 4 conditions (C) Influence of lower curing time (5min instead of 30min) and (D) curing power density on surface layer thickness. (E) Analysis of surface layer thickness (F) Influence of UV power density on stiffness of thermally crosslinked GelMA hydrogels (G) Successully replicated 250µm half spheres in high DoS GelMA fabricated with 90mW/cm2 (H) 10mW/cm2 (I) or Plasma + BSA treated molds (J) MCF10a cells after day 4 on High DoS GelMA, thermal crosslinking + 5min 90mW/cm2 (K) Surface layer present for cylindrical features (indicated by red arrows) as seen in BF image.

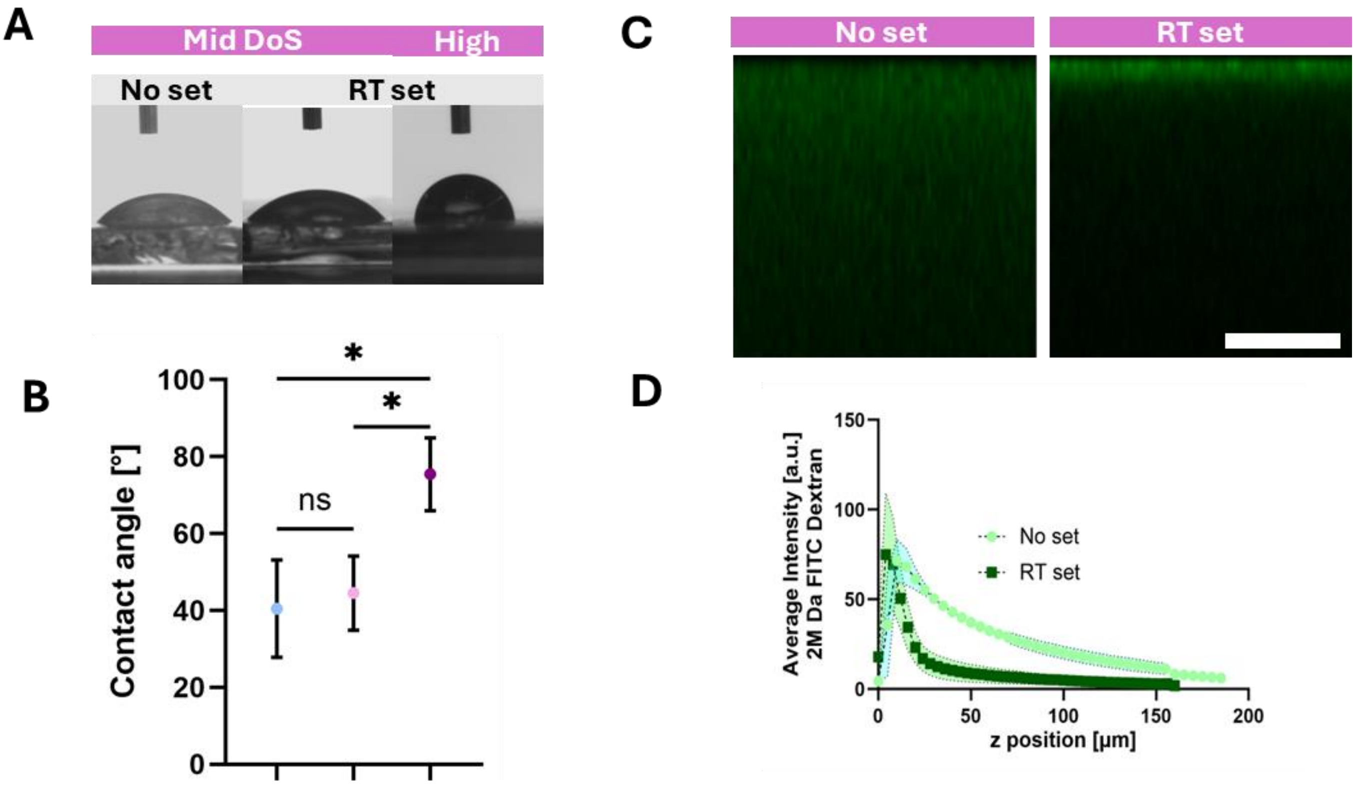


**Figure S7: Analysis of GelMA surface properties:** (A) Sessile droplet measurement on Mid DoS GelMA without thermal crosslinking (left), thermal crosslinking at RT (middle) and High DoS GelMA with thermal crosslinking (right) (B) Corresponding contact angle measured on the three different conditions. (C) Confocal image of surface layer composition perfused with high MW FITC Dextran for Mid DoS GelMA (D) Average Intensity analysis of diffused FITC dye starting from hydrogel surface.

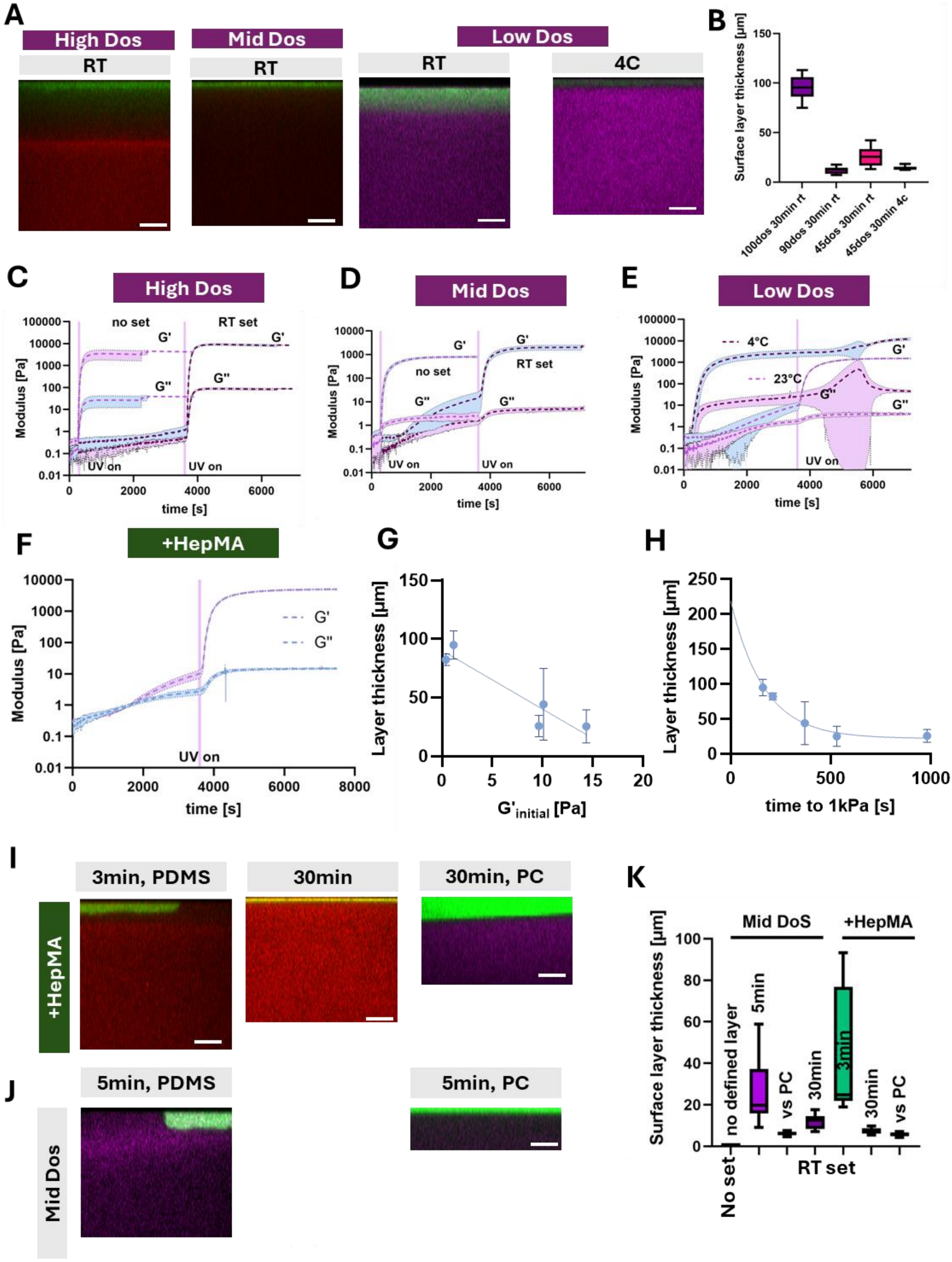


**Figure S8: Influence of material composition on surface layer formation for GelMA based hydrogels.** (A) influence of degree of substitution and set temperature on surface layer formation (B) Analysis of surface layer thickness (C) Rheological data showing curing behaviour for thermal precrosslinking vs direct UV crosslinking (Storage Modulus: G'; Loss Modulus G'') for High DoS GelMA (D) Mid DoS, (E) Low DoS, (F) and Mid DoS+ HepMA. (G) Surface layer thickness for different degree of Substitution pooled vs their initial Storage modulus, linear fit applied. (H) Pooled layer thickness vs the time it takes to reach 1 kPa Storage modulus. One phase decay fit applied (I) Surface Layer thickness for HepMA with varying curing time and against PC (J) Surface layer formation for MiD DoS GelMA for lower curing time (left) and against PC(right) (K) Quantification of surface layer thickness for the shown conditions.

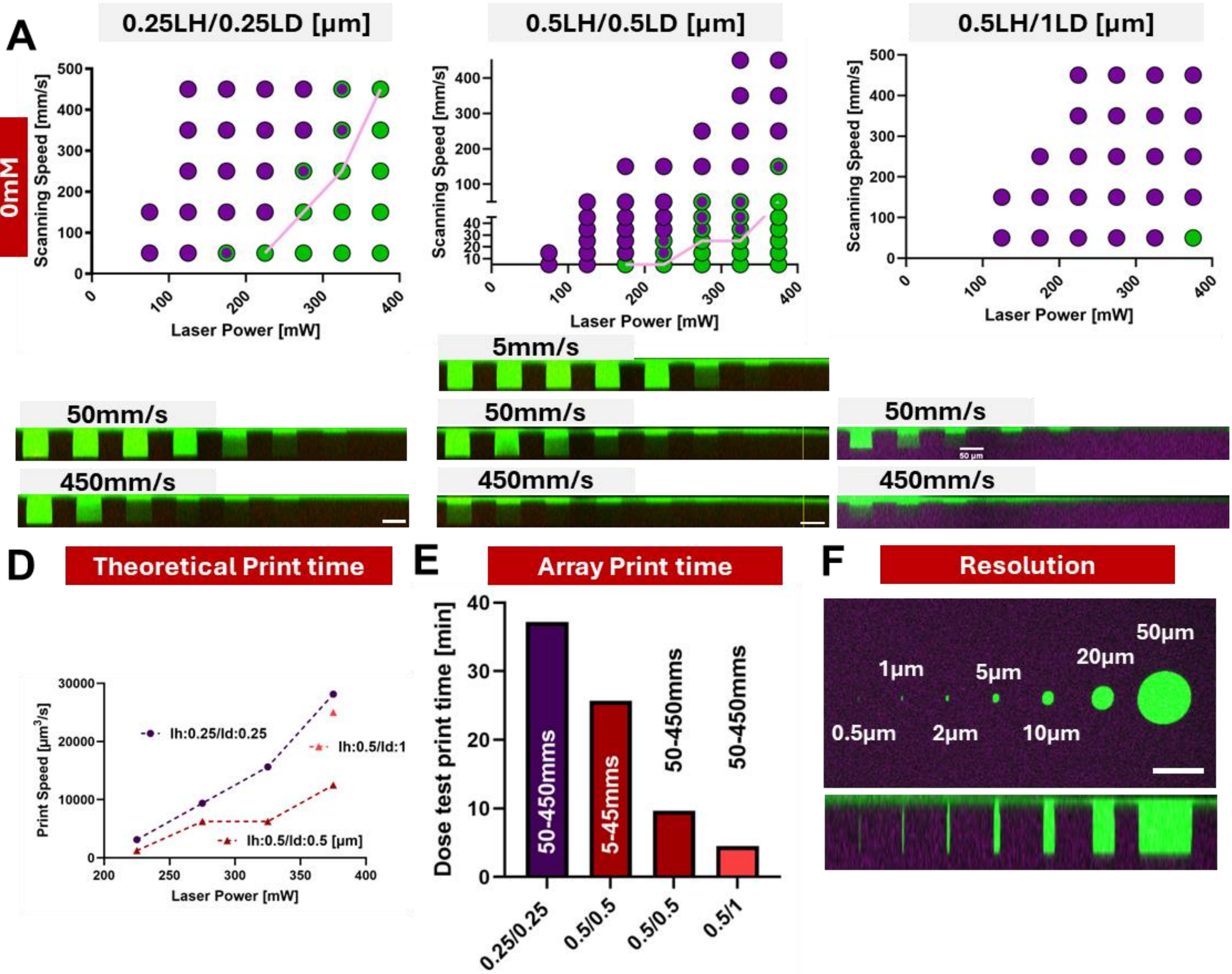


**Figure S9: Parameter screening in Mid DoS GelMA without any Sensitizer:** (A) Dose tests of 50µm cylinders with Laser Power varying from 25mW to 375mW and scanning speed from 50 to 450mm s-1 using a layer height LH of 0.25µm and a Line distance LD of 0.25µm. Top: Median value from 3 samples Bottom: side view of dose test for 50 and 450 mm s-1.(B) Dose test with LH:0.5µm and LD:0.5µm. Scanning speed was varied from 5to450mm s-1. (C) Dose test with LH:0.5µm and LD:1µm. (D) theoretical print time calculated from the maximum speed that allows ablation for a given Laser power for the three tested hatching distances. (E) Actual array print time of dose tests (F) Resolution test: ablated cylinders ranging from 0.5µm to 50µm.

**Ablation in GelMA without sensitizer.**

In figure S9 (Supplementary information) we investigated the potential of ablation in Mid DoS GelMA hydrogels. We found that while being successfully able to ablate 50µm cylinders we needed to choose low LD and LH values (0.5µm) using low scanning speeds of 50mm $s^{-1}$ or even reduce LD+LH further to 0.25µm to use higher scanning speeds. Successful ablation was determined by co-staining with high MW FITC Dextran that does not penetrate into gels and Rhod B that was crosslinked into gels (S10, Supplementary info). Using the maximum speed ($v_{max}$) that allowed ablation we calculated the theoretical printing speed Q using equation 1 including Layer height (LH) and line distance (LD).

$$(1)\ Q_{print} = v_{max} * LH * LD$$

We found that a lower hatching distance leads to a higher print speed due to an increased $v_{max}$. In contrast to this calculation, we find the actual print time required for a dose test to be lower for 0.5µm hatching distance compared to 0.25µm. The scanning laser needs to accelerate and decelerate at each line. For smaller structures this change in speed becomes more dominant, increasing the influence of LH and LD on theoretical print speed compared to $v_{max}$.

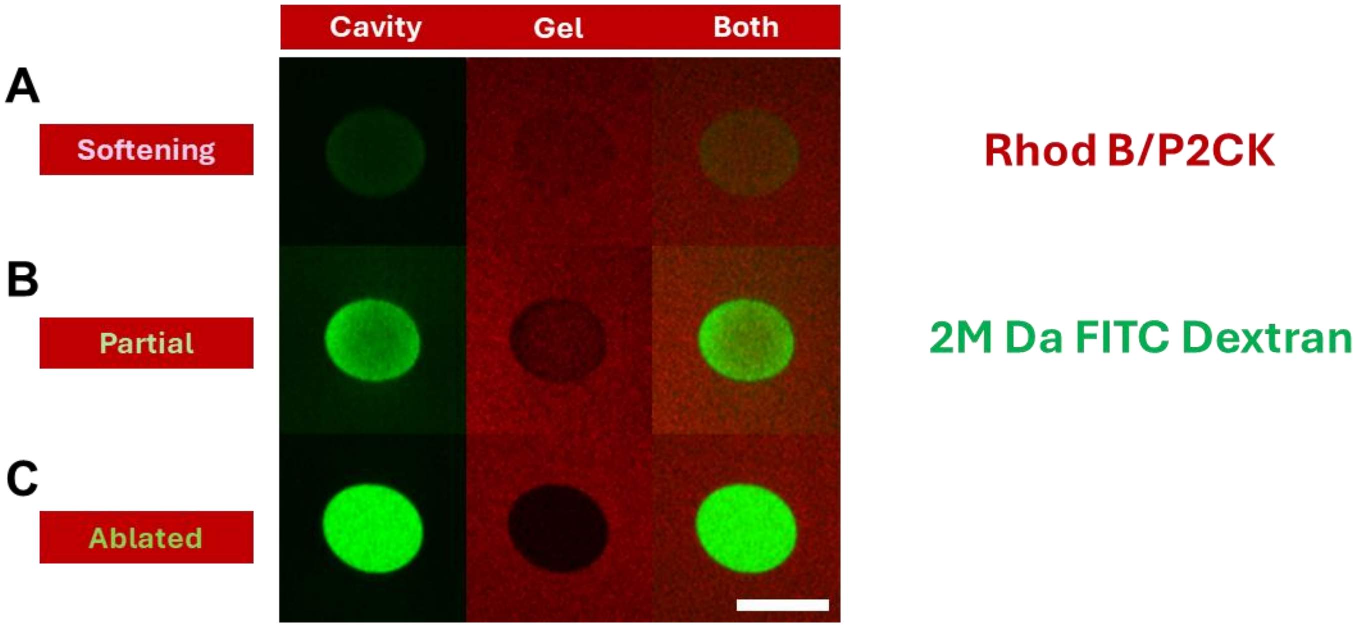


**Figure S10: Determination of ablation quality: All gels were crosslinked with rhodamine B acrylate and perfused with High MW (2M Da) FITC Dextran. Signal from remaining P2CK sensitizer was also measured in Rhod B channels.** (A) Slight increase in intensity of FITC channel and slight decrease of signal in Rhod B channel (B) Clearly defined border starts to appear in Rhod B channel, bright green signal at edges of ablated structures. (C) Bright signal from FITC channel and absence of Signal from Rhod B.

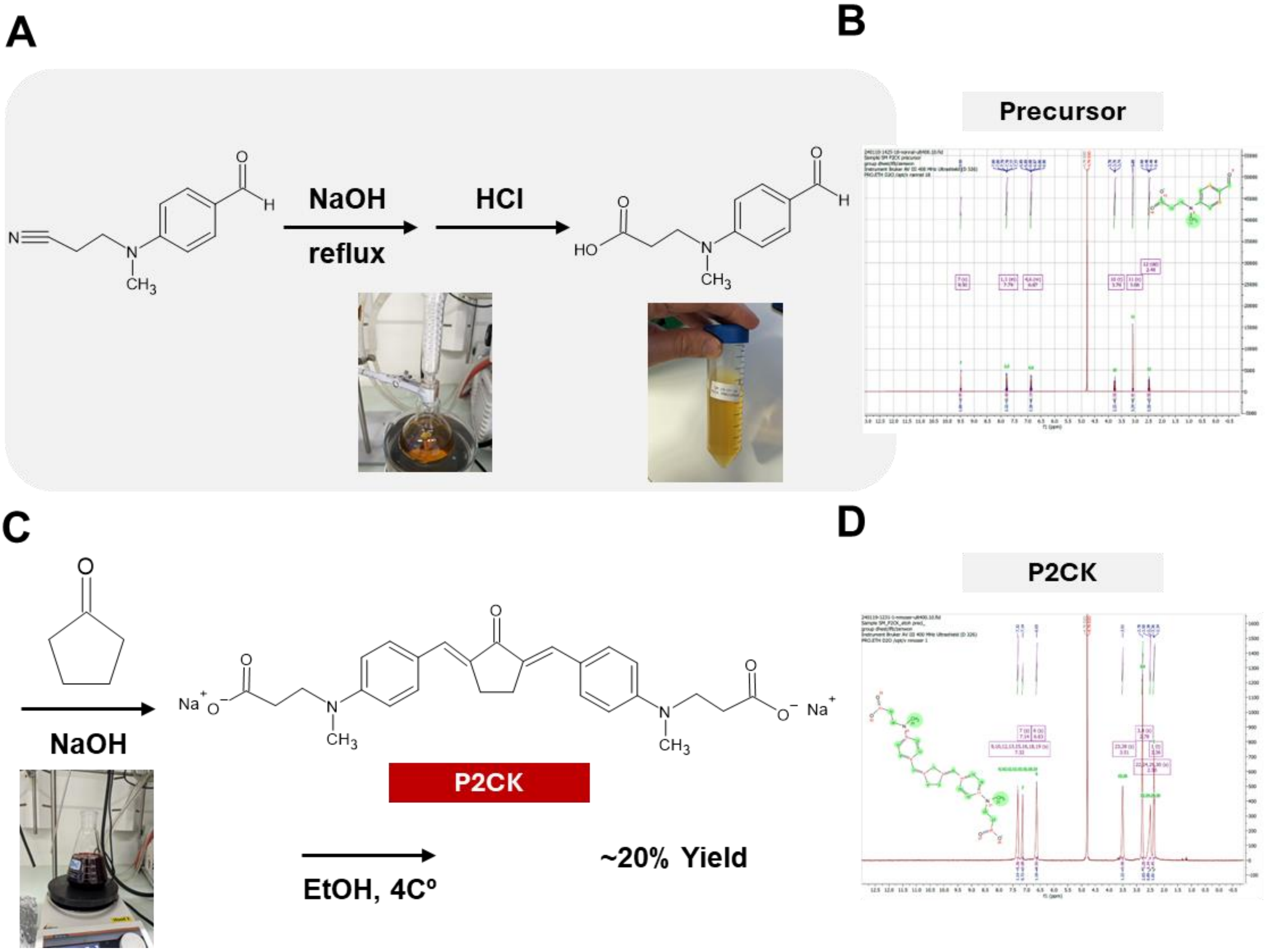


**Figure S11: Synthesis of 2PP Sensitizer P2CK** (A) In water 3-((-4-formylphenyl)-(methyl)-omin)propanetrile was reflux with the addition of NaOH. The product was then neutralized with HCl and freeze dried. (B) H-NMR of synthesized precursor. (C) Cyclopentanone was added to the dissolved precursor in NaOH solution and refluxed to obtain the sensitizer. P2CK was precipitated adding Ethanol on ice to get a yield of 20% (D) H-NMR of Synthesized P2CK.

### P2CK Synthesis

In 200ml DI water 9g of 3-((-4-formylphenyl)-(methyl)-omin)propanetrile (FPMOP, BLP Pharma BD8236) and 5.6g NaOH (SA) were mixed for 30min and ultrasonicated for 5min. The solution was refluxed at 600rpm, increasing temperature from 115ºC to 150ºC. Complete dissolution was observed after 1h30min and the solution was kept for 5h afterwards. HCl was added to the filtrate until precipitation formed. The precipitate was washed 3 times by centrifugation and decanting. pH was adjusted to 7.09 measured with a pH meter by adding 1M NaOH and HCl after which the product was freeze dried.

Of this precursor 0.574g (2.5mmol) were added together with 110µl cyclopentanone (2.5mmol, SA C112402-100ML) and 500ul 1M NaOH in 4.5ml DI water. The mixture was refluxed again for 5 h, increasing the temperature from 90ºC to 105ºC. After cooling down, 40ml EtOH, was added and stirred on ice for 1h until precipitation formed. Additionally, the supernatant was centrifuged at 2547 rpm at 0ºC for 1h. All precipitation was washed with EtOH twice. After decanting the powder was put into oven at 40ºC to let dry.[1,2]

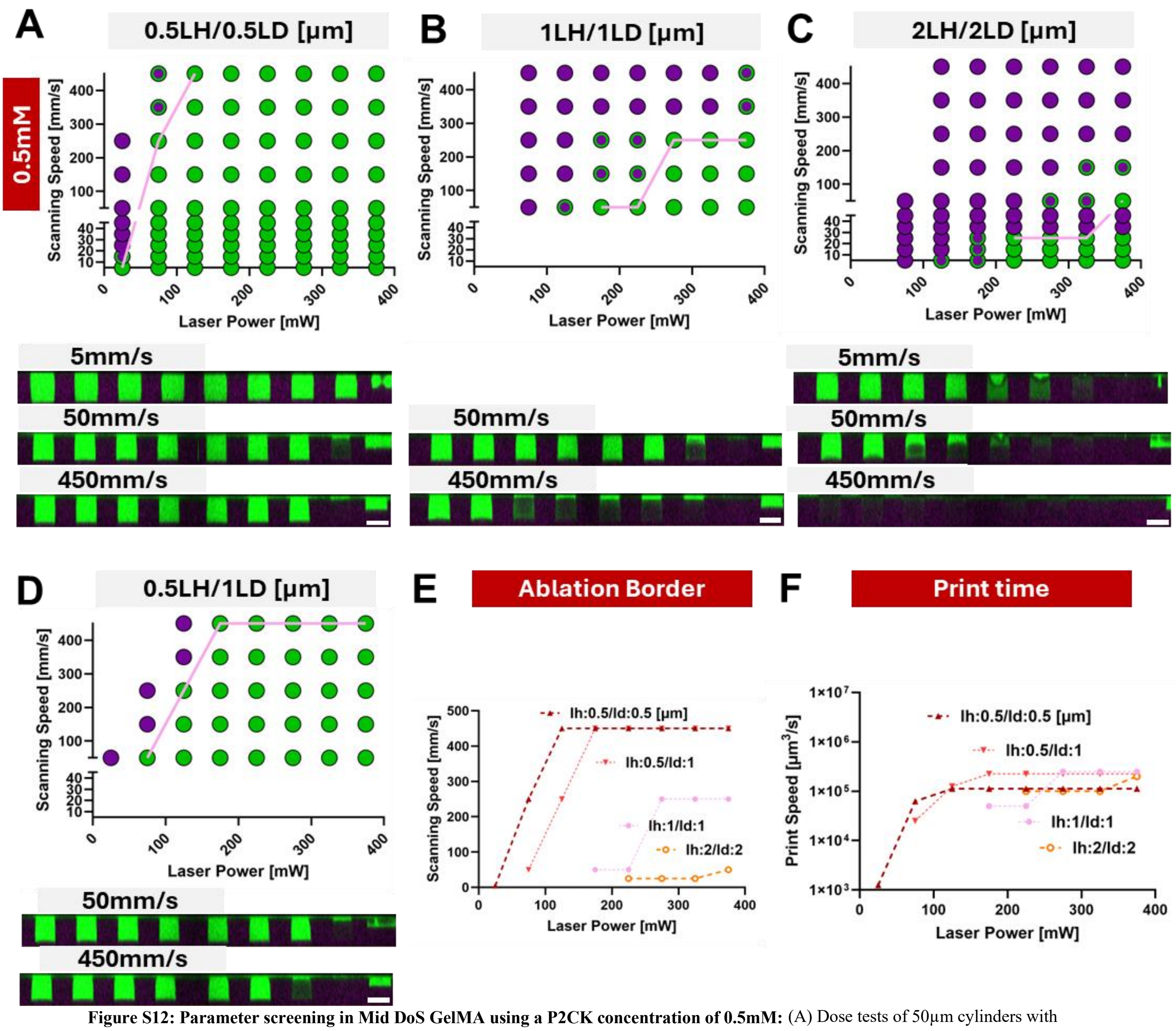


**Figure S12: Parameter screening in Mid DoS GelMA using a P2CK concentration of 0.5mM:** (A) Dose tests of 50µm cylinders with Laser Power varying from 25mW to 375mW and scanning speed from 5 to 450mm s-1 using a layer height LH of 0.5µm and a Line distance LD of 0.5µm (B) Dose test using a LH of 1µm and LD of 1µm. (C) Layer height and line distance are increased to 2µm. (D) LH of 0.5µm and LD of 1µm. (E) Border where ablation starts for the tested hatching distances. (F) calculated print time for the 4 conditions.

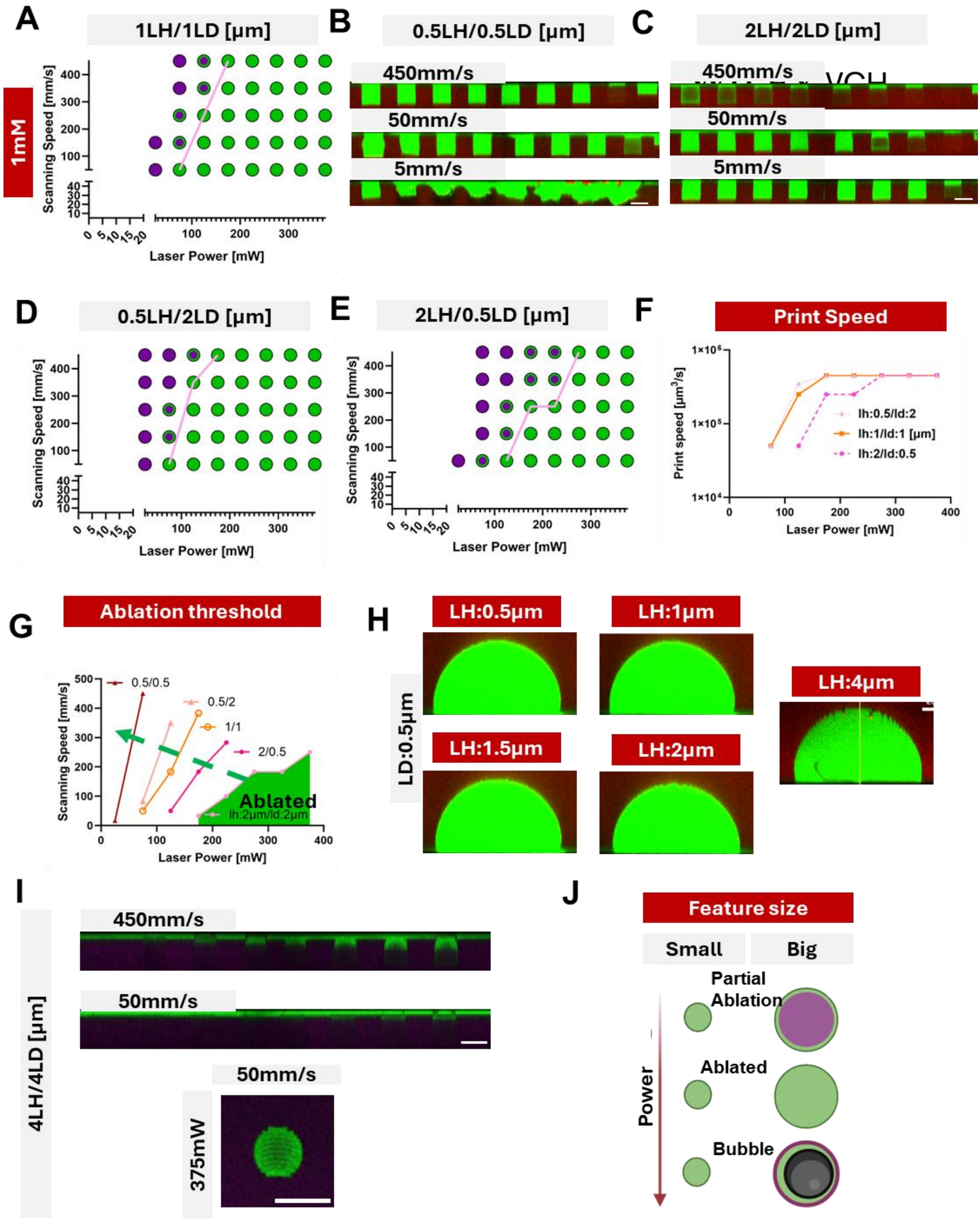


Figure S13: Parameter screening and effects of layer height and line distance on surface structure in Mid DoS GelMA using a P2CK concentration of 1mM. (A) Dose test overview for LH 1µm and LD 1µm Laser Power: 25mW-375mW scanning speed: 5-450mm s-1) (B) Side view of dose test from figure 3D for 5, 50 and 450mm s-1. (C) Side view of figure 3E, LH and LD are 2µm (D) Dose test with hatching distance LH 0.5µm and LD 2µm (E) Hatching distances flipped to LH 2µm and LD 0.5µm (F) Theoretical print speed for various hatching distances with the same total energy deposited (G) Ablation threshold border influenced by hatching distances. To the right of the line ablation occurs. (H) Effects of Layer height on Surface quality for a given line distance of 0.5µm. Layer height is increased from 0.5 to 2µm (1mM P2CK) and to 4µm (2mM P2CK). (I) Dose test outcome for LH 4µm and LD 4µm using 1mM P2CK. Top: Side view of 50 and 450mm s-1 dose test. Bottom: top view over condition exposed to highest Laser Power (375mW) and lowest scanning speed (50mm s-1). (J) Size effect in ablation with bigger structures needing more power for complete ablation while onset of cavitation bubble formation is at lower Powers.

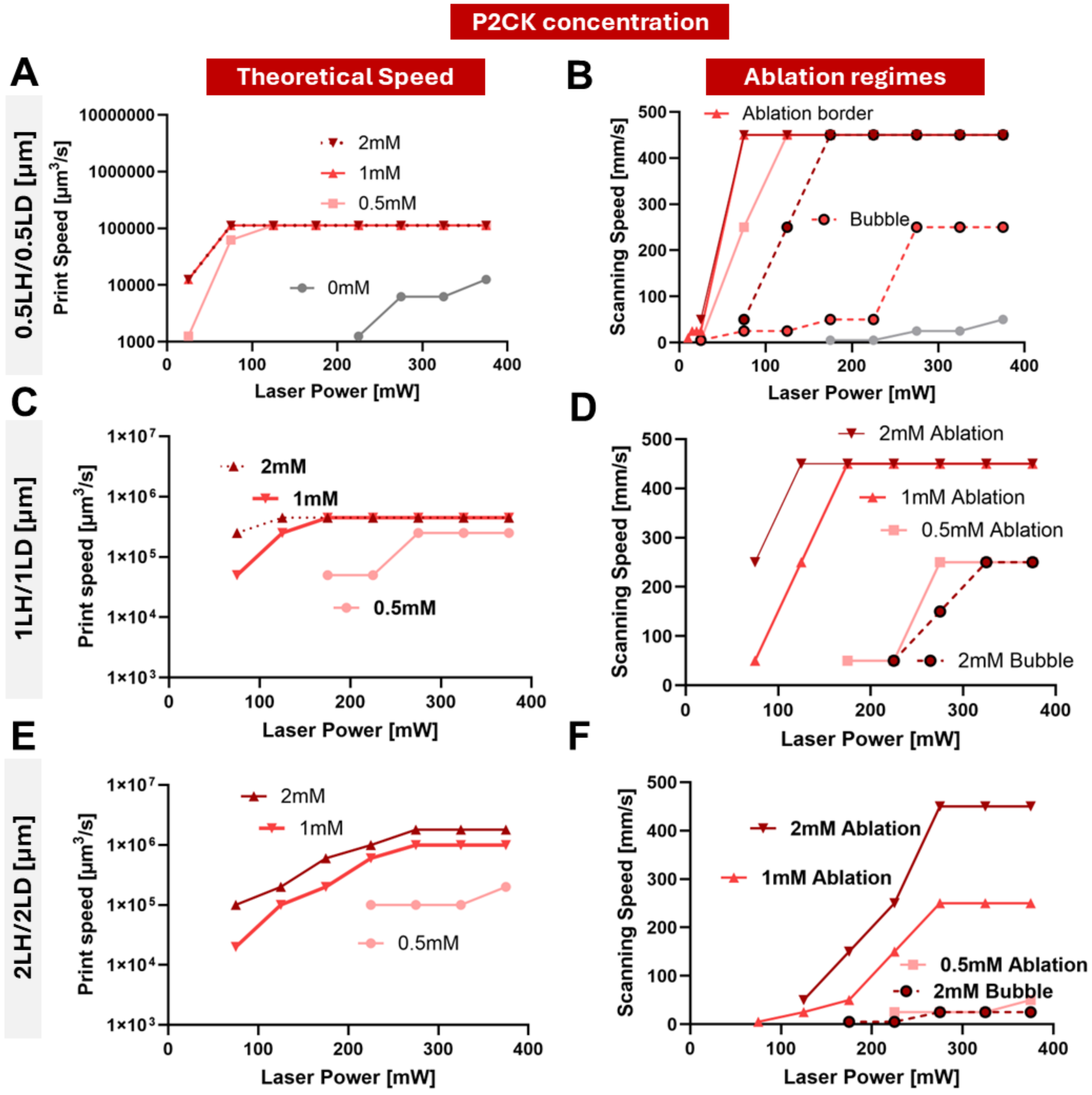


**Figure S14: Comparison of Sensitizer concentration effect on Theoretical print speed and ablation regimes for different LH and LD.** (A) Calculation of theoretical print speed as a function of Laser power for 0mM. 0.5mM, 1mM and 2mM for a hatching distance of 0.5µm. (B) Border of ablation regime and bubble formation (1mM and 2mM only) for the same concentrations. (C) Print speed for 0.5mM 1mM and 2mM and Ablation regime (D) for LH:1µm and LD:1µm and for LH&LD:2µm (E) & (F) n=3, Median is shown and used to calculate theoretical print speed

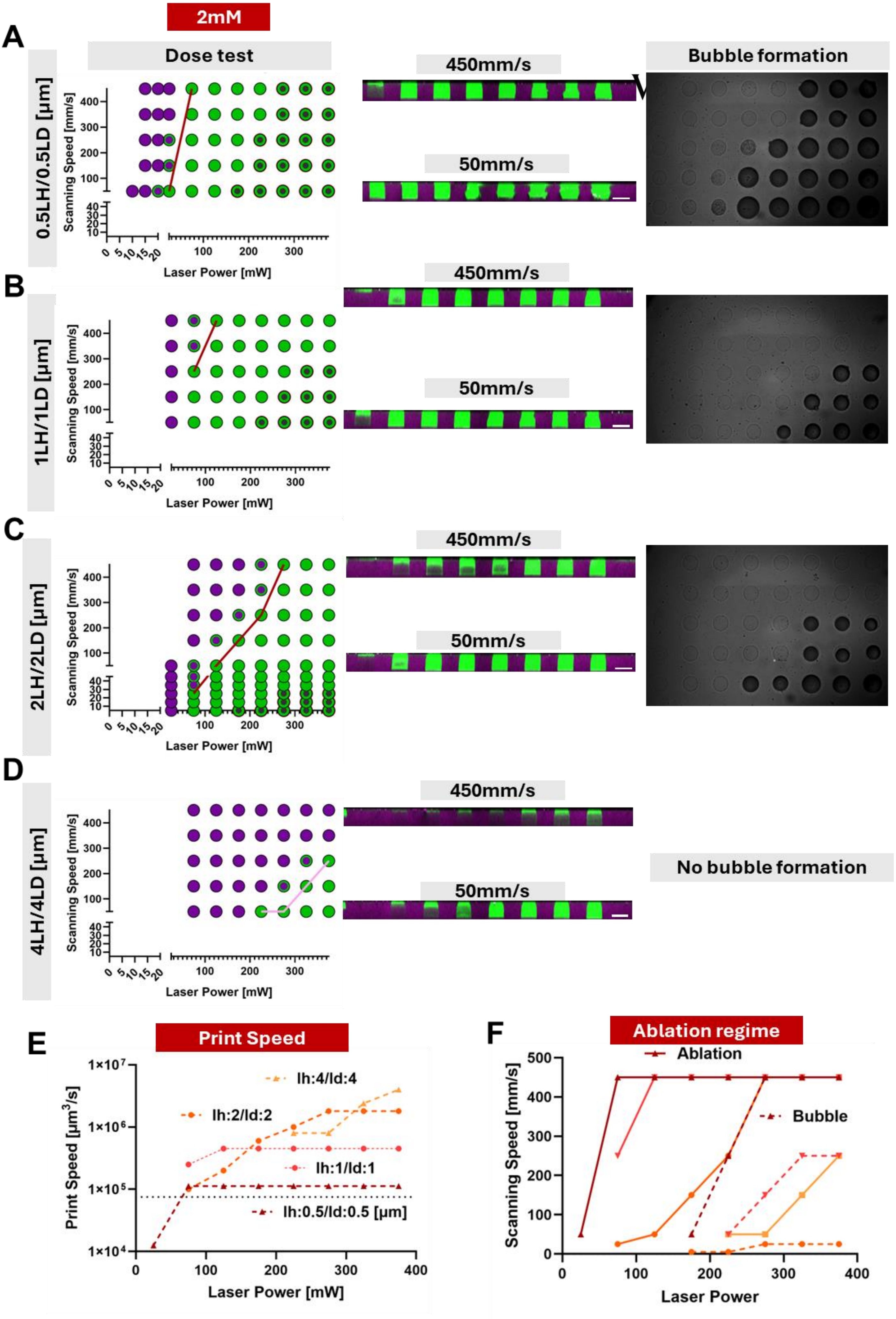

2mM
A
Dose test
0.5LH/0.5LD [µm]
Scanning Speed [mm/s]
Laser Power [mW]
450mm/s
50mm/s
Bubble formation
B
1LH/1LD [µm]
C
2LH/2LD [µm]
D
4LH/4LD [µm]
No bubble formation
E
Print Speed
Print Speed [µm³/s]
lh:4/ld:4
lh:2/ld:2
lh:1/ld:1
lh:0.5/ld:0.5 [µm]
F
Ablation regime
Ablation
Bubble
Scanning Speed [mm/s]
Laser Power

**Figure S15: Parameter screening using 2mM P2CK.** (A) Dose test overview (left) side view with increasing Power from 25mW to375mW for 50 and 450 mm s-1 (middle), bubble formation during printing (right panel) for 0.5LH/0.5LD. (B) 1LH/1LD (C) 2LH/2LD (D) 4lH/4LD, no bubble formation was observed (E) Print Speed and (F) Ablation/Bubble formation border for the tested hatching distances.

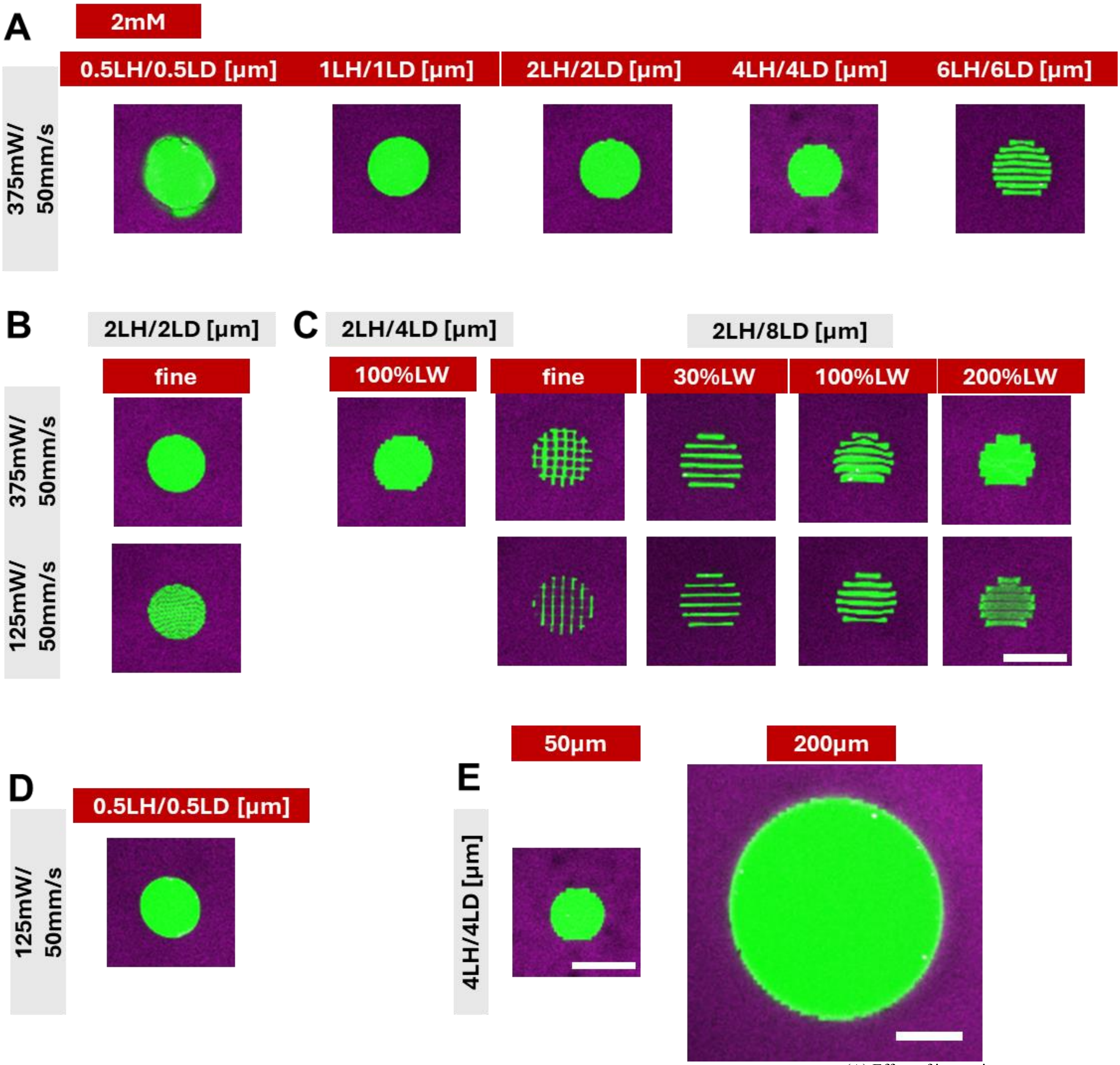


**Figure S16: Effect of Hatching distance on ablation and surface quality using a P2CK concentration of 2mM.** (A) Effect of increasing hatching distance (ltr: 0.5, 1, 2, 4, 6 µm). (B) fine mode for a hatching distance of 2 µm using high power (top row) and lower power (bottom row). (C) Effect of line width increasing from fine mode to 30% (of LD), 100% and 200% (LH:2µm LD:8µm) for two conditions (top row: 375mW, bottom row: 125mW). (D) Improved parameters (lower power) for hatching distance of 0.5µm. (E) Size effect on surface quality for 50 and 200µm diameter.

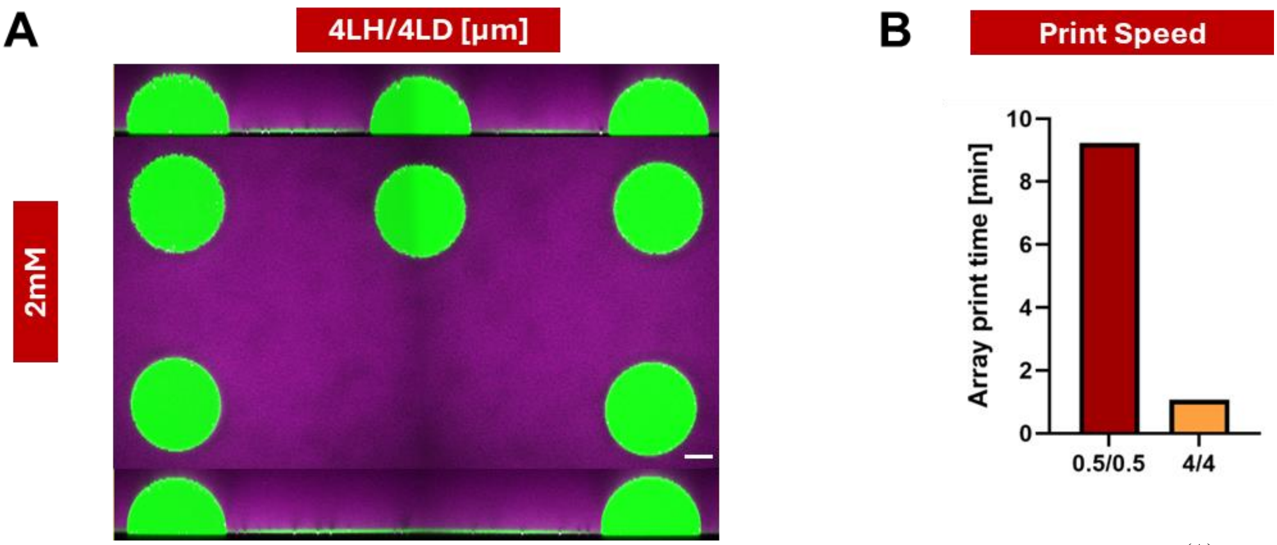


**Figure S17: Increase of actual print time by using higher hatching distance in combination with higher P2CK concentration** (A) Surface roughness array of five 200µm half spheres with varying surface roughness using 2mM P2CK. (B) Actual Print time using a hatching distance of 0.5µm vs a hatching distance of 4µm

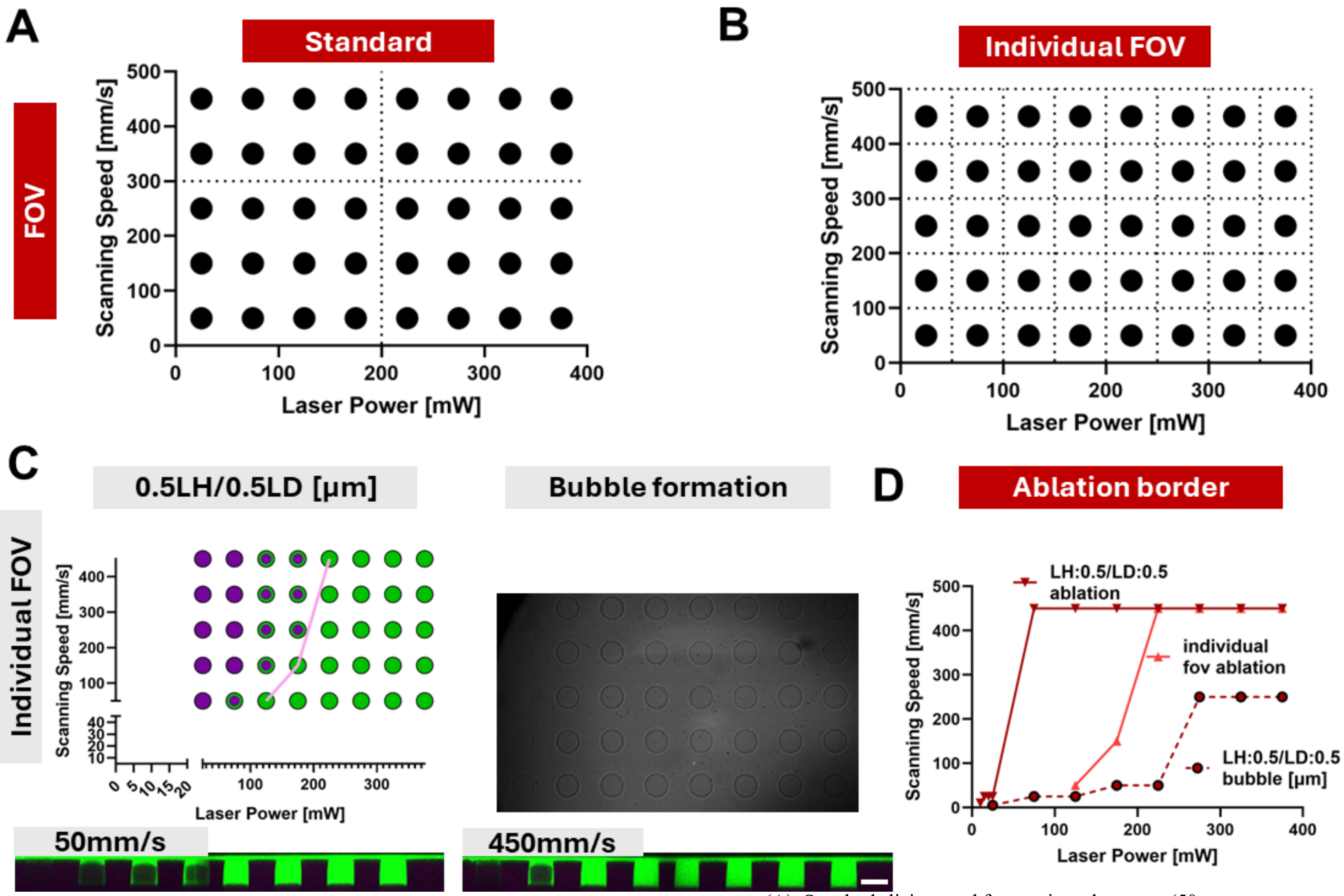


**Figure S18: Effect of Field of view (FOV) on ablation and bubble formation:** (A) Standard slicing used for previous dose tests (50µm cylinder) (B) Dose test with each cylinder located in an individual FOV (C) 0.5µm dose test overview (left) using individual FOV, absent bubble formation during printing (right), side view of ablated dose test cylinders for 50 and 450mm s-1. (D) Ablation and bubble formation border compared to standard FOV.

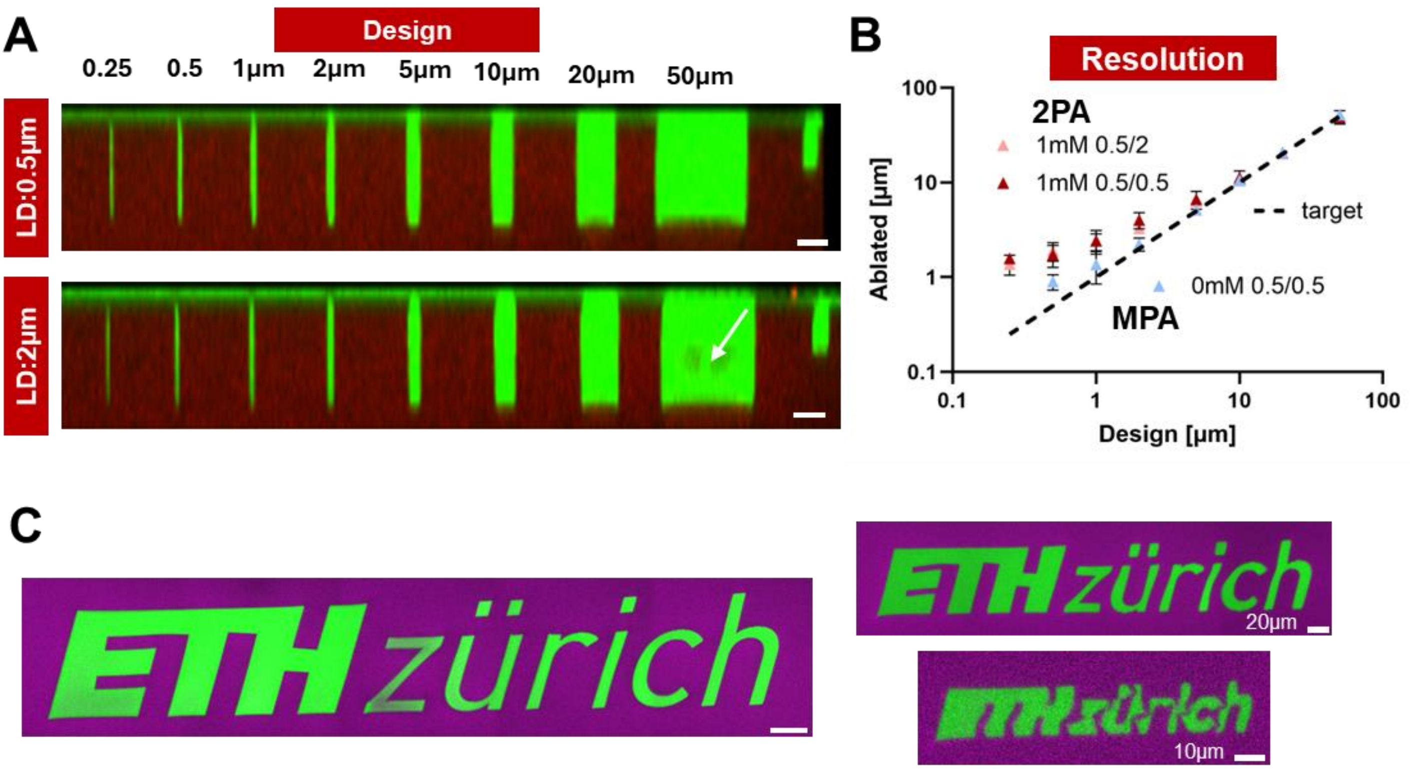


**Figure S19: MPA Resolution in MiD DoS GelMA with 1mM P2CK Sensitizer.** (A) Resolution test of cylinder with design diameter indicated on top (0.25 0.5 1 2 5 10 20 50µm ltr). Top row: LH: 0.5µm LD: 0.5µm; Bottom row: LH: 0.5µm LD: 2µm. (B) Measured diametr versus the design diameter of the channel for both conditions and LH: 0.5µm LD: 0.5µm without Sensitizer (0mM). (C) ETH Logo of varying size fabricated with MPA.

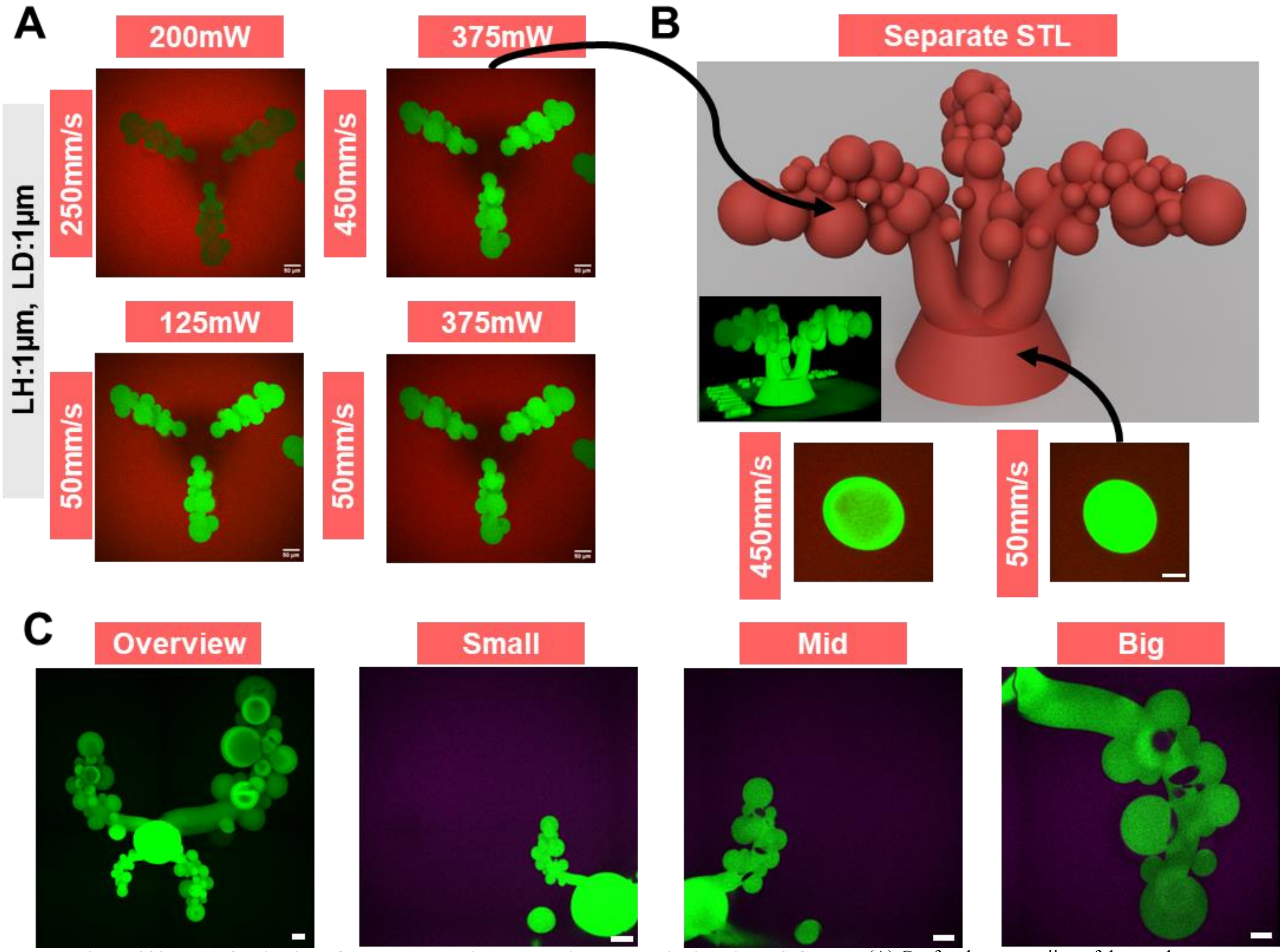


**Figure S20: MPA fabrication of complex alveoli plus duct like geometries in Mid DoS GelMA.** (A) Confocal crosssectiion of duct and alveoli model using a hatching distance of 1µm (top left) 200mW 250mm s-1 (top right) 375mW 450mm s-1 (bottom left) 125mW 50mm s-1 (bottom right) 375mW 50mm s-1 (B) Top row: Render of stl file and 3d view of ablated structure (inset). Bottom row: base printed using 375mW and 450mm s-1 (left) and 50mm s-1 (right). (C) Varying size of duct + alveoli geometries (ltr) overview, small, medium size and largest structure.

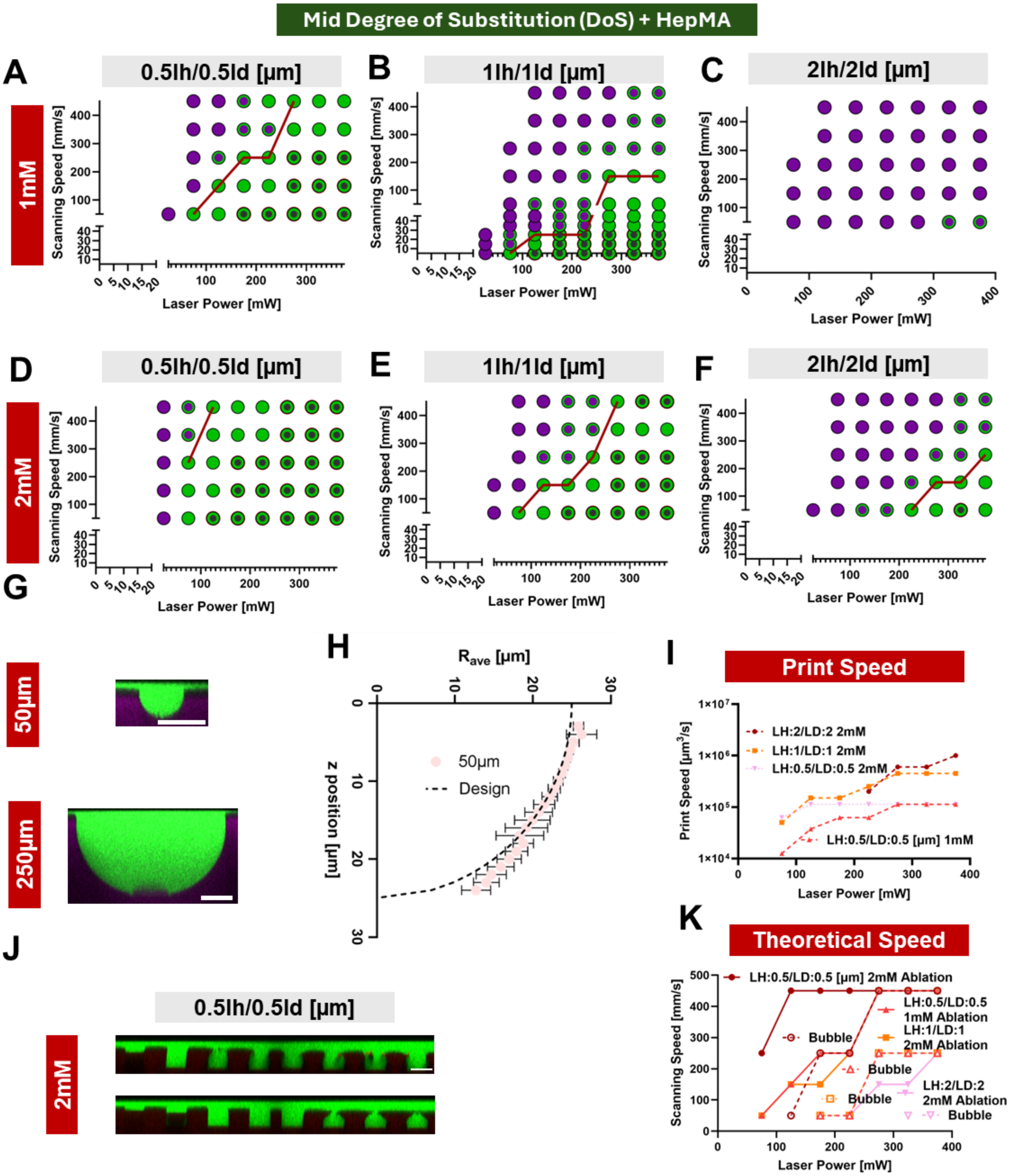


**Figure S21: Parameter screening and ablation in Mid DoS GelMA + HepMA using 1-2mM P2CK sensitizer.** (A) Dose test using 1mM P2CK with hatching distance of 0.5µm (B) 1µm (C) 2µm and 2mM ((D) 0.5µm (E) 1µm (F) 2µm) (G) Representiative confocal xz crosssection image of 50µm and 250µm ablated half sphere using MPA with 2mM P2CK. (H) Variability of Average radius of an ablated 50µm half sphere as a function of z position. (I) Theoretical print speed as a function of Lasert calculated for 0.5um hatching distance (1mM, 2mM P2CK), 1um and 2um (2mM P2CK). (J) Example crosssection of dose test with parameters used for ablating geometries (hatching distance 0.5, 2mM P2CK). (K) Ablation regime and bubble formation for all investigated hatching distances using 2mM P2CK.

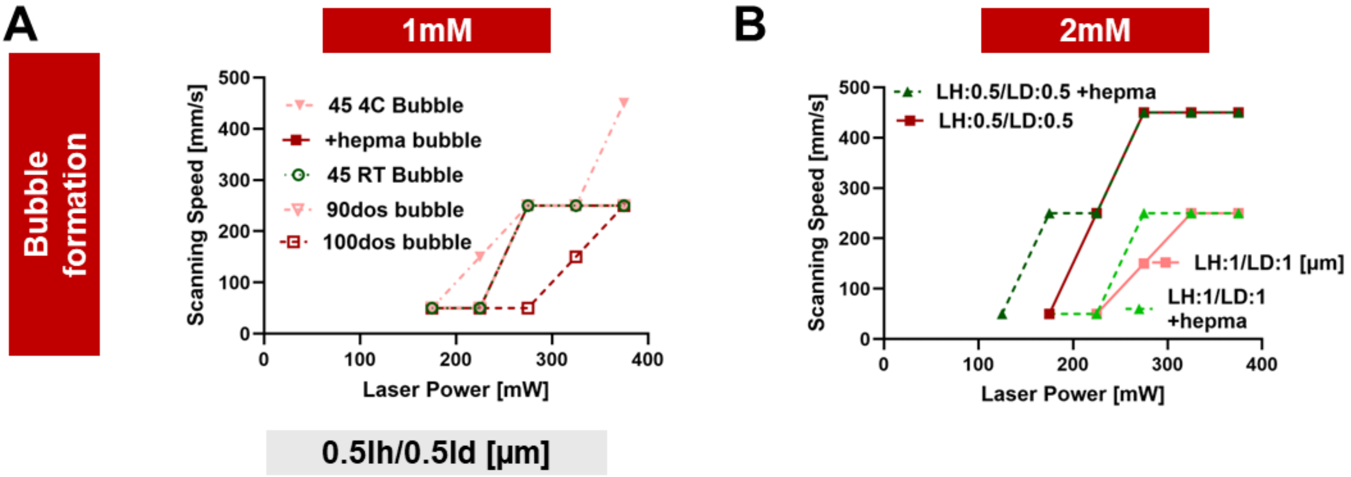


**Figure S22: Bubble formation regimes for various material systems and hatching distances.** (A) Using a hatching distance of 0.5µm and 1mM P2CK bubble formation onset is shown for Low DoS GelMA(4C and RT precrosslinked), Mid DoS, Mid DoS + HepMA and High DoS. (B) Bubble formation for increased P2CK concentration (2mM) for Mid DoS and Mid DoS + HepMA.

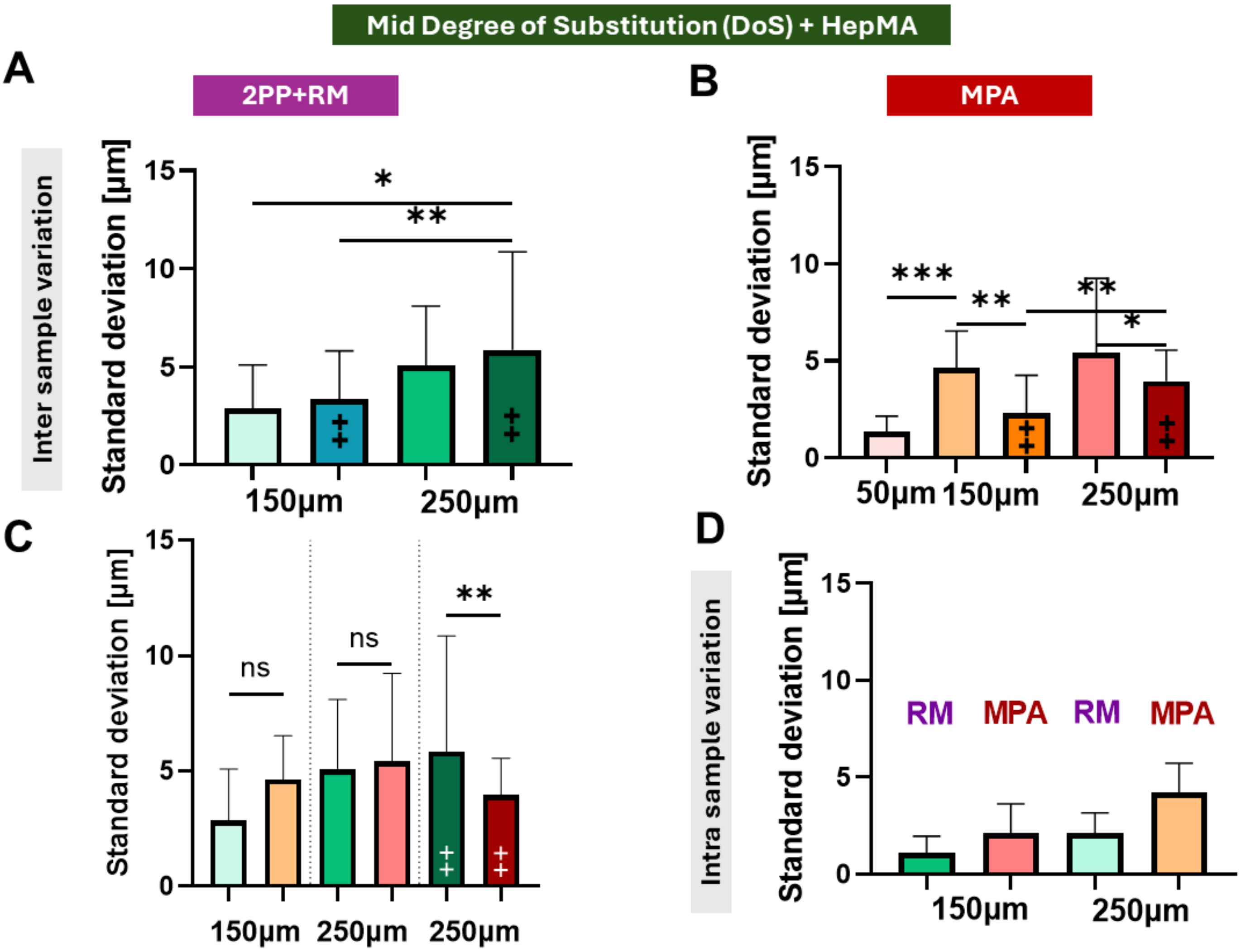


**Figure S23: Comparison between variability between 2PP+RM and MPA fabricated curvatures in Mid DoS+ HepMA hydrogels.** (A) Inter sample variability (averaged over z position) of 150 and 250µm half spheres and ellipsoids (++) fabricated with 2PP+RM (B) Variability of same curvatures fabricated with MPA, in addition variability of 50µm half sphere is shown. (C) Direct comparison of 2PP+RM and MPA for 150µm, 250µm and 250µm++ (D) Intra sample variation of 150 and 250µm half spheres fabricated with 2PP+RM and MPA.

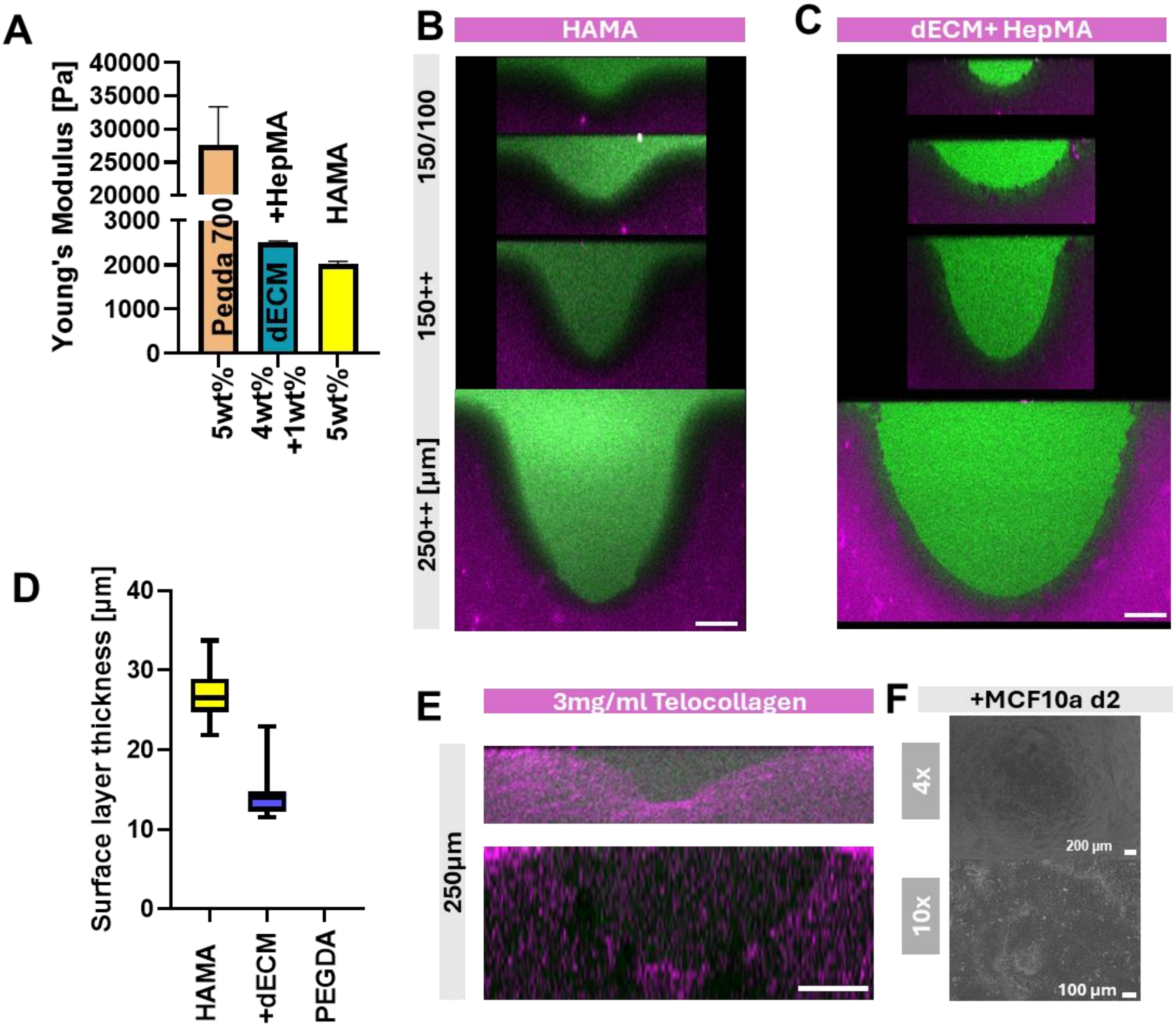


**Figure S24: Fabrication of curvatures using different material systems with 2PP+RM.** (A) Young's Modulus measured by compression test for PEGDA 700, dECM+HepMA and HAMA hydrogels (B) Array of 100 150 150++ 250++ µm ellipsoids in HAMA replicated from PDMS molds (C) Same curvatures in dECM+HepMA (D) Soft surface layer thickness measured for HAMA, HepMA + dECM and PEGDA (E) Curvatures fabricated with soft and viscoelastic telocollagen (Adcanved Biomatrix, 3mg/ml). (F) MCF10a cells attaching on 3mg/ml Telocollagen gels.

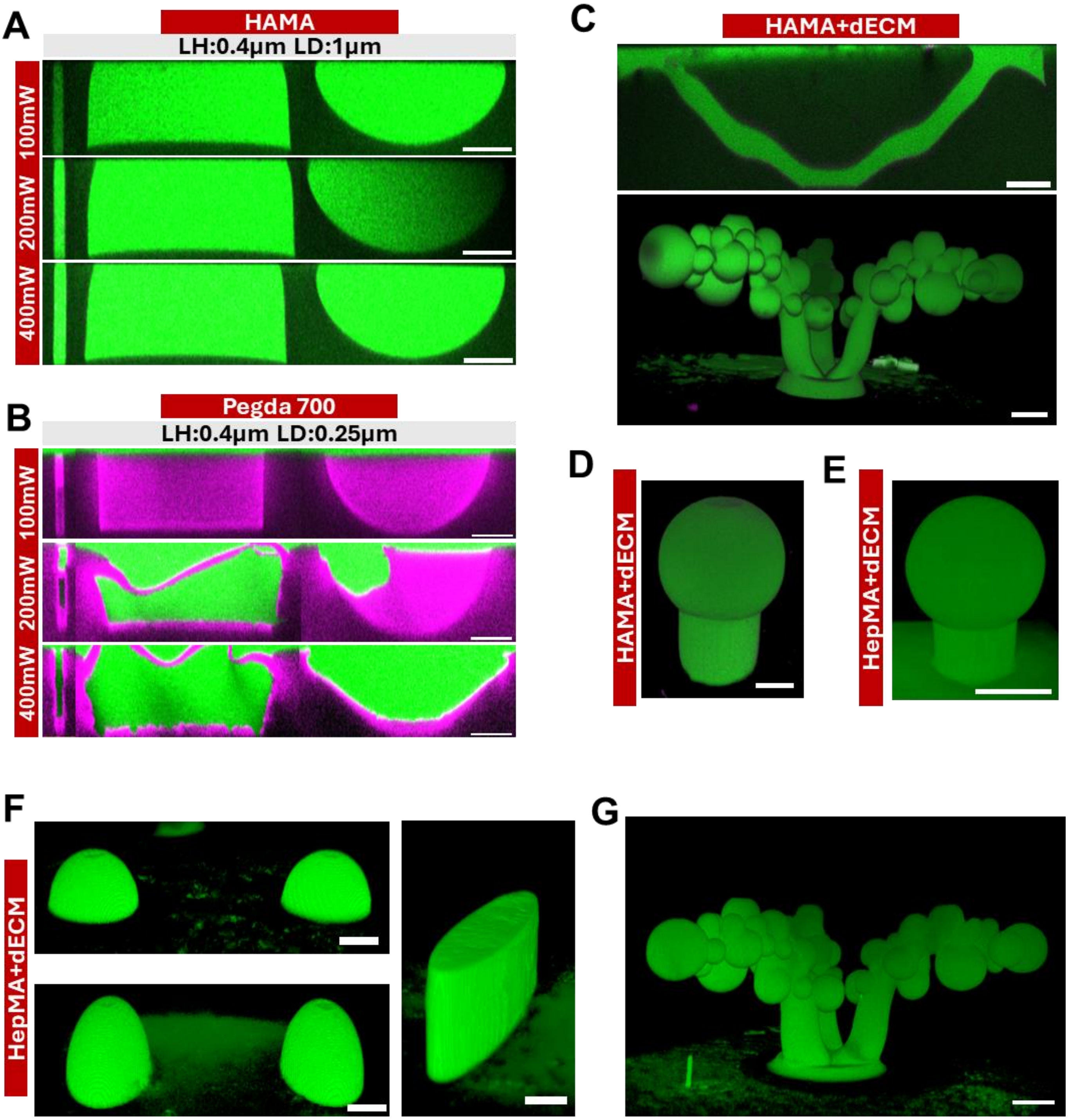


**Figure S25: Curvatures fabricated with MPA in various material systems.** (A) Crosssection of 10µm and 200µm half spheres and 200µm cylinder 100mW (top), 200mW (mid), 400mW (bottom). (LH:0.4µm LD:1µm Scanning speed 400mm s-1 (B) Same structures in PEGDA 700 hydrogels. 100mW (top), 200mW (mid), 400mW (bottom). (LH:0.4µm LD:0.25µm Scanning speed 400mm s-1 ). (C) Channel and duct/alveoli geometries ablated in HAMA with the addition of dECM (D) Combination of spherical (top) and cylindrical(bottom) geometry ablated in HAMA+dECM and (E) HepMA+dECM. (F) Concave Gaussian curvatures (left, 150µm) and elliptical cylinder ablated in HepMA + dECM. (G) Duct+Alveoli geometry in HepMA+ dECM.

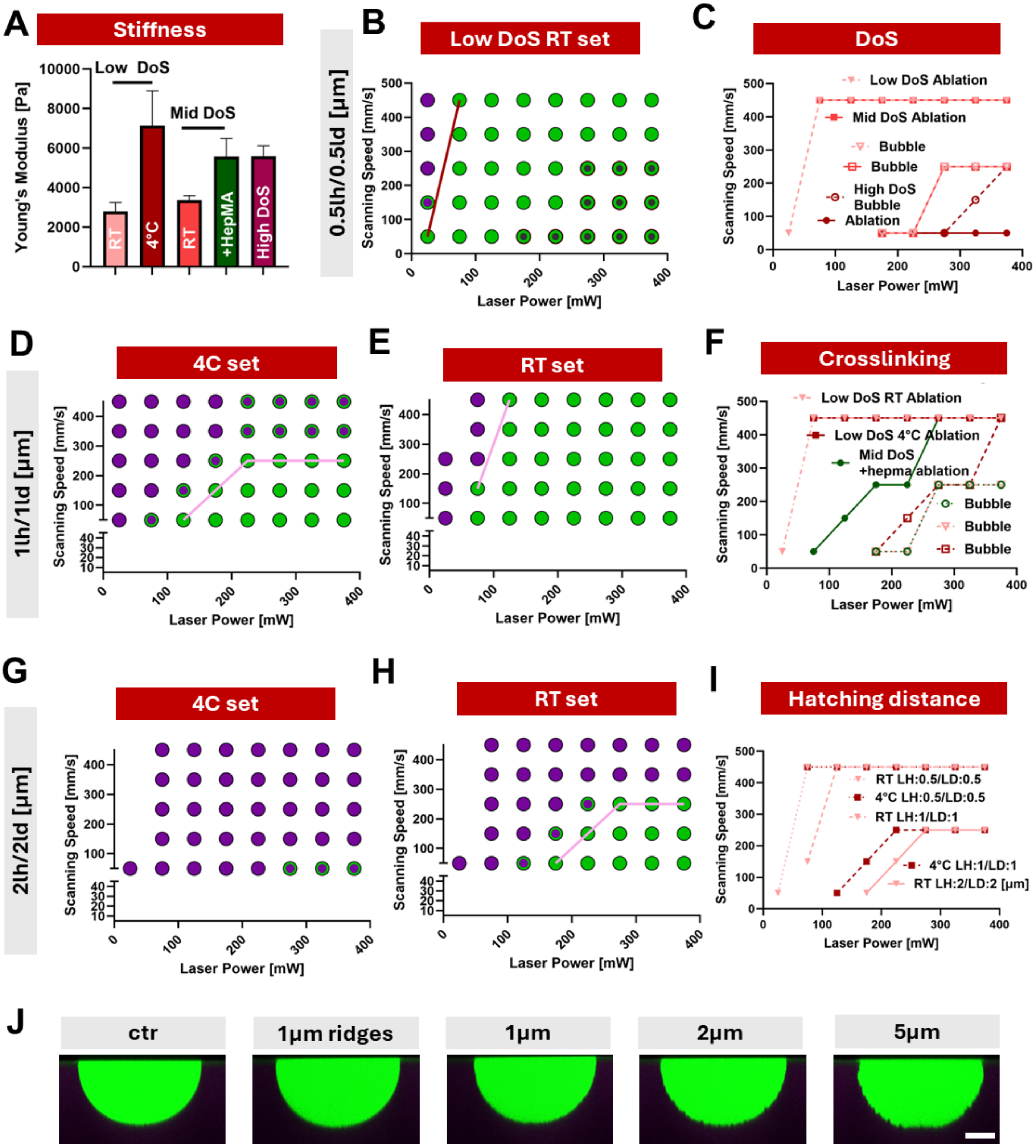


**Figure S26: Influence of thermal precrosslinking on ablation regimes.** (A) Youngs Modulus (compression test) of various degrees of substitution GelMA and with the addition of HepMA (B) Dose test in Low DoS GelMA precrosslinked at RT with 1mM P2CK using a hatching distance of 0.5µm. (C) Ablation and bubble formation regimes for different degree of substitution (D) Dose test with hatching distance of 1µm crosslinked at 4ºC (E) and at RT. (F) Effect of crosslinking mechanism (effect of RT vs 4ºC and addition of HepMA) on ablation and bubble formation. (G) Dose test for higher hatching distance 2µm in 4ºC set gels (H) and RT set gels (I) Influence of hatching distance on ablation regime depending on the crosslinking temperature for low DoS GelMA (J) Varying curvatures ablated in Low DoS GelMA using 1mM P2CK.

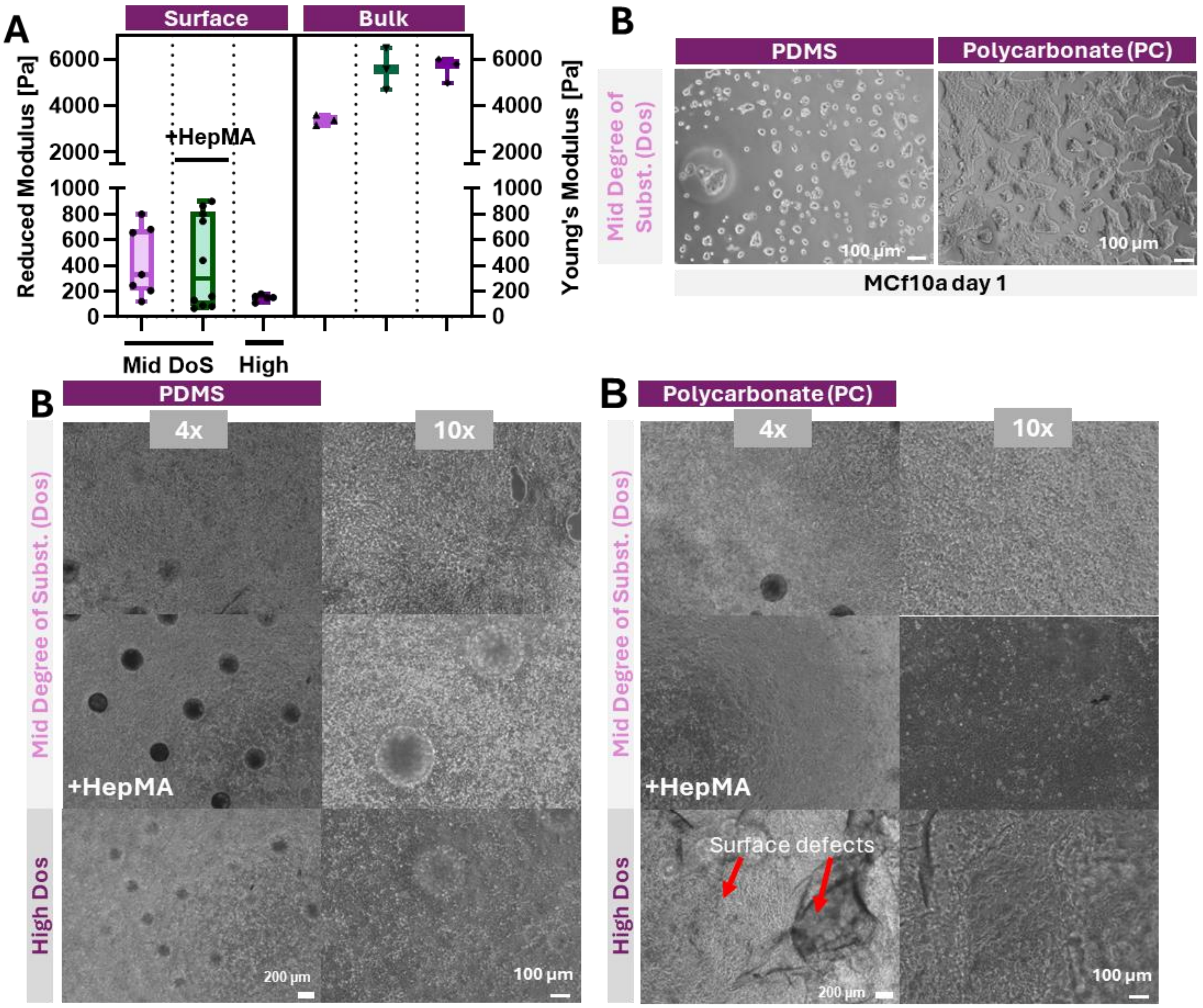


**Figure S27: MCF10a cell layer after twelve days of culture on thermally crosslinked (RT) GelMA hydrogels.** (A) Hydrogels molded against PDMS. From top to bottom: Mid DoS, Mid DoS+HepMA, High DoS. (B) Same hydrogels molded against Polycarbonate (PC). Surface defects introduced by molding against PC are indicated with red arrows.

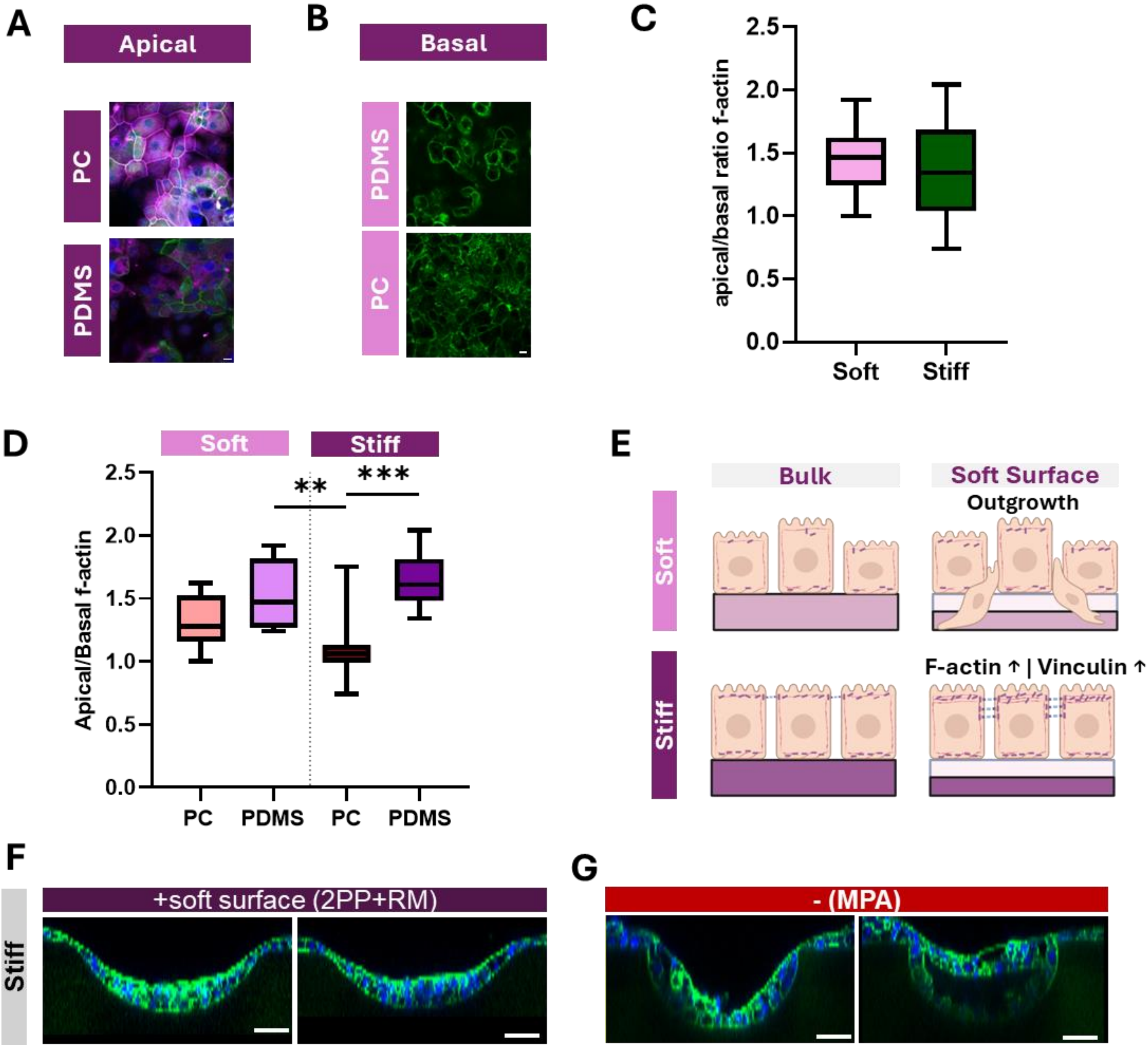


**Figure S28: Mammary epithelial cell layer polarization depending on substrate properties.** (A) MCF10a stained with Hoechst (blue), Phalloidin (green) and Vinculin (magenta). Apical view over cell layer on Mid DoS+HepMA molded against PDMS, and molded against PC. (B) Basal surface of MCF10a cells on soft hydrogels molded against PDMS (top and PC(bottom) (C) Pooled ratio of apical vs basal f-actin intensity for soft and stiff hydrogels including both samples molded against PDMS and PC. (D) Apical vs basal f-actin ratio for the individual investigated conditions. (E) Mechanism of polarization in depending on substrate stiffness and surface mechanical properties. (F) Representative images of MCF10a cells after d12 of cell culture on 250µm half spheres produced with stiff Mid DoS + HepMA gels. Left: representative structures fabricated with 2PP+RM including a soft surface layer; Right: representative structures fabricated with MPA.

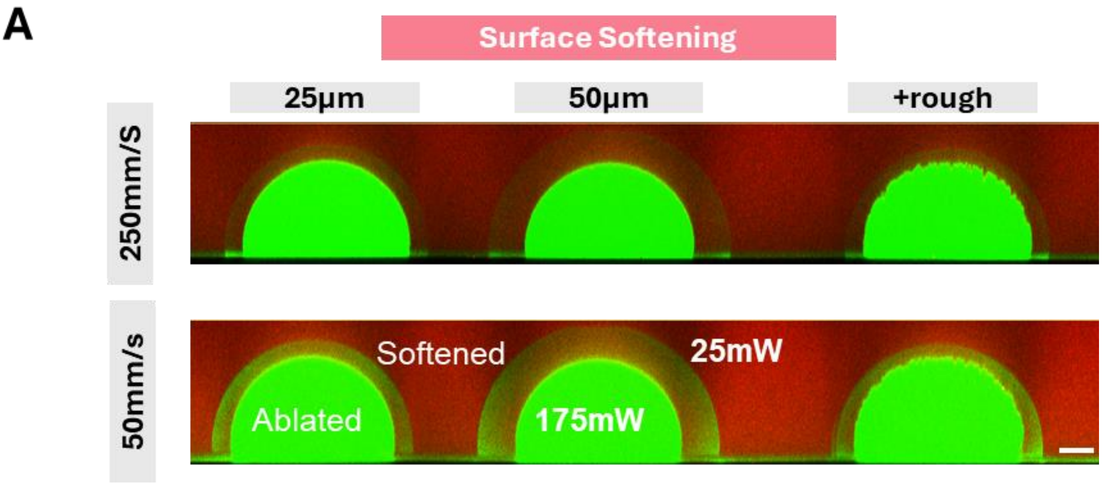


**Figure S29: Surface softening in Mid DoS GelMA.** (A) A 250µm half spheres ablated with 175mW (LH:0.5µm, LD:0.5µm) with a smooth surface (left, mid) and a rough surface (+/-2.5µm, right). A 300µm (left,right) and 350µm (mid) half sphere was softened around the ablated half sphere using a lower laser power of 25mW. Top row: high scanning speed (250mm/s) with less softening compared to lower scanning speed (50mm/s, bottom row).

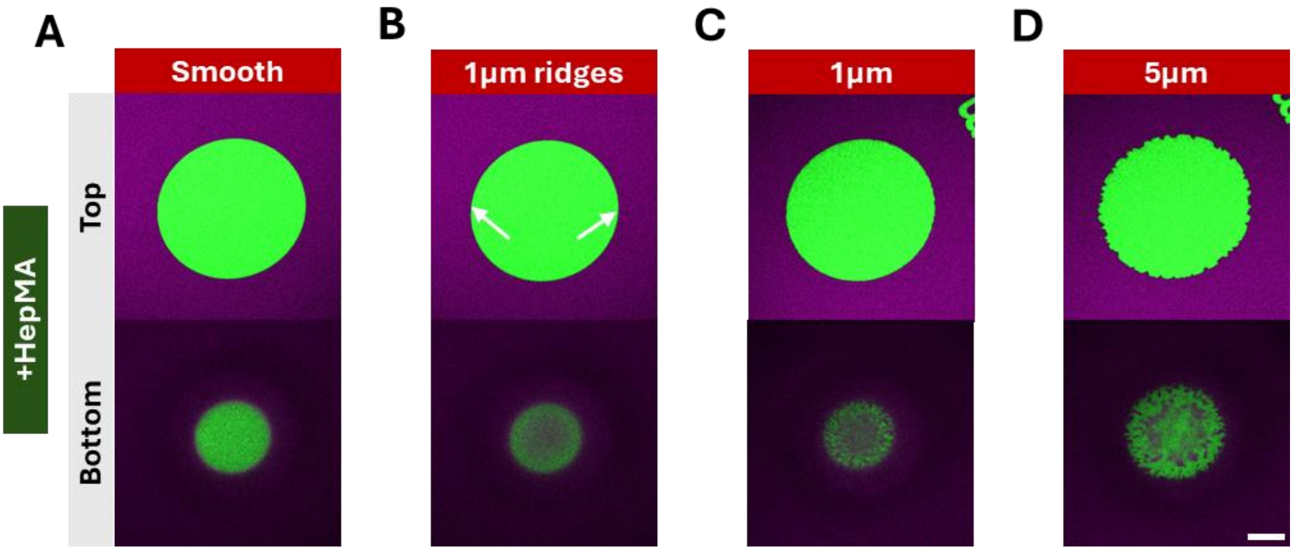


**Figure S30: Surface roughness introduced in Mid DoS + HepMA hydrogels.** (A) Smooth surface half sphere (200µm) (B) Ridges of 1um depth (indicated with white arrows) (C) random roughness of +/- 1µm and (D) +/- 5µm. A Sensitizer concentration of 2mM was used.

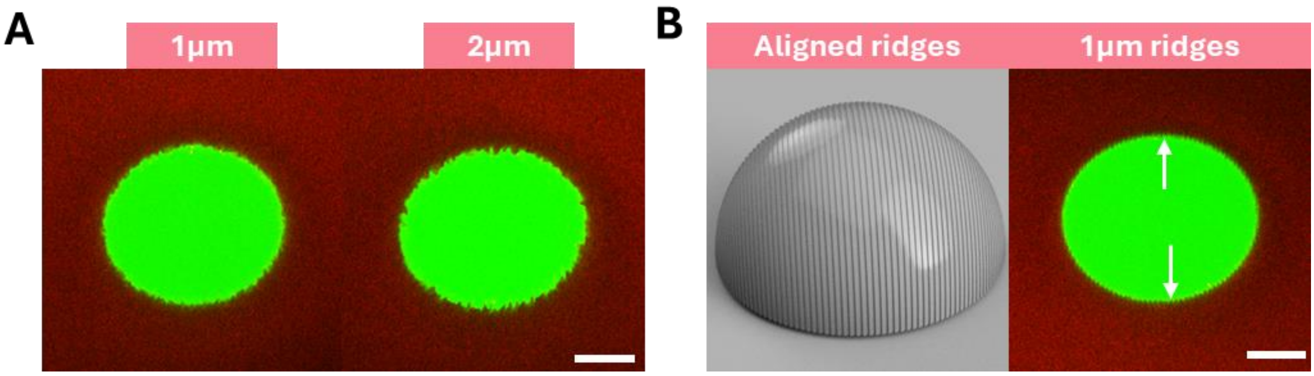


**Figure S31: Surface roughness introduced in MiD DoS GelMA.** (A) A 200µm half sphere with a surface roughness of +/- 1µm (left) and +/-2µm (right). (B) Design of 1µm ridges and ablated structure, white arrows indicate ridges.

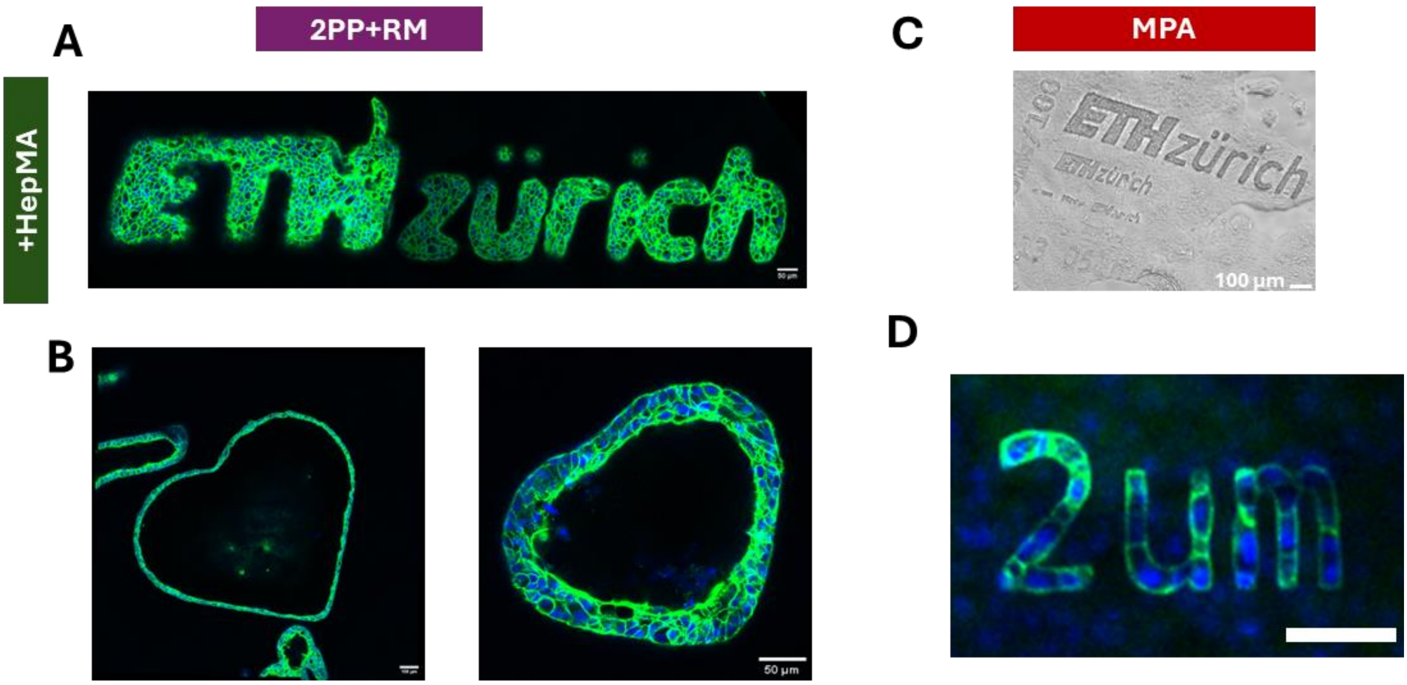


**Figure S32: MCF10a cells after twelve days on Mid DoS + HepMA hydrogels fabricated with both techniques.** (A) Mammary epithelial cells on medium sized ETH Logo fabricated with 2PP+RM. Cells are stained with Hoechst (Blue) and Phalloidin (green). (B) MCF10a cells on medium and small sized heart. (C) Brightfield image of MCF10a cells on ETH Logos of various sizes fabricated with MPA. (D) MCF10a cells on small description text fabricated with MPA.

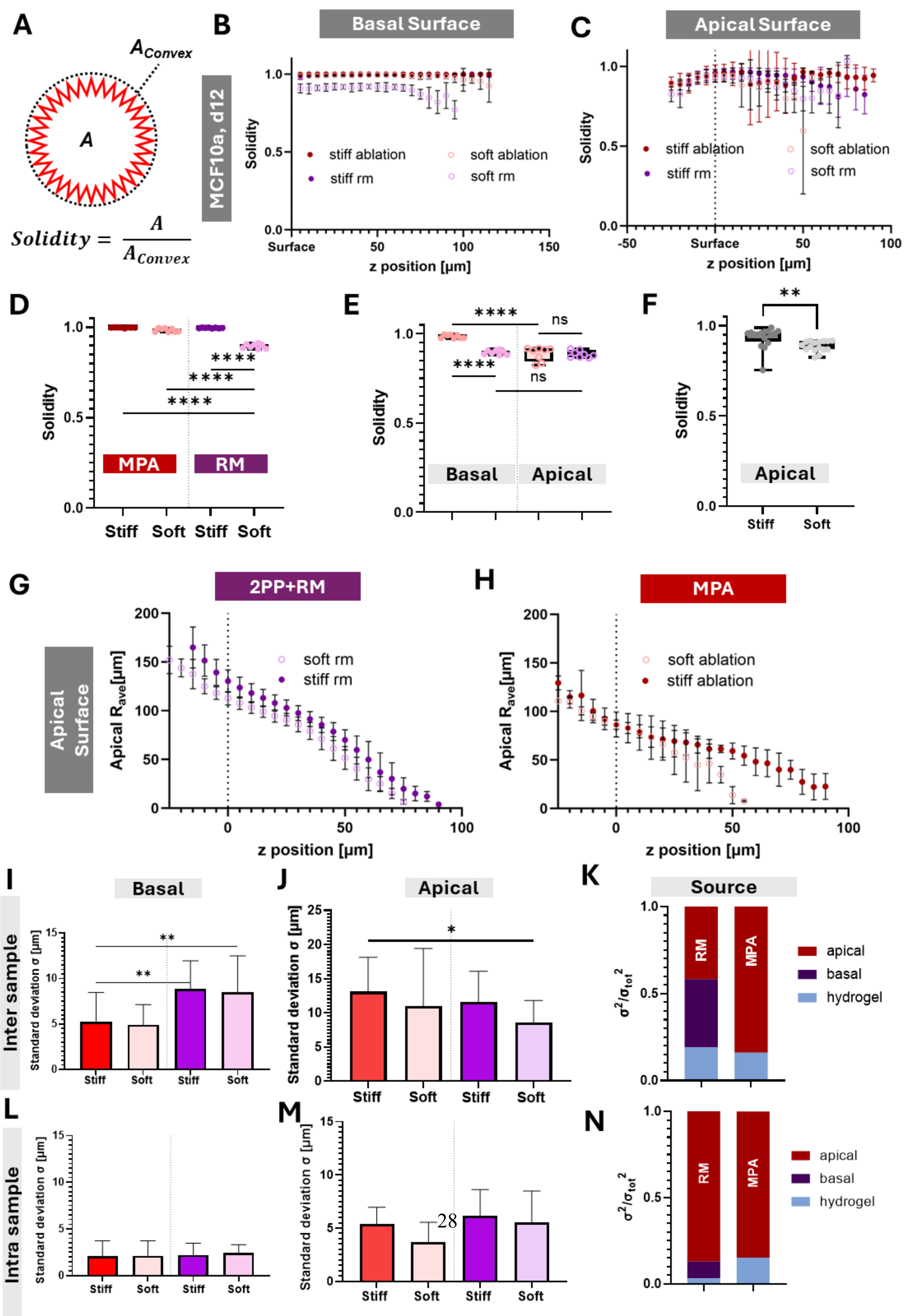
A
$A_{Convex}$
A
$Solidity = \frac{A}{A_{Convex}}$
MCF10a, d12
B
Basal Surface
Solidity
stiff ablation
soft ablation
stiff rm
soft rm
z position [µm]
C
Apical Surface
D
MPA
RM
Stiff
Soft
E
Basal
Apical
ns
F
G
2PP+RM
Apical Surface
Apical $R_{ave}$[µm]
H
MPA
I
Basal
Inter sample
Standard deviation σ [µm]
J
Apical
K
Source
apical
basal
hydrogel
$\sigma^2/\sigma_{tot}^2$
L
Intra sample
M
N

**Figure S33: Solidity and Variance of cell surfaces (apical and basal) after twelve days of culture for soft (Mid DoS) and stiff hydrogels (Mid DoS + HepMA).** (A) Calculation of Solidity as the ratio of area over the convex area. (B) Solidity of basal surface (towards substrate) over z position for soft and stiff hydrogels fabricated both with 2PP+RM and MPA (C) Solidity of apical surfaces (towards lumen) over z position (D) Average solidity for soft and stiff hydrogels fabricated with both techniques on apical side and (E) basal side. (F) Comparison of pooled solidity for stiff vs soft hydrogels including both techniques. (G) Average size of lumen/apical surface for 250µm half spheres fabricated with 2PP+RM and (H) MPA. (I) Inter sample variations for all tested conditions and fabrication techniques on the basal surface and (J) apical surface. (K) Relative share of the total standard deviation resulting from hydrogel fabrication, on the basal side and apical side for both 2PP+RM and MPA. (L) Intra sample variation on basal surface, (M) apical surface and relative share of standard deviation coming from hydrogel fabrication, basal surface and apical surface. n>3, w=2.

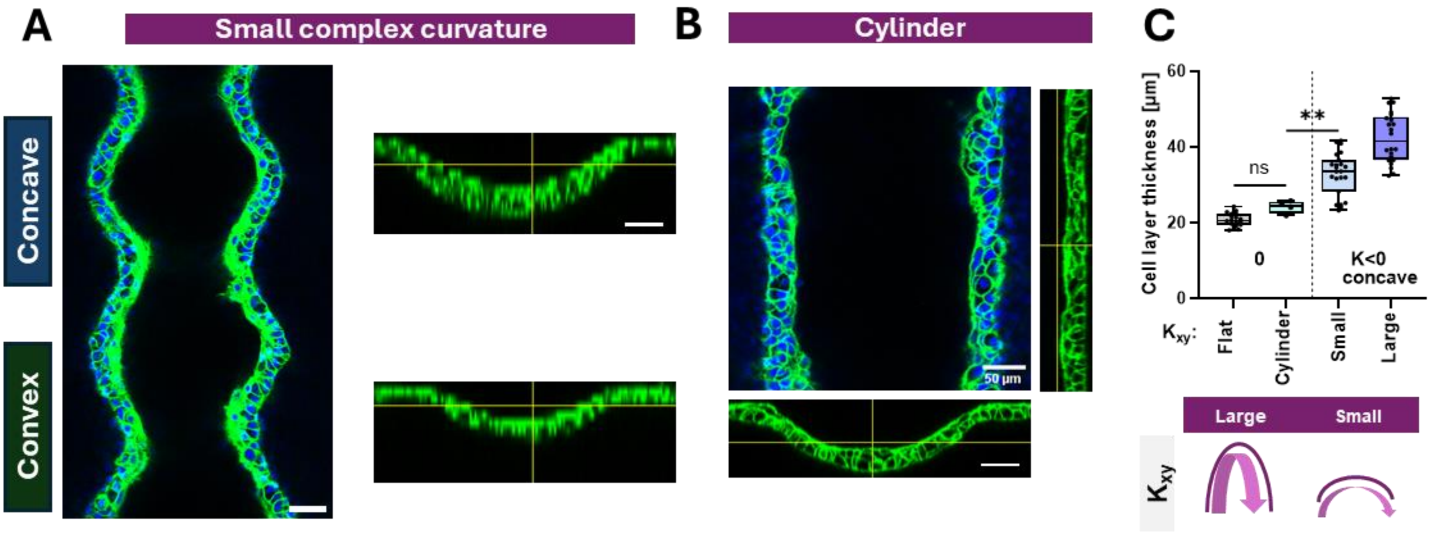


**Figure S34: Comparison of concave gaussian curvature and cylindrical curvature (K=0) with only one concave axis influencing MCF10a cells.** (A) Small complex curvature with smaller Amplitude fabricated with Mid DoS + HepMA. MCF10a cells were cultured for twelve days on the construct. (B) Cylindrical shape of the same material occupied by MCF10a cells (C) Quantification of cell layer thickness for flat, cylindrical and complex curvatures with two concave axis. n=4 w>2.

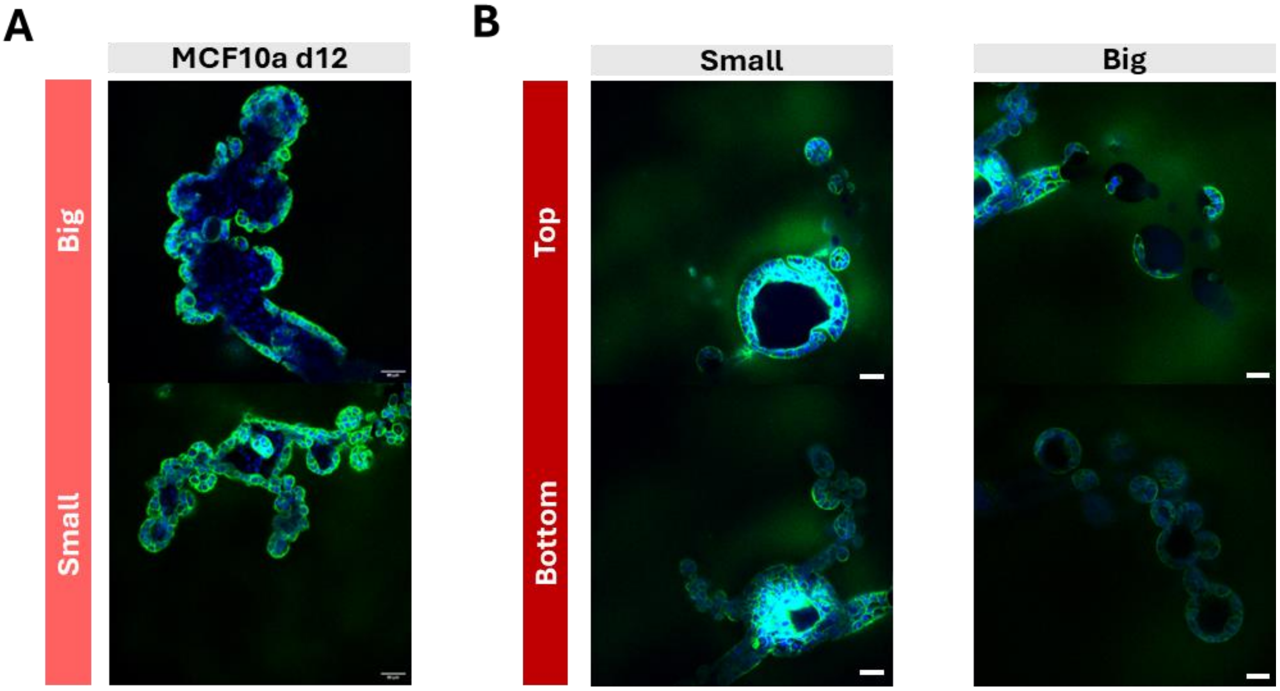


**Figure S35: MCF10a cells occupying alveoli + duct like shapes fabricated with MPA.** (A) MCF10a cells after twelve days of culture in mid DoS GelMA. Top row: big gland, bottom row: smaller glands. (B) MCF10a cells in Mid DoS + HepMA gels with cross-section on the top and bottom of the ablated small glands (C) and big glands. n=3

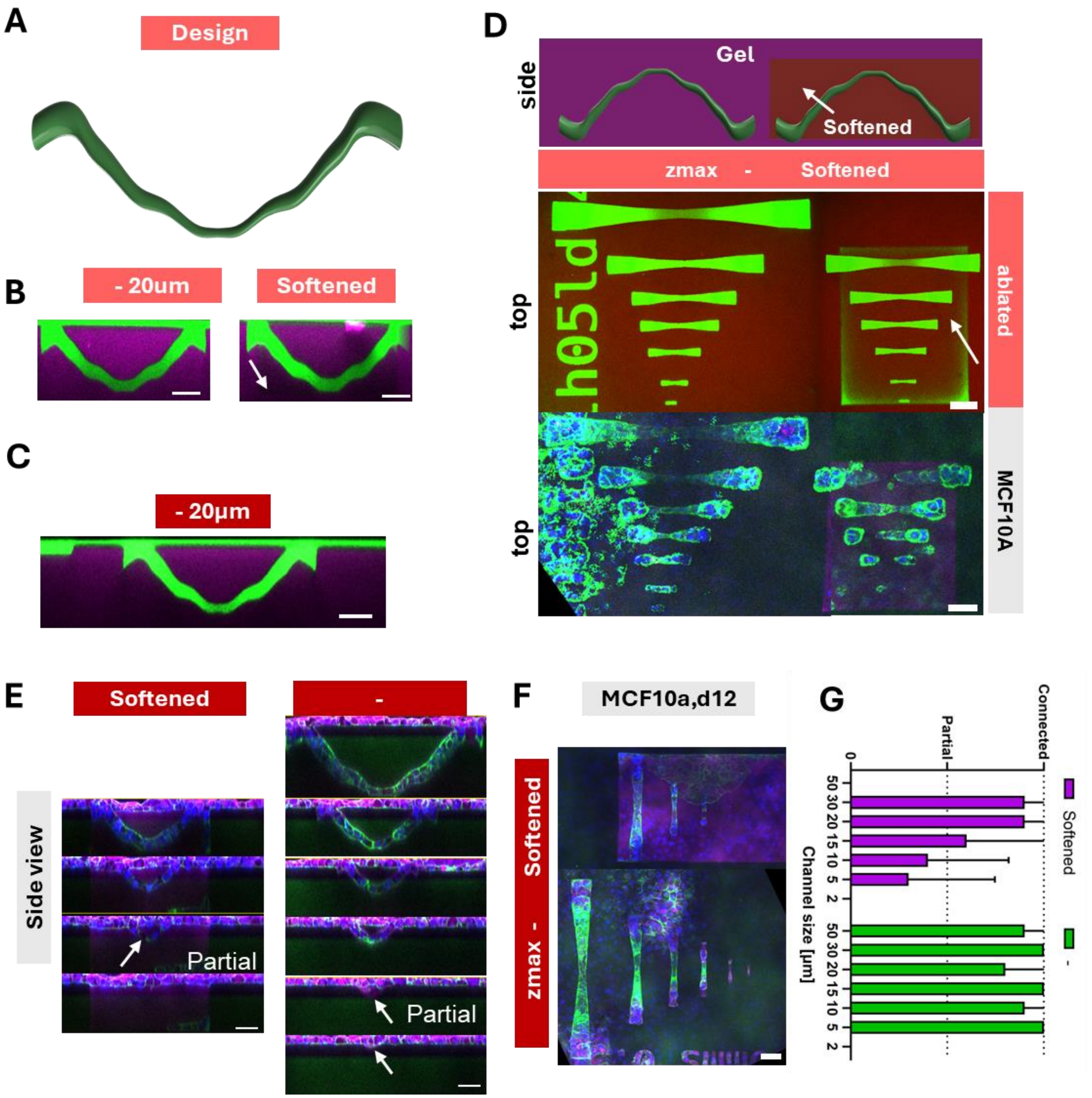


**Figure S36: Channel like geometries fabricated with MPA and cell infiltration into the cavities by MCF10a cells.** (A) Design of connected channels. (B) Ablated channels in Mid DoS GelMA and (C) and Mid DoS + HepMA (D) Z projection of ablated channels in Mid DoS GelMA (overview). Design (top), ablated channels(mid) and MCF10a cells grown into the channels. Softened matrix around ablated channels using a reduced Laser power softened box is indicated with a white arrow. (E) Side view of MCF10a cells grown into softened and non softened channels in stiff Mid DoS + HepMA hydrogels. (H) Z-projection (max intensity) for both conditions (I) Quantification of ingrowth in softened vs non softened channels. n=3, w=2

**ST1: Synthesis of GelMA with different degree' s of Substitution**

| | Gelatin [g] | MAA:Gelatin [ml/g] | MAA tot [ml] | MAA per addition [ml] | CB buffer [M] | Targeted DoF [%] |
|---|---|---|---|---|---|---|
| **Low DoS** | 20 | 0.025 | 0.5 | 0.1 | 0.5 | 45 |
| **Mid DoS** | 20 | 0.05 | 1 | 0.2 | 0.5 | 90 |
| **High DoS** | 20 | 0.4 | 4 | 0.8 | 0.5 | 99 |

**ST2: Ablation parameters for different materials**

| Material | Design | Layer height [µm] | Line distance [µm] | Scanning Speed mm/S | Laser Power [mW] | Comment |
|---|---|---|---|---|---|---|
| **Mid Dos GelMA** | 3 armed duct + alveoli | 1 | 1 | 50 | 375 | |
| | Duct+alveoli different sizes | 1 | 1 | 375 | 450 | |
| | Channels | 1 | 1 | 375 | 250 | |
| | Roughness | 0.5 | 0.5 | 200 | 200 | |
| | Roughness (high throughput) | 4 | 4 | 400 | 100 | 2mM P2CK |
| | Surface softening | 0.5 | 1 | 175 | 250 | |
| **+HepMA** | Duct+alveoli different sizes | 0.5 | 1 | 400 | 100 | **2mM P2CK** |
| | Channels (softening) | 0.5 | 0.5 | 200 | 400 (75mW) | **2mM P2CK** |
| | Roughness | 0.5 | 1 | 400 | 100 | **2mM P2CK** |
| **dECM+HepMA** | Duct+alveoli | 0.4 | 1 | 150 | 100 | |
| | 3 armed duct + alveoli | 0.4 | 1 | 150 | 200 | |
| | Channels | 0.4 | 1 | 150 | 200 | |
| | Half spheres | 0.4 | 1 | 150 | 200 | |
| **HAMA+dECM** | 3 armed duct + alveoli | 0.4 | 1 | 150 | 200 | |
| | Channels | 0.4 | 1 | 150 | 200 | |
| | Duct+Alveoli | 0.4 | 1 | 150 | 200 | |
| **HAMA** | Half spheres | 0.4 | 1 | 400 | 400 | |

### ST3: Fluorophore and Antibody table

| Material | Manufacturer | Catalog Nr | Host | Dilution |
|---|---|---|---|---|
| **Fluorophores** | | | | |
| Fluoreszeinisothiocyanat–Dextran (FITC-Dextran) 2M Da | Sigma Aldrich | FD2000S | - | 1mg/ml |
| Acryloxyethyl thiocarbamoyl rhodamine B | Sigma Aldrich | 908665 | - | **Stock:** 10mg/ml<br>**Resin:** 1ul/ml |
| **Primary antibodies** | | | | |
| Vinculin | Proteintech | 66305-1-Ig | Mouse | 1:100 |
| **Secondary antibodies + dyes** | | | | |
| Hoechst 33342 | Invitrogen | H1399 | - | 1:100 |
| Phalloidin-iFluor 555 Reagent | abcam | ab176756 | - | 1:500 |
| Goat anti-Mouse IgG1 Cross-Adsorbed Secondary ab Alexa Fluor™ 647 | Invitrogen | A-21240 | - | 1:500 |
| Goat anti-Rat IgG (H+L) Cross-Adsorbed Secondary Antibody, Alexa Fluor™ 488 | Invitrogen | A-11006 | - | 1:500 |

FITC-Dextran was diluted in 1x PBS, Rhod B in DMSO, all antibodies+dyes in 1x PBS + 1wt% BSA + 0.01% Triton X

*References*

1 Li, Zhiquan, et al. "Initiation efficiency and cytotoxicity of novel water-soluble two-photon photoinitiators for direct 3D microfabrication of hydrogels." *RSC advances* 3.36 (2013): 15939-15946.

2 Yang, Wei, et al. "Carboxylate modified benzylidene cyclopentanone dyes for one-and two-photon excited photodynamic therapy." *Journal of Photochemistry and Photobiology A: Chemistry* 222.1 (2011): 228-235.
